# Sub-optimal temperature leads to tighter coupling between photosynthetic electron transport and CO_2_ assimilation under fluctuating light in maize

**DOI:** 10.1101/2025.04.12.648513

**Authors:** Cristina R. G. Sales, Stéphanie Arrivault, Tomás Tonetti, Vittoria Clapero, Richard L. Vath, Lucía Arce Cubas, Mark Stitt, Johannes Kromdijk

## Abstract

The C_4_ carbon concentrating pathway promotes high CO_2_ assimilation rates. To keep C_4_ photosynthesis energetically efficient, electron transport reactions and downstream biochemistry need to be carefully balanced. Here we use a combination of non-invasive measurements and metabolic profiling to study the efficiency of C_4_ photosynthesis in maize under two conditions that can lead to decoupling between electron transport and carbon assimilation: fluctuating light and suboptimal temperature. Measurements were performed for three fluctuating light regimes and three temperatures, providing the most detailed study to date of the interaction between fluctuating light and suboptimal temperature on the photosynthetic performance of maize, an important global crop. At room temperature, CO_2_ assimilation rates were decoupled from photosynthetic electron transport under fluctuating light regimes, in contrast to tight coordination observed under constant light. This decoupling was underpinned by metabolic flexibility and buffering by large pools of C_4_ transfer metabolites. Surprisingly, at sub-optimal temperatures, CO_2_ assimilation rates became more tightly coupled to photosynthetic electron transport rates under fluctuating light regimes. This appeared to be caused by strong feedback downregulation of electron transport and a stronger degree of light-saturation of CO_2_ assimilation at low temperature. Low temperature impacted carbon assimilation rates more strongly than metabolite pools or intercellular metabolite distribution, which could reflect negative effects on diffusional metabolite transfer through plasmodesmata. Altogether, these results show that maize is able to maintain energetic efficiency by buffering light transitions under room temperature, as well as avoid oxidative damage by strongly downregulating electron transfer under short-term exposure to low temperature.

**One-sentence summary:** Analysis of maize CO_2_ assimilation under fluctuating light shows significant decoupling from photosynthetic electron transport at room temperature, supported by metabolic flexibility and buffering by large pools of C_4_ transfer metabolites, but tight coordination is restored under suboptimal temperature due to enhanced feedback regulation of electron transport and a stronger degree of light saturation of CO_2_ assimilation.

## Introduction

The C_4_ crop *Zea mays* (maize) was domesticated by ancient farmers in Mexico approximately 9000 years ago (Matsuoka et al. 2002), and its cultivation has since expanded dramatically, currently being the most widely produced food crop in the world (FAO 2023). C_4_ photosynthesis is an adaptation for hot and dry environments and generally does best under high-light conditions (Sage et al. 1999, Sage et al. 2012). Initial carbon acquisition and subsequent assimilation are spatially separated within the leaves of C_4_ species. In species with ‘Kranz’ anatomy, this involves two different photosynthetic cell types, mesophyll cells (MC), and bundle sheath cells (BSC), respectively. The initial carbon fixation takes place in MC via carboxylation of phosphoenolpyruvate (PEP). The resulting C_4_ acids diffuse into BSC where they are decarboxylated, elevating the concentration of CO_2_ around ribulose bisphosphate carboxylase oxygenase (Rubisco), which is exclusively expressed in the chloroplasts of BSC. Reduced 3-carbon metabolites diffuse back to MC, and PEP is regenerated at the expense of ATP. The 10- to 100-fold elevation in CO_2_ concentration around Rubisco in BSC of C_4_ plants compared to MC of C_3_ plants (Furbank and Hatch 1987) strongly suppresses Ribulose-1,5-bisphosphate (RuBP) oxygenation and associated flux through the photorespiration pathway. However, as a result of the additional energetic expense of the carbon concentration mechanism (CCM), C_4_ plants are typically more strongly light-limited across a larger range of light intensities than C_3_ plants (Sales et al. 2021). To maintain light use efficiency, coordination is required between the thylakoid provision of ATP and NADPH, and demand by C_4_ cycle and the Calvin-Benson-Bassham (CBB) cycle reactions across both cell types. In non-stressed maize plants, tight coupling is typically evident from significant linear correlations between the quantum yield of photosystem II (ΦPSII) and the net rate of CO□ assimilation (*A*_CO2_) under a wide range of measurement conditions (e.g., Genty et al., 1989).

Despite the tight coupling observed under non-stressed steady-state conditions, there are two environmental scenarios where light-dependent reactions and *A*_CO2_ have been observed to decouple, i.e., deviate from the common linear correlation. Firstly, fluctuating light (FL) can create a temporal mismatch between the thylakoid reactions and downstream biochemistry (reviewed by Kaiser et al. 2015). In C_3_ species, stomatal opening, as well as activity of CBB enzymes such as Rubisco, chloroplastic fructose-1,6-bisphosphatase (FBPase) and sedoheptulose-1,7-bisphosphatase (SBPase), lag behind rapid increases in incident light, leading to impaired assimilation rates compared to those observed under steady-state light. Rapid decreases in light intensity also negatively impact *A*_CO2_ due to the post-illumination respiratory burst (Prinsley et al. 1986) and slow relaxation of photoprotective energy quenching (Zhu et al. 2004, Kromdijk et al. 2016). In C_4_ species, it has been proposed that the substantial metabolite pools required for transfer between cell types (Leegood 1985, Stitt and Heldt 1985, Arrivault et al. 2017, Medeiros et al. 2022) may be used as a buffer for ATP and reducing equivalents to supplement the provision from the thylakoid reactions during light fluctuations (Stitt and Zhu 2014a), but up to now this theoretical idea has not been experimentally tested. In addition, the reversible reactions involving 3-phosphoglycerate (3PGA) and triose phosphates (TP) have been suggested to provide an intercellular buffering system to balance ATP and NADPH demands between MC and BSC (Leegood and von Caemmerer 1988). Both mechanisms may underpin the observed supra-steady-state *A*_CO2_ following high to low-light transitions observed in C_4_ species across three phylogenetically controlled comparisons by Arce Cubas et al. (2023a). However, metabolite sampling during fluctuating light treatments would be required to verify if the observed stimulation stems from the hypothesized role of metabolites as a capacitor in C_4_ plants during high-to-low-light transitions. In addition, C_4_ photosynthesis may be less efficient during activation from dark-adaptation (Arce Cubas et al. 2023b), or when exposed to rapid increases in light intensity, which may disrupt coordination between the CCM and downstream CBB-cycle carbon assimilation leading to incomplete suppression of photorespiration (Kromdijk et al. 2010, Medeiros et al. 2022) or increases in bundle sheath leakiness, i.e., concentrated CO_2_ which subsequently retro-diffuses from BSC to MC (Sage and McKown 2006, Li et al. 2021, Kubásek et al. 2013, Kromdijk et al. 2014, Wang et al. 2022). Both photorespiration and leakiness represent energetic inefficiencies (von Caemmerer and Furbank 1999, Kromdijk et al. 2014) and therefore alter the stoichiometry between photosynthetic electron transfer and CO_2_ assimilation.

Secondly, coupling between photosynthetic electron transport and *A*_CO2_ is also altered by exposure to sub-optimal temperature. Even though C_4_ photosynthesis should theoretically provide an advantage under cool conditions, and some C_4_ grasses have adapted to cool climates (Du et al. 1999, Long 1999, Long and Spence 2013), C_4_ plants are often particularly sensitive to suboptimal temperature. Temperatures below 15°C are low enough to cause chilling stress in maize (Hu et al. 2017, Frascaroli and Revilla 2019, Burnett and Kromdijk 2022) and in combination with exposure to light may give rise to chilling-induced photoinhibition (Taylor and Craig 1971, Long 1983). Maize leaves that develop under low temperature tend to show strong decoupling between *A*_CO2_ and linear electron flow. More specifically, strongly increased ratios between electron transport and net CO_2_ assimilation compared to those found in unstressed leaves are typically observed in leaves kept at suboptimal temperature (e.g., Fryer et al. 1998). This may suggest that exposure to suboptimal temperature enhances energy flow to alternative electron sinks to mitigate the decline in carbon assimilation, but the specific mechanisms behind this phenomenon remain unclear. The water-water cycle, whereby linear electron flow from PSII to photosystem I (PSI) is sustained by reduction of O_2_ and subsequent formation of water, may be elevated in C_4_ grasses (Siebke et al. 2003) and has been suggested to increase under low temperature in both C_3_ and C_4_ species to provide protection against photoinhibition (Ort and Baker, 2002). However, experimental verification could not confirm significant flux through the water-water cycle (Driever et al. 2011). In addition, it is not clear if increases in electron flow per fixed CO_2_ occur in response to short-term low temperature exposure (hours) and if so, whether these get more pronounced under sharp fluctuations in light intensity, in which case the alternative electron sink may work like a safety valve, by helping to avoid over-reduction of electron carriers in the thylakoid membrane.

Tight coordination between photochemical supply of NADPH and ATP, C_4_ shuttle and CBB cycle carbon assimilation are important features for efficient carbon fixation in C_4_ species (Kromdijk et al. 2014, von Caemmerer 2000), and studying conditions in which this coordination breaks down can help identify prerequisites for efficient C_4_ photosynthesis. Analysis of metabolite profiles provides a means to assess bottlenecks and determine molecular mechanisms underpinning decoupling of CO_2_ assimilation and electron flow. In C_3_ plants the ratio between 3PGA and dihydroxyacetone phosphate (DHAP) provides a simple proxy for the provision of ATP and reductant to drive phosphorylation and reduction in the CBB cycle (Dietz and Heber 1986). For C_4_ plants, early work by Labate et al. (1990) showed that this ratio decreases with temperature in maize down to 12°C, but then increased with further decreases in temperature (to 8°C), which may suggest a restriction in electron transport becomes dominant somewhere between 8 – 12°C, possibly due to photosynthetic control at the cytochrome *b*_6_*f* complex. However, since 3PGA and DHAP form opposing diffusional gradients between the BSC and MC and 3PGA equilibrates with PEP in the MC, changes in the gradients of 3PGA and DHAP at low temperature may complicate this interpretation in C_4_ plants. Indeed, Labate et al. (1990) observed decreases in whole leaf 3PGA and DHAP pools, which were hypothesized to reflect a decline in intercellular diffusion between BSC and MC, but experimental verification of MC and BSC specific pools at sub-optimal temperatures is still lacking.

While current literature shows that both fluctuating light and suboptimal temperature can lead to decoupling between photosynthetic electron transport and *A*_CO2_, to our knowledge there is currently no published data on their combined effects, nor on the metabolic changes that may accompany these conditions and potentially explain non steady-state patterns of *A*_CO2_. In this work we therefore aimed to address these knowledge gaps by studying maize CO_2_ fixation as a function of three fluctuating light treatments with differing step lengths (6, 30 and 300 s) at ambient and two sub-optimal temperatures (25, 15 and 7°C). We specifically hypothesized that:

i) decoupling between electron transport and carbon fixation will be more pronounced under light fluctuations with shorter light step lengths;
ii) after a transition from high to low-light, CO_2_ fixation will be transiently supported by energy and reductant from metabolite pools;
iii) immediately after a transition from low to high-light, slow build-up of metabolite pools will cause a lag in CO_2_ fixation;
iv) short-term exposure to suboptimal temperatures will exacerbate decoupling between electron transport and carbon fixation in line with longer-term acclimatory responses (e.g. Fryer et al. 1998);
v) Suboptimal temperature will decrease intercellular metabolite diffusion between BSC and MC as hypothesized by Labate *et al*. (1990).

To address these hypotheses, we non-steady state leaf gas exchange measurements in conjunction with chlorophyll fluorescence and near-infra-red differential absorption to monitor electron transfer efficiencies across the two photosystems. In addition, rapid freeze-sampling during gas exchange measurements (Xu et al. 2021) was used to capture metabolite profiles at key time-points and leaf-fractionation by serial filtration over liquid nitrogen was used to resolve distribution of key metabolite pools between M and BSC. Our results reflect substantial metabolic flexibility at 25°C, but surprisingly showed that coupling between electron transport and *A*_CO2_ was progressively enhanced under low temperature, which we suggest originates both from strong regulation of electron transport to minimize oxidative stress, as well as the strong decrease in CO_2_ assimilation flux under low temperature, which decreases the rebalancing requirements in response to changes in incident irradiance.

## Results

### Low temperature dampens the effect of fluctuating light on CO_2_ assimilation

Photosynthetic CO_2_ assimilation was first measured under steady-state conditions in maize plants kept at 25, 15, or 7°C for 2h. Light response curves were performed (Fig. 1A). Parameters derived from the curve fitting are shown in Supplemental Table S1. Compared to 25°C, light saturated CO_2_ assimilation (*A*_sat_) was reduced by 52%, and 86% at 15°C and 7°C, respectively; mitochondrial respiration (*R*_d_) decreased approximately 46% of the rate at 25°C in plants measured at 15°C, and the reduction was ∼93% in plants measured at 7°C. At 15°C, the maximum quantum yield of CO_2_ assimilation (□CO_2,_ (μmol CO_2_ mol^−1^ photons) did not differ significantly from 25°C (0.062 vs 0.068, respectively) but decreased by 59% when measured at 7°C (0.039). Finally, the saturation of A_CO2_ with light was markedly affected by sub-optimal temperature, reaching 95% of the rate at 2000 µmol m^−2^ s^−1^ PFD at approximately 1800 µmol m^−2^ s^−1^ for measurements at 25°C, but at much lower intensities of ∼1000 and 400 µmol m^−2^ s^−1^ for measurements at 15 and 7°C respectively. CO_2_ response curves were used to further assess the biochemical limitations of *A*_CO2_ to changes in temperature (Fig. 1B). The maximum *in vivo* carboxylation rate of phosphoenolpyruvate carboxylase (*V*_pmax_) was reduced by 54% and 84% at 15°C and 7°C, respectively, of the rate at 25°C. Finally, *V*_max_, the CO_2_-saturated rate of photosynthesis, was reduced by 56% at 15°C, and by 80% at 7°C (Supplemental Table S1).

**Figure 1.**
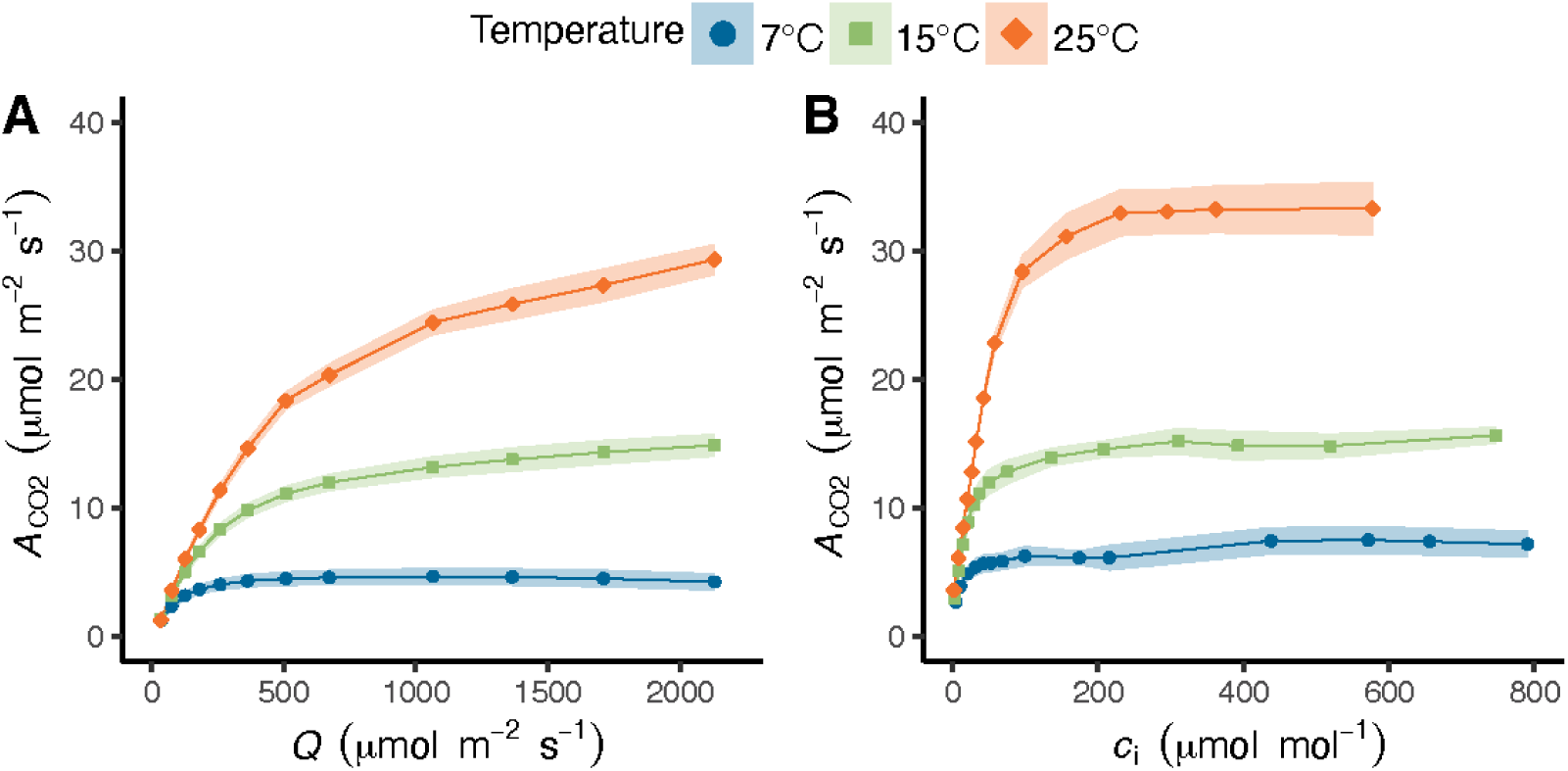
Response curves of net CO_2_ assimilation (*A*_CO2_) in maize leaves measured at different temperatures. A) Net CO_2_ assimilation as a function of photosynthetically active radiation (Q), and (B) net CO_2_ assimilation as a function of intercellular CO_2_ concentration (*c*_i_). Measurements were performed on maize plants acclimated at 7, 15 or 25°C for at least 2 h. Ribbons represent standard error of the mean (*n* = 4-5 biological replicates).

To compare how these temperature effects play out under fluctuating light conditions, we measured short-term photosynthetic responses to three distinct FL treatments, under the same three measurement temperatures. The FL regimes consisted of repetitively switching between high-light and low-light for one hour. Photosynthetically active radiation was 1500 µmol m^−2^ s^−1^ in the high-light phase, and 200 µmol m^−2^ s^−1^ in the low-light phase. Each light step lasted either 6, 30, or 300 s, called from now on as 6sFL, 30sFL, and 300sFL (last 10 minutes shown in Fig. 2, full hour in Fig S1). As expected, the *A*_CO2_ response showed increases in the 1500 µmol m^−2^ s^−1^ phase, followed by a decline when light switched to 200 µmol m^−2^ s^−1^. At 25°C, the difference between *A*_CO2_ values at 1500 and 200 µmol m^−2^ s^−1^ progressively increased with increasing light step length (Fig. 2 A-C). In addition, following the transition from low to high-light in the 300sFL, *A*_CO2_ showed a biphasic increase with a shoulder around 30 s. When *A*_CO2_ was measured under FL at lower temperatures, the effect of light step length on the difference between *A*_CO2_ values at the two light levels decreased for 15°C (Fig 2 D-F) and was completely lost at 7°C (Fig. 2 G-I). The shoulder at 30 s during the 300sFL 1500 µmol m^− 2^ s^−1^ phase was also lost at lower temperatures (Fig. 2 F and I).

**Figure 2.**
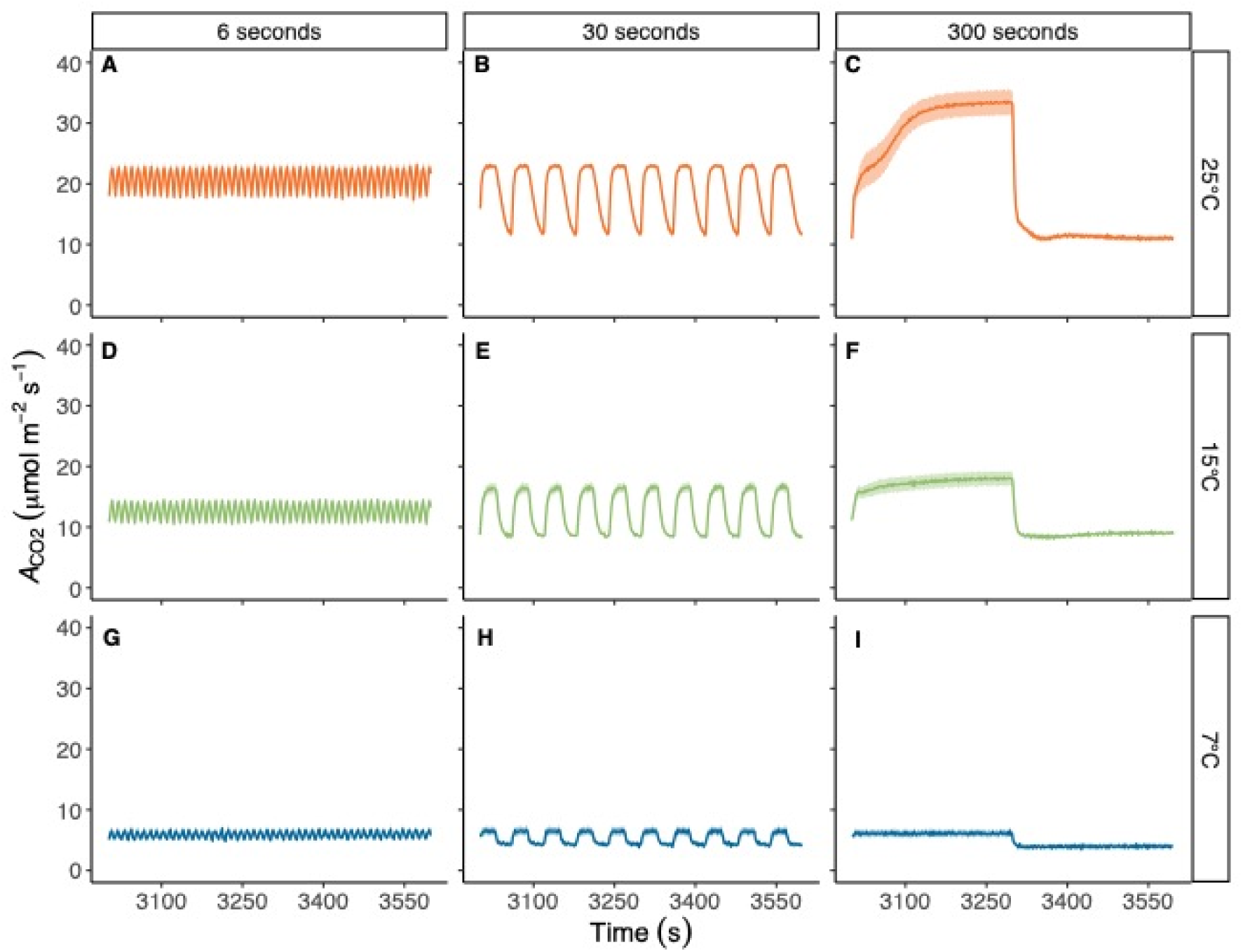
Net CO_2_ assimilation (*A*_CO2_) in maize plants measured under three different fluctuating light regimes, at 25 °C (A, B, and C), 15 °C (D, E, and F), or 7 °C (G, H, and I). In each fluctuating light regime, leaves were exposed to repetitive changes between low (200 µmol m^−2^ s^−1^) and high (1500 µmol m^−2^ s^−1^) light-steps with duration of either 6 s (FL6; A, D, G), 30 s (FL30; B,E, H) or 300 s (FL300; C, F, I). Measurements were performed on maize plants acclimated at 7, 15 or 25°C for at least 2 h. Fluctuating light regimes were started after leaves were acclimated to steady state at light intensity of 600 µmol m^−2^ s^−1^ and lasted 1 hour. Data are shown from the final ten minutes of each light regime. Ribbons represent standard error of the mean (*n*=4-5). Full timeseries data are provided in Supplemental Fig S1 and timing of the changes in light intensity across each FL regime are provided in Supplemental Fig S2.

Average values of *A*_CO2_ at high or low-light were obtained for the last 10 min cycle for each FL treatment by temperature combination (Fig. 3). To facilitate comparisons, values of *A*_CO2_ under steady-state taken from the light response curves (Fig. 1A) at 1500 and 200 µmol m^−2^ s^−1^ are also shown. At 25°C, the more rapid FL treatments (6sFL and 30sFL) caused a significant reduction of ∼30% in *A*_CO2_ at high-light (Fig. 3A and B) compared to 300sFL and steady-state conditions (Fig. 3C and D). On the other hand, at low-light conditions, the more rapid FL treatments gave rise to the highest integrated *A*_CO2_, with 6sFL showing a significant ∼238% increase in *A*_CO2_ (Fig. 3E) compared to steady-state conditions (Fig. 3H). Thus, at 25°C, higher fluctuation frequency reduced the amplitude of A_CO2_ changes between light intensities, with A_CO2_ staying intermediate between photosynthetic rates reached in both light conditions at lower fluctuation frequencies. At 15°C, these patterns were less pronounced, with the most rapid FL treatment (6sFL) causing a reduction of ∼20% in *A*_CO2_ at 15°C (Fig. 3A) compared to 300sFL and steady-state conditions (Fig. 3C and D) at high-light and a ∼185% increase in *A*_CO2_ (Fig. 3E) compared to steady-state conditions (Fig. 3H) at low-light. At 7°C, the effects of light fluctuations on *A*_CO2_ were fully suppressed and none of the FL treatments led to significantly different *A*_CO2_ compared to steady-state conditions.

**Figure 3.**
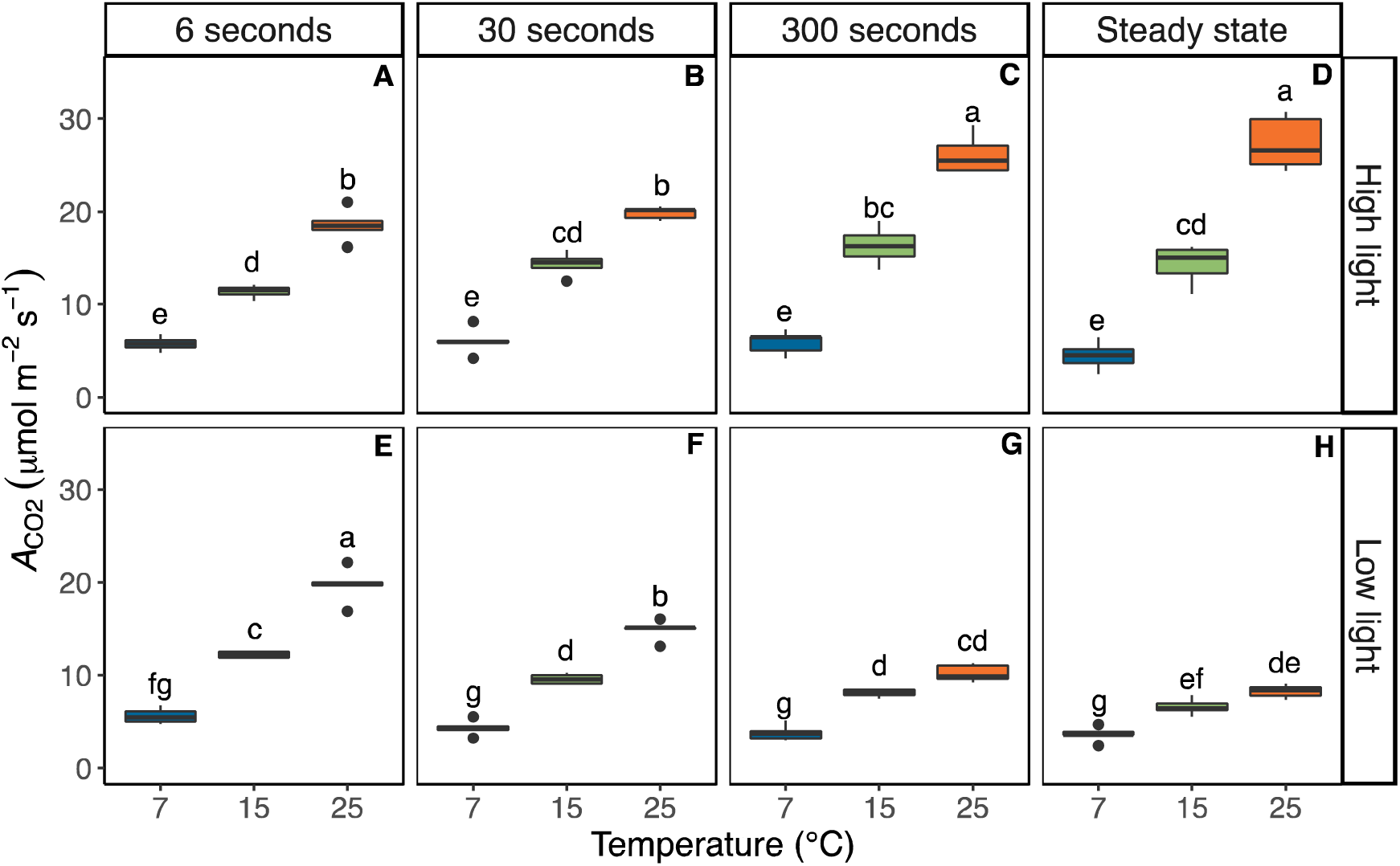
Integrated values of net CO_2_ assimilation (*A*_CO2_) as a function of temperature and fluctuating light regime. Measurements were performed at 25 °C, 15 °C, or 7 °C and three fluctuating light regimes. Measurements were performed on maize plants acclimated at 7, 15 or 25°C for at least 2 h. In each fluctuating light regime, leaves were exposed to repetitive changes between low (200 µmol m^− 2^ s^−1^) and high (1500 µmol m^−2^ s^−1^) light-steps with duration of either 6, 30 or 300 s (FL6, FL30, FL300). Integrated assimilation rates were calculated from the area under the curve (AUC) of *A*_CO2_ for each period of low light (A-C) or high light (E-G) illumination during the last 10 minutes of each fluctuating light treatment. Numbers were converted to a rate for ease of comparison. Measurements at steady state at matching light intensities and temperatures were obtained from light response curves shown in Fig 1A and included for comparison (D, H). Box edges represent the lower and upper quartiles, the solid line indicates the median, and points represent outliers beyond 1.5 times the interquartile range (*n* = 4-5 biological replicates). Statistical analyses were run on square root transformed data. Two-way ANOVA was used to test the effect of temperature and fluctuation length. Different letters indicate statistical differences according to Tukey test (*p*<0.05).

### Low temperature dampens fluctuations in Φ_PSII_ and Φ_PSI_ and enhances coupling between linear electron transport and CO_2_ assimilation

Combined chlorophyll fluorescence and near-infrared differential absorption measurements were used concurrently with the gas exchange measurements in Fig. 1 and 2 to estimate PSII and PSI operating efficiency, both during the steady-state light response curves as well as at selected time-points (3 s after the start and 3 s before the end of the high light, and 3 s after the start and 3 s before the end of the low light phase); see scheme of measurement timing in Fig. S2).

Similar to the *A*_CO2_ measurements, Φ_PSII_ responses to the light fluctuation were pronounced at 25 °C, but strongly dampened at 15°C and Φ_PSII_ at 7°C was similar across all measurements performed within each light intensity. Φ_PSII_ measurements were most variable in the 300sFL treatment, whereas under more rapid FL regimes (6sFL and 30sFL), no significant differences were observed in Φ_PSII_ between the start and the end of each light step (Fig. 4A-B and E-F). At 25 °C, Φ_PSII_ increased significantly from the start to the end of the high light step in the 300sFL regime (Fig. 4C). Just after transitioning from low-light to high-light (labelled ‘Start’), Φ_PSII_ showed a ∼33% decrease compared to steady-state (Fig. 4C vs 4D). In contrast, the measurement at the end of the high-light step (labelled ‘End’), showed significantly higher Φ_PSII_ compared to steady-state (Fig. 4G vs 4H). Φ_PSII_ can be impacted by the redox state of Q_A_, which determines PSII electron acceptor-side limitation and by the level of NPQ which impacts PSII donor-side limitation. At 25 °C, Q_A_ was highly reduced at the ‘start’ timepoint, but less so at the ‘end’ timepoint of the 300sFL high light step (Fig S3C), which was also significantly lower than steady state (Fig S3D). In contrast, NPQ was significantly lower at the ‘start’ timepoint than steady state, but increased at the ‘end’ timepoint, which was similar to steady state (Fig S4C vs D). Thus, at 25 °C with increasing duration of high light exposure, a shift from PSII acceptor-side to donor-side limitation was observed. This pattern was similar but dampened under 15 °C, whereas under 7 °C, Q_A_ stayed highly reduced (Fig S4A-C) and NPQ stayed high and invariable at all timepoints and FL regimes (S4A-C).

**Figure 4.**
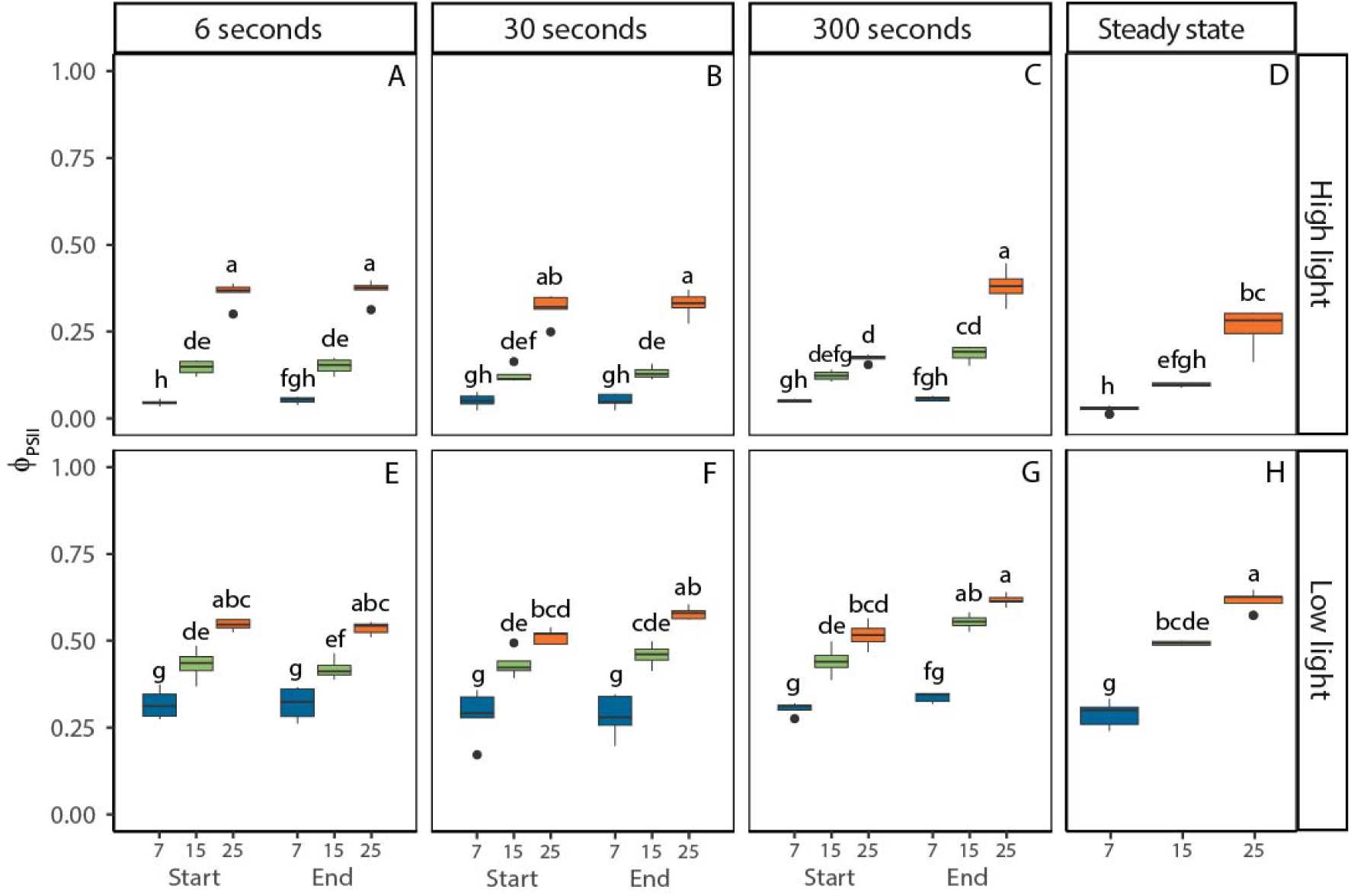
Photosystem II quantum yield (ϕ_PSII_) in maize leaves as a function of temperature and fluctuating light regime. Measurements were performed at 25 °C, 15 °C, or 7 °C and three fluctuating light regimes. Maize plants were acclimated at 7, 15 or 25°C for at least 2 h prior to measurements. In each fluctuating light regime, leaves were exposed to repetitive changes between low (200 µmol m^−2^ s^− 1^) and high (1500 µmol m^−2^ s^−1^) light-steps with duration of either 6, 30 or 300 s (FL6, FL30, FL300). In each fluctuating light regime, ϕ_PSII_ was determined 3 s into a new light intensity (‘Start’) and 3 s before switching (‘End’). Thus, a total of four measurements were taken for each biological replicate, two measurements (‘Start’ and ‘End’) during high light (A-C) and two measurements during low light (E-G). Precise timings of measurements in each fluctuating light regime are provided in Supplemental Fig. S2. Measurements at steady state at matching light intensities and temperatures were obtained from light response curves shown in Fig 1A and included for comparison (D, H). Box edges represent the lower and upper quartiles, the solid line indicates the median, and points represent outliers beyond 1.5 times the interquartile range (*n* = 4-5 biological replicates). Statistical analyses were run on Box-Cox transformed data. Three-way ANOVA was used to test the effect of temperature, fluctuation length, and measurement time. Different letters indicate statistical differences according to Tukey test (*p*<0.05).

The low light Φ_PSII_ measurements in the 300sFL regime also significantly increased from ‘start’ to ‘end’ at both 15 °C and 25 °C (Fig 4G), with the ‘end’ measurements indistinguishable from steady state (Fig 4G vs H). Acceptor-side limitation by reduced Q_A_ was generally low and invariable across the low light measurements and therefore the increase in Φ_PSII_ from ‘start’ to ‘end’ predominantly reflected the relaxation of donor-side limitation by NPQ, which started high due to the preceding high light period and was significantly lower at the ‘end’ than the ‘start’ timepoint (P<0.05, Fig. S4F and G). At 7 °C low light Φ_PSII_ measurements were significantly lower than measurements at higher temperatures and were invariable between FL regimes, timepoints and steady state (Fig 4E-H). Consistently, the Q_A_ pool remained approximately 50% reduced throughout all low light measurements at 7 °C (Fig S3E-H) and NPQ remained high and became unresponsive to changing light levels in the FL regimes (Fig S4E-H).

Operating efficiency of PSI (Φ_PSI_) followed similar patterns as observed for Φ_PSII_ but showed much smaller changes. At 25°C in the high-light phase, Φ_PSI_ was generally higher than steady-state, significantly so for both measurements at 6sFL treatment (Fig. 5A, P<0.05), as well as the ‘end’ measurement at 300sFL, (Fig. 5C, P<0.05). These differences did not persist at lower temperatures, where Φ_PSI_ was statistically similar across all FL regimes and timepoints. Electron flow at the donor side of PSI showed a restriction at the cytochrome-b_6,f_-complex as evident from near-infrared differential absorption estimates of plastocyanin redox state, which was fully oxidised throughout all high light measurements, regardless of temperature (Fig. S5A-D). In contrast, the redox state of ferredoxin was only fully reduced at 7 °C, but significantly more oxidised at 15 and 25 °C, in line with the increasing demand for NADPH with increasing temperature (Fig. S6A-D). Under low-light, Φ_PSI_ was close to 1 at both 25 and 15°C, regardless of the FL regime. At 7°C, Φ_PSI_ was significantly lower than the values observed at higher temperatures and was generally similar in all FL regimes and steady state (Fig. 5E-H). These patterns could reflect both slower electron flow from plastocyanin to PSI, which was fully oxidized at 7 °C but significantly more reduced at higher temperatures (P<0.05, Fig S5E-H), as well as slower electron flow from PSI towards ferredoxin, which was most reduced at 7 °C, but significantly more oxidized at higher temperatures (P<0.05, Fig S6E-H).

**Figure 5.**
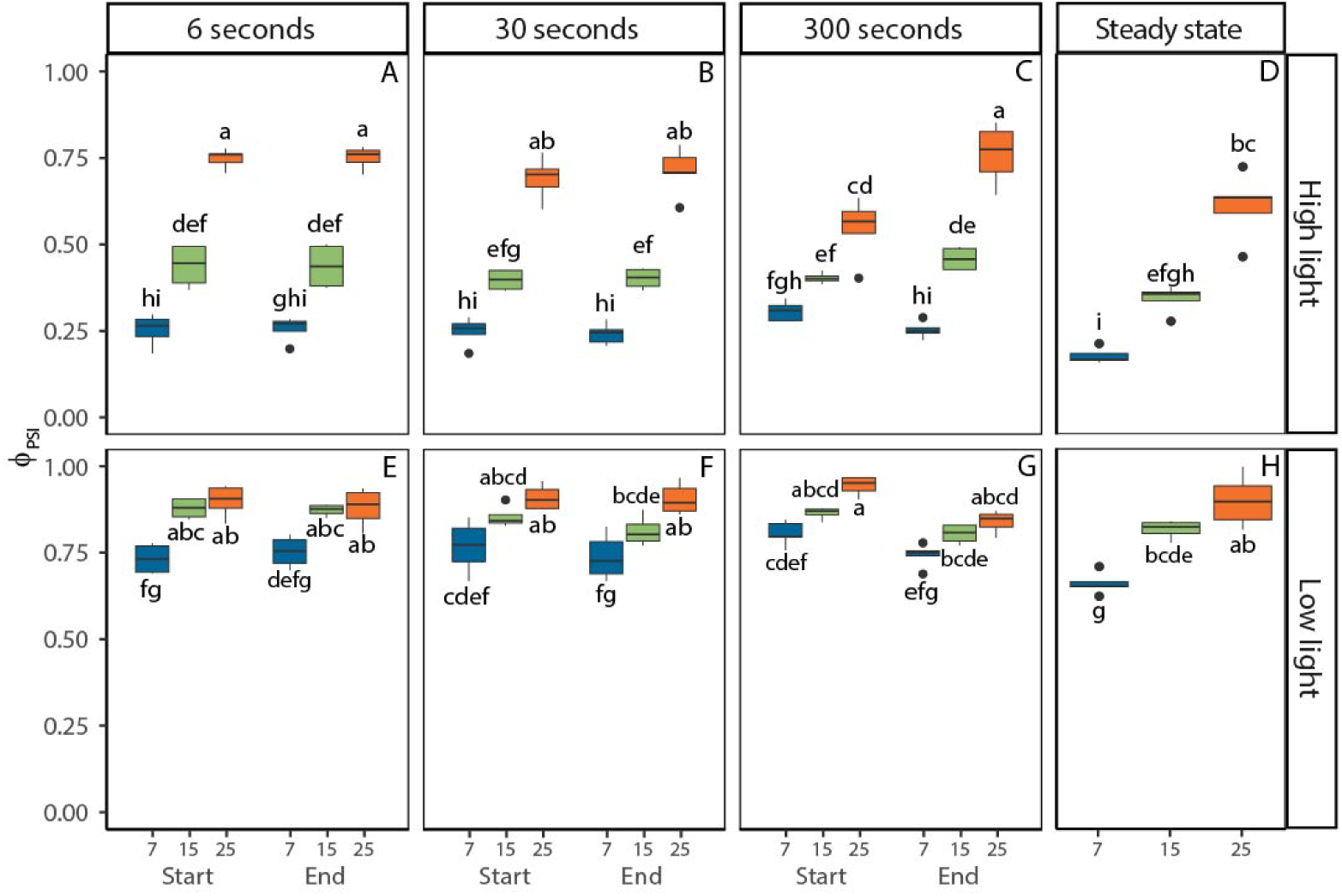
Photosystem I quantum yield (ϕ_PSII_) in maize leaves as a function of temperature and fluctuating light regime. Measurements were performed at 25 °C, 15 °C, or 7 °C and three fluctuating light regimes. Maize plants were acclimated at 7, 15 or 25°C for at least 2 h prior to measurements. In each fluctuating light regime, leaves were exposed to repetitive changes between low (200 µmol m^−2^ s^− 1^) and high (1500 µmol m^−2^ s^−1^) light-steps with duration of either 6, 30 or 300 s (FL6, FL30, FL300). In each fluctuating light regime, ϕ_PSI_ was determined 3 s into a new light intensity (‘Start’) and 3 s before switching (‘End’). Thus, a total of four measurements were taken for each biological replicate, two measurements (‘Start’ and ‘End’) during high light (A-C) and two measurements during low light (E-G). Precise timings of measurements in each fluctuating light regime are provided in Supplemental Fig. S2. Measurements at steady state at matching light intensities and temperatures were obtained from light response curves shown in Fig 1A and included for comparison (D, H). Box edges represent the lower and upper quartiles, the solid line indicates the median, and points represent outliers beyond 1.5 times the interquartile range (*n* = 4-5 biological replicates). Statistical analyses were run on Box-Cox transformed data for high light measurements. Three-way ANOVA was used to test the effect of temperature, fluctuation length, and measurement time. Different letters indicate statistical differences according to Tukey test (*p*<0.05).

The ratio between Φ_PSII_ and Φ_CO2_ calculated from gas exchange at matching timepoints (Fig. S7) provides a measure of the number of electrons transferred per CO_2_ fixed (e^−^ PSII/CO_2_, Fig. 6), i.e., the degree of coupling between whole-chain electron transport and CO_2_ assimilation (Krall and Edwards 1990). Under steady-state high-light conditions, e^−^PSII/CO_2_ was 17 ± 5 and did not differ significantly between different measurement temperatures (Fig. 6D). In contrast, the impact of temperature was pronounced for the high light steps in the FL regimes. At 25 °C, e^−^ PSII/CO_2_ at the ‘start’ timepoint was significantly higher than steady state (P<0.05) for 6sFL (Fig. 6A) and 30sFL (Fig. 6B). Under sub-optimal temperatures, high light e^−^ PSII/CO_2_ was significantly higher than steady state (P<0.05) only in the 6sFL regime at 15 °C (21 ± 3 e^−^ PSII/CO_2_) and was similar to the steady-state ratio across all FL treatments at 7 °C (Fig. 6A-B). At low-light, the steady state e^−^PSII/CO_2_ ratio was 14 ± 2 electrons per CO_2_ fixed and did not vary with temperature (Fig. 6H). In contrast, at the ‘start’ timepoint of the 25 °C measurements, e^−^PSII/CO_2_ was significantly lower than steady state across all FL regimes at an average value of 5 ± 1 e^−^PSII/CO_2_ (Fig. 6E-G, P<0.05). At 15°C, similar but dampened differences were observed, with e^−^PSII/CO_2_ showing average values of 9 ± 3 at the ‘start’ timepoint. At 7 °C no significant differences were observed between timepoints or FL regimes with all measurements showing very similar values at 13 ± 3 e^−^PSII/CO_2_ (Fig. 6E-H). Altogether, these findings show that the tight correlation between electron transport and net CO_2_ fixation is maintained under low temperature and steady state light conditions, but that shortly after a change in light intensity, the electron requirements per CO_2_ fixed can vary dramatically, increasing up to 76% following a change from low to high light and decreasing by 64% following a change from high to low light, both of which could be consistent with metabolic buffering of CO_2_ assimilation. Contrary to our hypothesis, these phenomena were only observed at 25 °C and measurements under suboptimal temperatures showed less variation in e^−^PSII/CO_2_. To find out more about the role of metabolic changes in each of these observations, we next assessed the variation in whole-leaf metabolite profiles.

**Figure 6.**
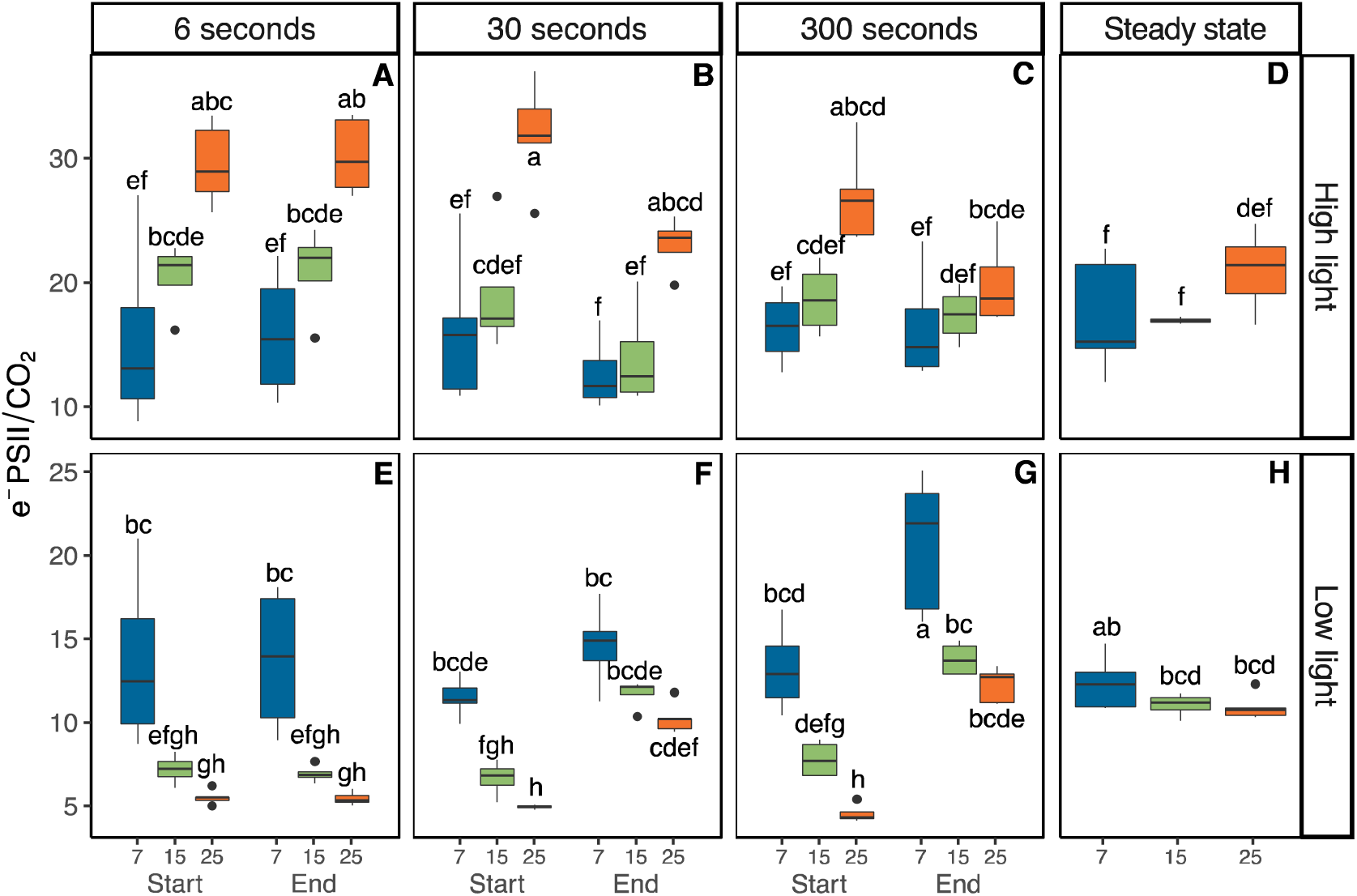
Linear electron transport requirements per fixed CO_2_ (e^−^ PSII/CO_2_) in maize leaves as a function of temperature and fluctuating light regime. Measurements were performed at 25 °C, 15 °C, or 7 °C and three fluctuating light regimes. Maize plants were acclimated at 7, 15 or 25°C for at least 2 h prior to measurements. In each fluctuating light regime, leaves were exposed to repetitive changes between low (200 µmol m^−2^ s^−1^) and high (1500 µmol m^−2^ s^−1^) light-steps with duration of either 6, 30 or 300 s (FL6, FL30, FL300). In each fluctuating light regime, ϕ_PSI_ was determined 3 s into a new light intensity (‘Start’) and 3 s before switching (‘End’). Thus, a total of four measurements were taken for each biological replicate, two measurements (‘Start’ and ‘End’) during high light (A-C) and two measurements during low light (E-G). Precise timings of measurements in each fluctuating light regime are provided in Supplemental Fig. S2. Measurements at steady state at matching light intensities were obtained from light response curves shown in Fig 1A and included for comparison (D, H). Box edges represent the lower and upper quartiles, the solid line indicates the median, and points represent outliers beyond 1.5 times the interquartile range (*n* = 4-5 biological replicates). Statistical analyses were run on square root transformed data for high light measurements, and Box-Cox transformed data for low light measurements. Three-way ANOVA was used to test the effect of temperature, fluctuation length, and measurement time. Different letters indicate statistical differences according to Tukey test (*p*<0.05).

### Metabolite profiles at 25 °C lag behind changes in light intensity

To further address the hypothesised metabolic origins of the strong decoupling between CO_2_ assimilation and linear electron transport observed primarily at 25 °C (Fig 6), we determined levels of 25 metabolites involved in the NADP-ME and PEPCK C_4_ pathways, in the CBB cycle and in the photorespiration pathway, as well as a number of key organic acids, amino acids, and sugars from leaf samples taken at key time-points during the 300sFL regime at 25 °C. Leaves were sampled 10 s prior and following each light switch (i.e. 10 s after light switch from low to high - FL300_10s_HL; 290 s after light switches from low to high light phase - FL300_290s_HL; 310 s after light switches from low to light phase - FL300_310s_LL; and 590 s after light switches from low to high light phase - FL300_590s_LL). Samples were also taken from steady state illumination at 200 µmol m^−2^ s^−1^ (low light, LL_Steady-state), or 1500 µmol m^−2^ s^−1^ (high light, HL_Steady-state).

Nine metabolites varied significantly with sampling time-point: aspartate, pyruvate, PEP, DHAP, S7P, Ru5P+Xu5P, ADPG, glycerate and 2OG (P<0.05, Supplemental Fig. S8). Principle component analysis (PCA) showed clustering of the steady-state LL and HL samples at negative and positive values of PC2, respectively (Fig 7A), with strong positive loadings for pyruvate, DHAP, ADPG, glycerate and S7P and strong negative loadings for aspartate, alanine, malate, FBP and RuBP (Fig 7C). Interestingly, positioning along PC2 of the four sampled time-points showed a clear time lag in response to light. Namely, while samples collected just before the end of the 300 s light step (FL300_290s_HL and FL300_590s_LL) clustered with samples taken at steady-state exposure to the same light level (HL and LL respectively; Fig 7B), samples taken 10 s after the light switch (FL300_10s_HL and FL300_310s_LL) grouped with the steady-state samples of the opposite light level (LL and HL, respectively), in line with the strong decoupling between linear electron transport and CO_2_ assimilation observed at these time points.

**Figure 7.**
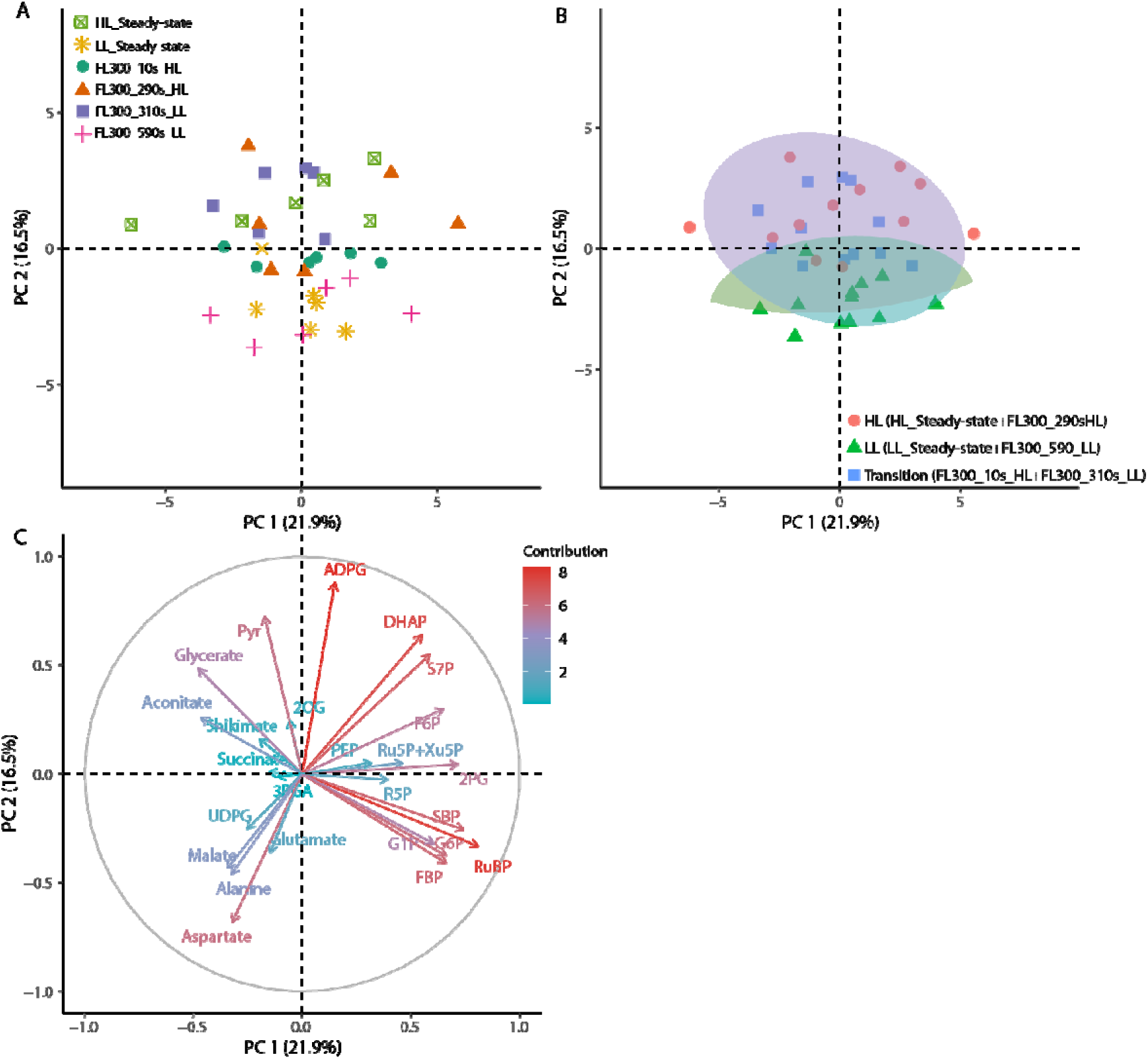
Principal component analysis of metabolite profiles of maize leaves exposed to constant or fluctuating light. For constant light, leaves were sampled following exposure with low (200 µmol m^−2^ s^−1^) and high (1500 µmol m^−2^ s^−1^) light intensity. For the fluctuating light regime, leaves were exposed to repetitive changes between low and high light-steps with duration of 300 s (FL300). Samples were taken at four different time points: 10 s after switching from low to high light (FL300_10s_HL), 290 s after switching from low to high light (FL300_290s_HL), 10 s after switching from high to low light (FL300_310s_LL), and 290 s after switching from high to low light (FL300_590s_LL). Plants were acclimated to 25 °C for at least 2h prior to sampling. Individual sample positions according to the first two PCs are shown in A), with different symbols and colours indicating the corresponding light intensity and sampling timepoints. Panel B shows the same sample positions as in A) grouped in three clusters indicated by different symbols and colours. Panel C shows the different loadings per metabolite. The complete metabolite data are provided in Supplemental Fig.S8

These data were further explored to look for metabolic changes which may explain sub-steady state A_CO2_ at FL300_10s_HL (Fig. S8Z). Interestingly, the levels of PEP sampled at FL300_10s_HL were lower than steady-state levels. This decrease in PEP might restrict PEPC activity, and in conjunction with low levels of DHAP at FL300_10s_HL, a positive regulator of PEPC activity, suggests that shortfalls both in the substrate supply and in activating metabolites are compounding negatively on PEPC activity.

Metabolite profiles at different timepoints were also scrutinized for changes that may underpin the observed supra-steady state A_CO2_ (Fig. S8Z) and decreased e^−^PSII/CO_2_ (Fig. 6E-H) at FL300_310_LL. Significant increases in 3PGA/PEP were found at time point FL300_310s_LL (P<0.05, Fig. S9B). Meanwhile, the level of pyruvate at FL300_310s_LL remained similar to the preceding FL300_290s_HL high-light time-point and the steady-state level at high-light, and was significantly higher than the level at FL300_590s_LL or at steady-state low-light (P<0.05, Fig. S8C). These patterns of pyruvate and 3PGA/PEP indicate that C_4_ cycle activity remained higher than under steady-state low-light in the FL300_310s_LL, consistent with significant decoupling between *A_CO2_* and linear electron transport following the transition from high to low-light (Fig. 6). While DHAP/RuBP ratios remained stable (Fig S9C), the 3PGA/DHAP ratio increased from FL300_290s_HL to FL300_590s_LL (Fig S9A), indicating a growing bottleneck of photochemical supply of NADPH, with the FL300_310s_LL time point falling between levels at FL300_290s_HL and FL300_590s_LL. Thus, the supra-steady-state levels of CO_2_ assimilation at the FL300_310s_LL may have been supported by provision of 3PGA from DHAP and subsequent interconversion to PEP in MC, resulting in continued substrate provision for C_4_ acid formation.

Maize uses both malate and aspartate-based C_4_ acid shuttles (Hatch 1971). It is important to emphasize that our approach used unlabelled metabolites, which although pragmatic, has limitations. In particular, it does not allow for resolution of the metabolic flux, nor for how much of the observed pools are ‘inactive’, i.e., do not turn-over as a result of the imposed conditions. This may be particularly noteworthy for malate, the main transfer C_4_ acid in maize, for which the inactive pool represents ∼60% (Hatch 1971, 1979, Leegood and von Caemmerer 1989, Szecowka et al. 2013, Arrivault et al. 2017, Medeiros et al. 2022), and significant inactive pools have also been observed for other metabolites such as SBP, G1P, and UDPG (Medeiros et al. 2022). Ratios between metabolic pools provide a relative measure of changes in the balance between different metabolic pathways, which can be less sensitive to the presence of inactive pools. Since malate is decarboxylated to pyruvate in the BSC, we used changes in Asp/Pyr as indirect estimates of relative changes in the aspartate and malate shuttles in response to the light treatments. Under low-light steady-state conditions and at FL300_590s_LL, Asp:Pyr showed a significant increase compared to steady-state high-light and to FL300_290s_HL (Fig. S9F). This suggests that the relative increase in aspartate compared to malate as the transfer metabolite, previously observed in low-light-grown plants (Usuda 1987, Doncaster et al. 1989, Ubierna et al. 2013, Medeiros et al. 2022), may also occur transiently during the dynamic light conditions presented here. As discussed by Medeiros et al. (2022), when aspartate is moved to the BSC, it must be coupled with return of an amino group to the MC to maintain N stoichiometry. This could happen via transfer of alanine. The Ala:Pyr ratio increased approximately 2-fold with the change from high to low-light (Fig. S9G), following a similar pattern as Asp:Pyr, in both cases mainly driven by a drop in pyruvate (Fig. S8C). However, changes in the Asp:Pyr ratio were more pronounced, increasing approximately 4-fold, which may suggest that N stoichiometry was further supported by alternative amino-shuttles such as the glutamate:2OG shuttle (Mallmann et al. 2014, Medeiros et al. 2022). Glutamate/2OG (Fig. S9H) varied significantly with sampling time point and were significantly higher at FL300_590s_LL time point compared to the FL300_310s_LL, caused by a significant drop in 2OG concentrations (P<0.05, Fig. S8U).

### Metabolite profiles are strongly affected by low temperature

To find out how different combinations of fluctuating light and suboptimal temperature affected metabolite pools, a second experiment was performed where metabolite profiles were sampled across four timepoints within the 300sFL regime for all three temperatures. The sampling timepoints were at 25 s after switching from low to high light (FL300_25s_HL), 290 s after switching from low to high light (FL300_290s_HL), 25 s after switching from high to low light (FL300_325s_LL), and 290 s after switching from high to low light (FL300_590s_LL), and were selected to maximise variation in *A*_CO2_ (Fig. S10Z). In addition, we sampled at 25 s after switching from low to high light in the 30sFL regime (FL30_25s_HL) to facilitate comparisons with the same sampling time-point at 300sFL (FL300_25s_HL). The effect of temperature on metabolite concentration was significant for 13 metabolites (Temp P<0.05, Fig. S10): aspartate, pyruvate, PEP, 3PGA, DHAP, fructose 6-phosphate (F6P), sedoheptulose 7-phosphate (S7P), ribulose 5-phosphate+xylulose 5-phosphate (Ru5P+Xu5P), glucose 6-phosphate (G6P), UDP-glucose (UDPG), 2-phosphoglycolate (2PG), glycerate and succinate, while significant effects of sampling timepoint (Time P<0.05, Fig. S10) were found for 8 metabolites: PEP, 3PGA, DHAP, sedoheptulose (SBP), ribose 5-phosphate (R5P), ADP-glucose (ADPG), 2PG and 2-oxoglutarate (2OG). Significant interactions between temperature and sampling time (Temp x Time P<0.05, Fig. S10) were found for only three metabolites: PEP, DHAP and 2PG. PCA (Fig. 8) showed that samples primarily clustered by temperature, aligning diagonally to PCA1 (24%) and PCA2 (20%). The four metabolites with the strongest loadings for PC1 were PEP, pyruvate, DHAP and 3PGA (Fig. 8B), which generally showed a decrease under low temperature (Fig. S10B,D, E,G). The second PC on the other hand aligned with variation in many of the phosphorylated sugars, such as F6P, G6P, and glucose 1-phosphate (G1P) (Fig. 8B), which increased under low temperature (Fig. S10H, P, Q).

**Figure 8.**
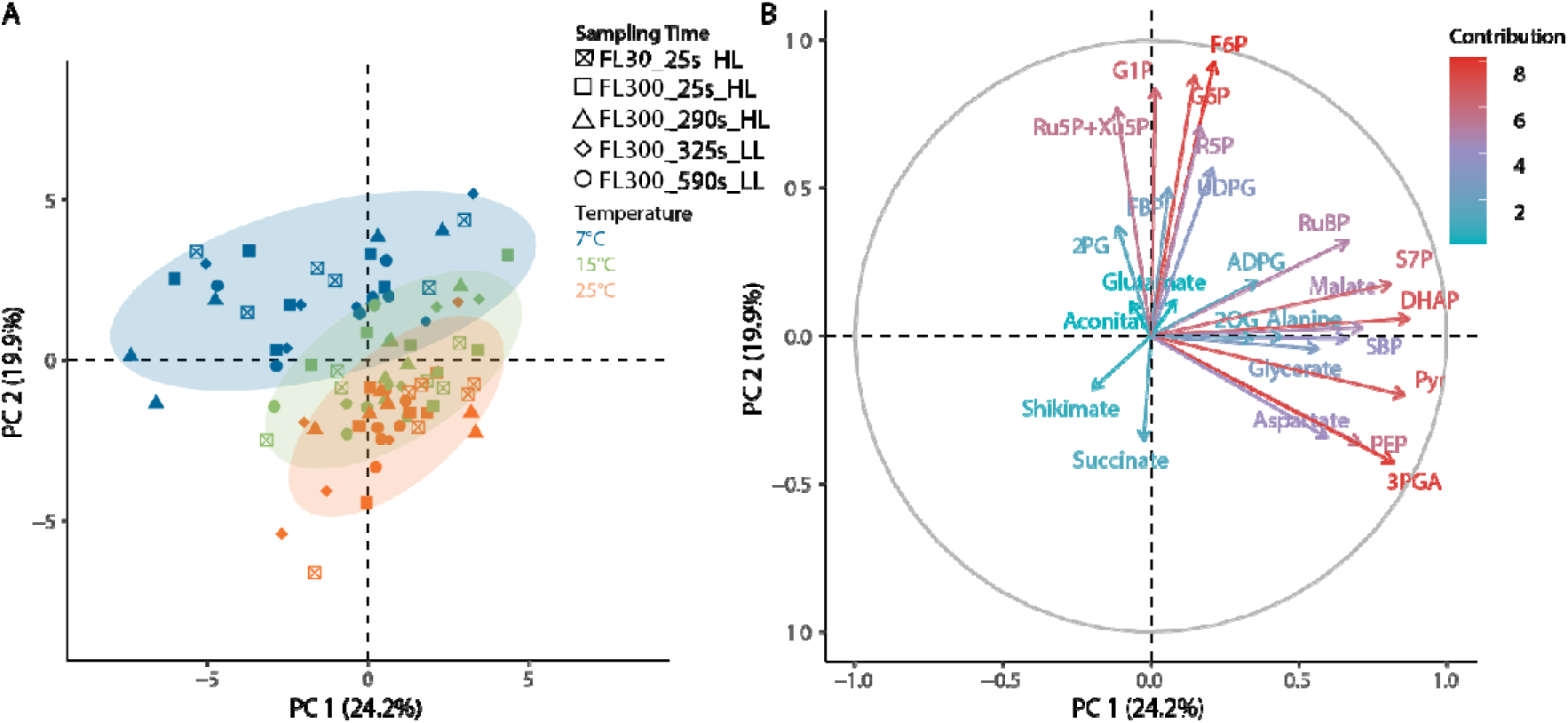
Principal component analysis of metabolite profiles of maize leaves exposed to fluctuating light regimes at three different temperatures. In each fluctuating light regime, leaves were exposed to repetitive changes between low (200 µmol m^−2^ s^−1^) and high (1500 µmol m^−2^ s^−1^) light-steps with duration of either 30 or 300 s (FL30, FL300). For the FL30 light regime, samples were taken 25 s after switching from low to high light (FL30_25s_HL). For the FL300 light regime, samples were taken at four different time points: 25 s after switching from low to high light (FL300_25s_HL), 290 s after light switches from low to high light phase (FL300_290s_HL), 25 s after switching from high to low light (FL300_325s_LL), and 290 s after switching from high to low light (FL300_590s_LL). Plants were acclimated to 25 °C, 15 °C, or 7 °C for at least 2h prior to sampling. Individual sample positions according to the first two PCs are shown in A), with different symbols indicating different sampling timepoints and different colours representing different temperatures. The loadings of individual metabolites are shown in (B). The complete metabolite data are provided in Supplemental Fig.S10.

Whereas at lower temperatures, CO_2_ fixation was relatively similar to steady state at 25 s following each change in intensity, this was not true for the 300sFL regime after the switch to HL at 25 °C where an initial plateau was observed at FL300_25s_HL (Fig 2C, Fig. S10Z). This initial plateau in A_CO2_ was similar to peak A_CO2_ at FL30_25s_HL (Fig 2B, Fig. S10Z), which is why both timepoints were sampled for metabolites. High levels of PEP were observed at FL30_25s_HL and FL300_25s_HL, compared to significantly lower levels at FL300_290s_HL, FL300_325s_LL and FL300_590s_LL (Supplemental Figure S4).. These results suggest that a transient bottleneck in the C_4_ cycle due to insufficient carbon availability could have suppressed A_CO2_ at FL30_25s_HL and FL300_25s_HL. This interpretation is further supported by the *c*_i_ pattern during these analyses (Fig. S11), as both FL30_25s_HL and FL300_25s_HL (Fig S11B and C) show a drop in intercellular CO_2_ concentration (*c*_i_) values with minima close to 100 µmol mol^−1^, which according to the observed *A*/*c_i_* responses (Fig 1B) could have become limiting to A_CO2_. Furthermore, consistent with a transient bottleneck in PEP carboxylation, the ratio between 3PGA/PEP (Fig S12B) was also lowest at time points FL30_25s_HL and FL300_25s_HL, while the ratios of 3PGA/DHAP and DHAP/RuBP (Fig S12A and C) were constant across all sampled high-light time-points showing that flux through the reductive and regenerative parts of the CBB cycle remained tightly coordinated.

### Proportional metabolite distribution between M and BSC is relatively robust under low temperatures

Since C_4_ photosynthesis requires large metabolite pools to drive intercellular metabolite transfer between MC and BSC via diffusion, it is possible that the pronounced low temperature effects on CO_2_ assimilation are in part driven by collapsed metabolite gradients (Labate *et al*. 1990). Because whole-leaf variation in PEP, pyruvate (Pyr), DHAP and 3PGA with temperature strongly determined PC1 in Fig 8, the effect of low temperature on the proportional distribution of these metabolites between MC and BSC was further characterised. Freeze-sampled leaf material was divided into fractions enriched in MC and BSC via serial filtration of leaf homogenates in liquid N_2_ (Stitt and Heldt 1985a). Metabolite proportions in MC and BSC were estimated by normalizing metabolite concentrations in each fraction against activities of phosphoribulokinase (PRK) or PEPC as markers of BSC and MC, respectively (Fig. S13). Whole leaf metabolite concentrations decreased at 7°C relative to 25°C for all four metabolites (Fig. 9A). This was most pronounced for 3PGA, which already decreased significantly at 15°C, whereas DHAP, Pyr and PEP were not significantly lower at 15°C, compared to 25°C. Consistent with previous findings (Stitt and Heldt 1985a,b), 3PGA distribution was highly asymmetric, being entirely partitioned to BSC (Fig. 9C) and undetectable in MC (Fig. 9B) at all three temperatures. In contrast, DHAP and PEP proportions were relatively similar between cell types. For Pyr, the distribution was BSC biased at 25 °C, but became more equal at lower temperatures. The higher proportion of total pyruvate in BSC at 25 °C is inconsistent with the MC biased distribution of labelled pyruvate observed by Arrivault et al. (2017), but the observations of approximately equal BSC and MC proportions at 15 and 7 °C are similar to earlier findings by Leegood (1985) and Stitt and Heldt (1985a,b) who also found pyruvate to be uniformly distributed between both cell types. Whereas the BSC-biased distribution of Pyr at 25 °C shifted to become uniform at 15 and 7 °C, low temperature did not impact the proportional distribution of 3PGA, DHAP and PEP between M and BSC. As a result, diffusion-driven intercellular metabolite transfer of 3PGA, DHAP and PEP is unlikely to be impacted by collapsing gradients beyond the effect of a s decrease in gradient size to due to a decline of whole leaf pool sizes at low temperature (Fig. 9A).

**Figure 9.**
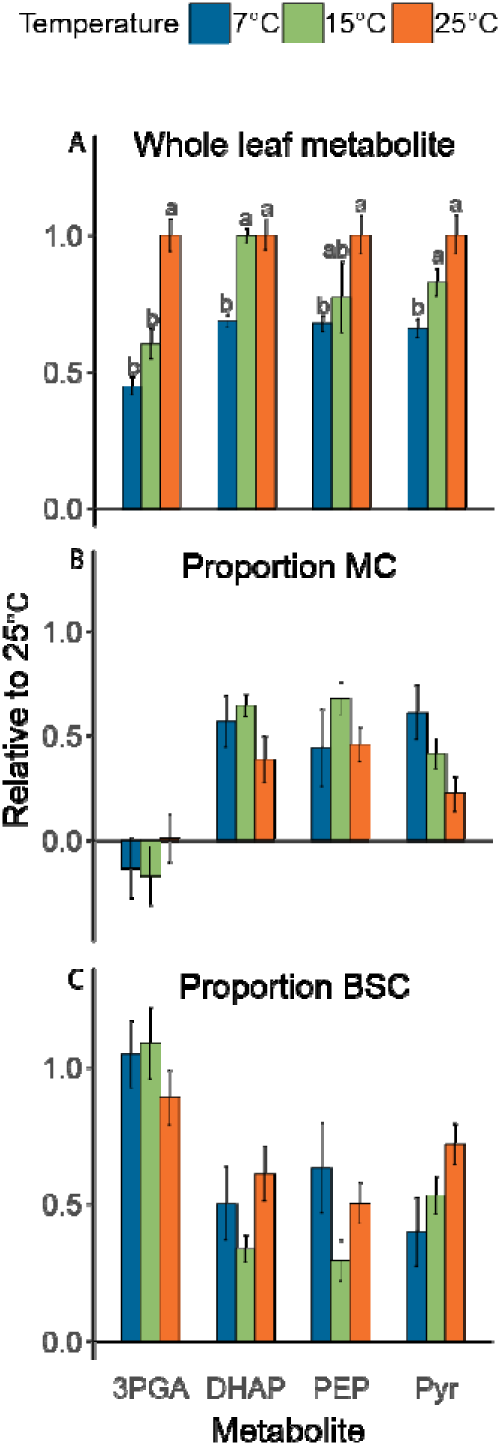
Relative distribution of 3-phosphoglycerate (3PGA), dihydroxyacetone phosphate (DHAP), phosphoenolpyruvate (PEP) and pyruvate (Pyr) between mesophyll and bundle sheath cells in maize leaves exposed to different temperatures. A) decrease of whole leaf metabolite concentrations relative to 25 °C. B) Relative proportion of whole leaf metabolite content partitioned in mesophyll cells. C) Relative proportion of whole leaf metabolite content partitioned in bundle sheath cells. Whole leaf metabolite contents in A) were normalised to 25°C prior to statistical analysis (bars represent means ± SEM; n = 5-6 biological replicates). Different letters indicate statistical differences between different temperatures according to Tukey (*p*<0.05). Mesophyll and bundle sheath proportions were estimated from analysis metabolite contents of fractionated leaf samples obtained from serial filtration over liquid nitrogen (n = 5-6 biological replicates). Mesophyll and bundle sheath metabolite proportions were estimated from linear regression analysis of metabolite concentrations per fraction as a function of fractional activity of marker enzymes for each cell type. Phosphoenolpyruvate carboxylase was used as a marker for mesophyll cells and phosphoribulokinase for bundle sheath cells. Linear regression analysis is summarized in Supplemental Table S2, marker enzyme activities per leaf fraction are provided in Supplemental Fig. S13 and all regression plots are provided in Supplemental Figs 14-17.

## Discussion

To keep C_4_ photosynthesis energetically efficient, it is important to maintain tight coordination between supply and demand of ATP and reductant. This is evident from highly significant linear correlations between rates of electron transfer and net CO_2_ assimilation which are typically found in unstressed plants (e.g., Genty et al 1989). Nevertheless, under fluctuating light at 25°C, this apparent coordination can be lost, which was confirmed here by the strong departures from steady state e^−^PSII/CO_2_ ranging from −64% under low light to +75% under high light, which were found to be related to distinct metabolic profiles. In addition, we show for the first time how these patterns of decoupling are mitigated by suboptimal temperature, with e^−^PSII/CO_2_ becoming essentially invariable with light regimes at the lowest temperature of 7 °C, in contrast to our hypothesis. In the following paragraphs we discuss how our data provides experimental evidence for metabolic buffering suggested previously, as well as the mechanisms underpinning the effect of low temperatures on maize photosynthesis and CO_2_ fixation as evident from whole -leaf and BSC and MC resolved metabolite profiles.

### Metabolic flexibility underpins decoupling between electron transport and CO_2_ fixation at 25°C

Our experiments demonstrate strong transient decoupling between electron transport and CO_2_ assimilation immediately following a change in light intensity at 25 °C (Fig. 6). At 200 µmol m^−2^ s^−1^ Q, e^−^PSII/CO_2_ decreased from 13 ± 2 at steady-state conditions to 5 ± 1 during the 6sFL (Fig. 6E) and 30sFL (Fig. 6F) treatments, and 3 s after the start of the low light phase at the 300sFL treatment (Fig. 6G, Start). The fact that C_4_ plants are able to sustain CO_2_ fixation above steady state rates for a short time after a transition from high to low-light is consistent with previous studies (Laisk and Edwards 1997, Li et al. 2021, Lee et al. 2022, Arce-Cubas et al. 2023a). Large pools of C_4_ transfer metabolites and flexibility in decarboxylation routes have been hypothesized underly this ability. Energy and redox requirements of CO_2_ fixation could be buffered from large metabolite pools and flexibility in energy supply and demand between MC and BSC could be met via adjustments to transfer metabolites (Furbank et al. 2011, Stitt and Zhu 2014, Slattery et al. 2018).

Our results provide experimental evidence for some of these hypotheses. Firstly, rapid changes were observed in the relative contribution of aspartate versus malate shuttles (Fig S8B and 9F), with the former getting more pronounced under low-light. A shift from malate to aspartate shuttles has been reported previously in maize plants measured following a transition from high to low-light (Usuda 1987, Doncaster et al. 1989, Ubierna et al. 2013) as well as in a comparison between steady-state rates at different light intensities (Medeiros et al. 2022) and affects both the redox equivalent moved between MC and BSC and energetic demands of both cell types. Namely, whereas malate transfer moves a redox equivalent from MC to BSC, aspartate transfer does not. And while regeneration of PEP using pyruvate from malate decarboxylation requires two ATP per PEP, decarboxylation of oxalo-acetate via PEPCK regenerates PEP at only one ATP/PEP (Furbank et al. 2011, Bellasio and Griffiths 2014). While the latter may have a quantum yield benefit for the overall pathway (Yin and Struik 2021), another reason suggested to underpin the increased contribution of aspartate at low-light, is that pyruvate movement may require active transport. This was suggested since the intercellular concentration gradient for this metabolite such as observed here for leaves sampled at 25 °C (Fig 9A) are not consistently found (Stitt and Heldt, 1985a, Arrivault et al. 2017). If so, due to the lower supply of ATP during limiting light conditions, pyruvate movement could become limiting or less effective in low-light conditions (Medeiros et al. 2022).

Secondly, PEP equilibrates with 3PGA via the reversible phosphoglycerate enolase and mutase reactions, which may help to buffer metabolite levels in the CBB cycle and the C_4_ cycle upon changes in light intensity (Huber and Edwards 1975, Furbank and Leegood 1984, Leegood and von Caemmerer 1989, Medeiros et al 2022). At equilibrium, 3PGA should be approximately two to four-fold higher than PEP levels (Leegood and von Caemmerer 1988, Ubierna et al. 2013, Stitt and Heldt 1985, Medeiros et al. 2022). This agrees well with the observed steady-state ratios around 3.5 (Fig. S9B). However, after the transition from high to low-light (FL300_310s_LL) the 3PGA/PEP ratio jumped to ∼7.8, which is significantly higher than the steady-state values. Equilibration of isotopic label between 3PGA and PEP was estimated to occur at 18-30% of the rate of CO_2_ fixation (Medeiros et al. 2022), which seems consistent with the transient departures from equilibrium ratios observed here. The increase in 3PGA/PEP was mostly due to decreases in PEP (Fig. S8E), suggesting that PEP carboxylation continued transiently at supra-steady-state rates following the switch from high to low-light, while PEP regeneration from pyruvate was downregulated more rapidly. Continued C_4_ cycling may also have driven 3PGA formation in BSC above low-light steady-state rates, in which case spatial separation of MC and BSC pools could further explain the transient increase in 3PGA/PEP ratio. In BSC cells 3PGA/PEP ratios of up to 20 have been observed (Stitt and Heldt, 1985ab), possibly due to the asymmetric distribution of PGA mutase and enolase activity between cell types, with the major fraction found in MC (Furbank and Leegood, 1984).

Thirdly, the reversible CBB cycle reactions between 3PGA and triose phosphates have been suggested to act as a buffer for ATP and NADPH (Stitt and Zhu 2014). Previous measurements of 3PGA/DHAP across increasing irradiance levels showed a steady decrease, inversely proportional to CO_2_ fixation rate (Leegood and Von Caemmerer 1988). Here, a significant increase in the 3PGA/DHAP ratio was observed at the end of the low-light phase (FL300_590s_LL, Fig S9A) compared to the end of the high-light phase (FL300_290s_HL). This increase under limiting light conditions is consistent with other findings (Usuda 1987, Leegood and von Caemmerer 1988) and can be explained by restriction of the conversion of 3PGA to triose phosphates in MC under low-light, leading to accumulation of 3PGA. The fact that 3PGA/DHAP gradually increases from FL300_290s_HL to FL300_590s_LL, with FL300_310s_LL showing intermediate values (Fig. S9A) while photosynthetic electron transport per unit CO_2_ fixed was significantly below steady-state following the switch from high to low-light (Fig 6G), suggests that conversion of DHAP to 3PGA may have contributed to the provision of NADPH and ATP for supra-steady-state CO_2_ fixation at low-light during the FL300_310s_LL time point (Fig. S8Z).

### Low temperature strengthens coupling between electron transport and CO_2_ assimilation during fluctuating light

Previous observations of maize plants exposed to chilling conditions showed a strong uncoupling of electron transport and CO_2_ fixation (e.g., Fryer et al. 1998), with rates of electron transport greatly exceeding those predicted by the ATP and NADPH demands of CO_2_ assimilation. On this basis, we hypothesized that short-term exposure to chilling temperature might have a similar effect, and potentially aggravate the decoupling caused by fluctuating light. Surprisingly the opposite was found, with chilling temperatures instead negating the decoupling effect of fluctuating light (Fig. 6). This suggests that the alternative electron sinks observed previously by Fryer et al. (1998) require longer adaptation to chilling and do not manifest during short-term exposure to low temperature. This could be consistent with previous work which shows that inhibition of CO_2_ assimilation was only marginal after 2 hours of low temperature exposure but became much more pronounced at longer treatment time (Long et al. 1983). Thus, despite their inherent cold-sensitivity, species like maize can tolerate short-term exposure to low temperature.

The enhanced coupling observed here under low temperature is remarkable, considering the strong decline in CO_2_ assimilation rate, meaning that the potential mismatch between the absorbed energy to generate NADPH and ATP from the light reactions and the demand for these products in downstream metabolic pathways should have become strongly unbalanced. The fact that this did not lead to uncoupling can be explained by two potential mechanisms, which are not mutually exclusive. On the one hand, the negative feedback of lower NADP+ and ADP regeneration via decreases in thylakoid lumen pH would provide a strong trigger to slow down photosynthetic electron transport via induction of sustained NPQ (Supplemental Fig. S4 and S15) and downregulation of whole-chain electron transport. The latter is particularly evident in the 7°C measurements at low-light (Fig. 5E-H), where ϕ_PSI_ does not recover fully, despite the low-light levels. This may reflect a constriction to electron transfer at the Cyt*b*_6_,*f* complex to keep plastocyanin and P700 in oxidized state, as observed previously by Labate et al. (1990), which together with the induction of NPQ would act to prevent runaway ROS formation. Secondly, the chilling temperature had a marked impact on the difference in A_CO2_ between high light and low light, which drastically decreased due to saturation of A_CO2_ at significantly lower light intensity (Fig 1A) as evident from the significant lack of reoxidation of ferredoxin (Fig. S6E-H) and Q_A_ (Fig S3E-H) under low light at 7 °C. As a result, the thylakoid reactions would have to undergo less adjustment following each change in light intensity to accommodate for changes in downstream demand for ATP and NADPH, which instead stayed rather constant between contrasting light levels.

In agreement with previous studies (Labate et al. 1990) 3PGA, PEP, pyruvate, and aspartate decreased with a decrease in temperature. The content of 3PGA was drastically reduced in this experiment, with only minor changes observed in DHAP, therefore the 3PGA:DHAP ratio was also reduced under low temperatures (Fig. S12A). G6P and F6P increased as the temperature decreased (Fig. S10I and P), which is inconsistent with results by Labate et al. (1990), who found that G6P remained constant and F6P decreased with decreasing temperature. Nevertheless, the observed increase in the G6P:F6P ratio at low temperature (Fig. S12A) is in agreement with previous work and reflects a shift towards the cytosolic compartmentation of these compounds (Gerhardt et al. 1987; Labate et al. 1990). Despite the increases in F6P and G6P, FBP content did not change significantly with temperature, which is consistent with observations by Labate et al. (1990) and reflects coordination of temperature-dependent downregulation of sucrose synthesis with the decrease in DHAP at low temperature. Assuming that FBP follows similar changes to DHAP (as they are linked via aldolase), low temperature may lead to a drop in FBP in the BSC but little change in the MC. This deduced maintenance of FBP levels in the MC could reflect inhibition of cytosolic FBPase at low temperature. Indeed, cytosolic FBPase is especially inhibited by low temperatures in C_3_ plants as it becomes more sensitive to inhibition by fructose 2,6-bisphosphate and adenosine 5′-monophosphate (AMP) as the temperature falls (Stitt and Grosse 1988). It is known that in C_4_ plants FBPase has a higher Km than in C_3s_ (Stitt and Heldt 1985b). While no studies have been performed for maize FBPase under lower temperature, based on our observations we suggest that its response is likely similar to those observed in C_3_ isoforms of the enzyme. If so, the rise in hexose phosphates but not in DHAP presumably reflects different impact of temperature on the balance between SPS and cytosolic FBPase.

### Proportional metabolite distribution between BSC and MC shows only minor impact of temperature

Labate et al. (1990) observed a strong decline in phosphorylated intermediates at low temperature in maize, in contrast with observations in barley, which showed an increase. Our measurements are consistent with the observations by Labate et al. (1990) on maize, and suggest that availability of inorganic phosphate did not restrict ATP regeneration. Based on their findings, Labate et al. (1990) hypothesized that the decrease in DHAP reflected a decrease in the rate of 3PGA transfer between MC to BSC and a decline in the diffusional gradient that is required to drive this movement. If so, the equilibration via phosphoglycerate mutase and enolase between 3PGA and PEP might result in decreased PEP in the MC because 3PGA transfer from BSC to MC would decrease under low temperature.

Here, we partially confirm these hypotheses. The diffusional gradient of 3PGA clearly declined under low temperature, but this was entirely due to a decrease in the BSC, since 3PGA in MC remained undetectably low across all conditions (Fig. 9B). Our data did not show evidence of strong equilibration between 3PGA and PEP, since PEP showed a more uniform distribution and whole-leaf pool sizes were less affected by temperature than 3PGA. Similar to PEP and 3PGA, the proportional distribution in DHAP also did not show a clear pattern with temperature, but the diffusional gradients for all three metabolites would have decreased with declines in whole leaf pools, most strongly for 3PGA (Fig. 9A). These findings are consistent with a mechanism whereby at 25°C 3PGA accumulates in BSC due to a shortfall in NADPH, and the accumulated 3PGA diffuses to the MC where it is reduced. At low temperature less 3PGA will accumulate due to the negative effect of low temperature on flux in the CBB cycle and Rubisco activity. Lower 3PGA accumulation in BSC in turn should drive a lower rate of diffusion to match the lower rate of photosynthesis. Assuming that transfer of 3PGA and DHAP is fully reliant on diffusion, flux would be expected to be exponentially dependent on absolute temperature and therefore change only moderately if the diffusional gradient is maintained. Of course, this is not the case here, but the diffusional decreases due to absolute temperature (∼6%) and whole leaf 3PGA concentration (55%) at 7 °C both seem too small to fully explain the strong decrease of 86% in light saturated CO_2_ assimilation rate (Fig 1 and 2). We therefore speculate that the latter may reflect additional restrictions under low temperature. It is well-known that plasmodesmata density are strongly enhanced in C_4_ compared to C_3_ leaves (Danila et al. 2019) to facilitate the diffusion of metabolites between mesophyll and bundle sheath cells (Hatch and Osmond 1976). Interestingly, closure of plasmodesmata in maize after four hours exposure to low temperature by the accumulation of callose and calreticulin has been reported (Bilska and Sowiński 2010). Thus, the disproportionate decrease in CO_2_ assimilation rate under low temperature relative to more moderate changes in metabolite concentration and rate of diffusion, may also be partly explained by increased diffusional resistance to metabolite transfer through the plasmodesmata under low temperature.

## Conclusion

Coordination between NADPH and ATP provision from the thylakoid reactions and flux through the C_4_ and CBB cycles is well-known to be important for efficient C_4_ photosynthesis. Here we investigated the interplay between low temperature and fluctuating light, two conditions previously suggested to lead to substantial decoupling. Our observations confirm significant decoupling in response to fluctuating light, provide evidence for mechanisms of metabolic flexibility which underpin these departures from steady state, but unexpectedly showed a tighter coupling between electron transfer and CO_2_ fixation at low temperature. We propose that the latter reflects both strong downregulation of electron transport to avoid excessive ROS formation, as well as a stronger degree of light saturation of CO_2_ assimilation at low temperature across both light levels used in our FL regimes, which may have decreased the need for extensive electron transfer adjustment following changes in light. How and when this situation of tight control transitions into the strong decoupling observed in C_4_ leaves following prolonged exposure to suboptimal temperature will be subject for further research.

## Material and Methods

### Plant growth conditions

Seeds of maize inbred line B73 were sown in Levington Advance M3 compost (Scotts, Ipswich, UK) in seed trays. After one week, seedlings were transplanted to 2 L pots (2 plants per pot), containing a mixture of 2:2:1 of Levington M3 compost: Top Soil (Westland, Dungannon, UK): Perlite 2.0-5.0 mm (Sinclair, Ellesmere Port, UK). Each pot was supplemented with 5 g of slow release 17N-9P-11K fertiliser (All Purpose Continuous Release Plant Food, Scotts Miracle-Gro, Marysville, OH, USA), 5 g of magnesium salts (Scotts Miracle-Gro, Marysville, OH, USA), and 10 g of garden lime (Westland, Dungannon, UK). Plants were grown in a Conviron walk-in controlled conditions growth chamber (Conviron Ltd., Winnipeg, MB, CA) at 28/20°C day/night with a photoperiod of 14 h, photosynthetic photon flux density (*Q*) of 600 μmol m^−2^ s^−1^, and 65% humidity, and watered every other day. Plants were grown under these conditions until stage V5 (fifth completely expanded leaf). Prior to physiological measurements, plants were transferred to a controlled environment chamber (Percival E-41HO, Perry, IA, USA), set up to 25°C, 15°C, or 7°C, and dark acclimated for one hour before the measurements were performed unless mentioned otherwise.

### Gas exchange measurements

After an hour of dark acclimation at the desired temperature, lights in the cabinet were turned on to a *Q* of 600 μmol m^−2^ s^−1^. Response curves of CO_2_ assimilation rate (*A*_CO2_) to increasing intercellular CO_2_ concentration (*c*_i_*; A*_CO2_x*c*_i_ curves), photosynthetic responses to steady light (light response curves; *A*x*Q* curves), and to fluctuating light conditions were performed on the youngest completely expanded leaf using a Li-6800 portable infrared gas analyser (IRGA) system (software version 1.4.05, LI-COR, Lincoln, NE, USA) with a 9 cm^2^ leaf chamber (6800-12A) equipped with a 6800-02 light source.

For the *A*_CO2_x*c*_i_ and *A*x*Q* curves the conditions inside the chamber were 410 μmol mol^−1^ reference CO_2_ concentration (CO_2__r), 55% relative humidity, and flow rate of 600 µmol s^−1^. Leaf temperature was controlled according to the temperature that plants were measured: 25°C, 15°C, or 7°C. For the *A*x*c*_i_ curves, leaves were acclimated at 1700 µmol m^−2^ s^−1^ PPFD actinic red light to allow *A*_CO2_ and stomatal conductance (*g*_s_) to reach steady-state. Subsequently, gas exchange was measured in the following CO_2__r concentrations: 1000, 850, 750, 600, 410, 300, 200, 150, 120, 100, 80, 60, 40, 20, and 410 μmol mol^−1^. Gas exchange parameters were logged between 120 and 180 s at each step, and before logging, the reference and sample IRGA signals were matched. The *A*_CO2_ x *c*_i_ response curves were fit to a nonrectangular hyperbolic function (von Caemmerer 2000). The initial part of the curve was used to estimate the maximum carboxylation rate of phosphoenolpyruvate carboxylase (*V*_pmax_). A linear model of *A*_CO2_ as a function of *c*_i_ was fitted and the breaking point detected. The response of *A*_CO2_ to *c*_i_ < breaking point was used to solve *V*_pmax_, and *K*_p_, the apparent Michaelis-Menten constant of PEPC for CO_2_, assumed to be 60, 93, and 154 μbar at 7, 15 and 25°C, respectively (Boyd et al. 2015). The *c*_i_-saturated rate of photosynthesis (*V*_max_) was estimated as the predicted value of each function for *c*_i_ > 2000 μmol mol^−1^.

For the *A*x*Q* curves, leaves were acclimated at 2100 µmol photons m^−2^ s^−1^ actinic red light to allow *A*_CO2_ and stomatal conductance (*g*_s_) to reach steady-state. Incident light intensity was then stepped down through 1700, 1350, 1000, 670, 500, 360, 260, 180, 130, 80, 40, and 0 µmol photons m^− 2^ s^−1^. Gas exchange parameters were logged between 60 and 180 s at each step, and before logging, the reference and sample IRGA signals were matched. Light response curves were fitted by a non-rectangular hyperbola (Marshall and Biscoe 1980) to estimate mitochondrial respiration (*R*_d_), maximum *A*_CO2_ assimilation under saturating light (*A*_sat_), convexity (θ), and light compensation point (LCP). Instantaneous quantum efficiency of CO_2_ assimilation (□CO_2_) was calculated at each light level as □CO_2_ = (*A*_CO2_ + *R*_d_)/ α_leaf_, where α_leaf_ is leaf light absorptance measured at 630 nm (as just red light was used in the experiment) in three maize plants grown under same conditions as the ones used in the experiment, and at the same developmental stage with an integrating sphere (Li-1800-12, LI-COR, Lincoln, NE, USA) optically connected to a miniature spectrometer (STS-VIS, Ocean Insight, Orlando, FL, USA) following manufacturer instructions (LI-COR, 1988). Leaf absorptance at 630 nm was 0.904 ± 0.002, and it was calculated as α_leaf_ = 1-*T*_s_-*R*_s_, where *T*_s_ and *R*_s_ are transmittance and reflectance of a diffuse sample, respectively.

To measure photosynthetic responses to fluctuating light (FL), leaves were first acclimated at 600 µmol m^−2^ s^−1^ actinic red light, until *A*_CO2_ and *g*_s_ reached constant levels. Using a custom program, leaves were then exposed to repetitive stepwise fluctuations in light intensity from 1500 to 200 µmol m^−2^ s^−1^ *Q* for 1 h, with gas exchange parameters logged every 2 s. Three different light treatments were tested, with each light step lasting 6, 30, or 300 s. To avoid interference with the shorter fluctuations and the data sampling interval, averaging time for head measurements was set to zero, with no additional averaging. Therefore, each log represented an average of the preceding 0.5 s, the inverse of the instrument digital update frequency of 2 Hz. The IRGAs were matched before the fluctuating light program started.

As described by Arce-Cubas et al. (2023a), a storage flux correction was applied to the measurements under light fluctuations, as they violate the steady-state assumption in default assimilation rate equations. These corrections followed the same principle stated by Saathoff and Welles (2021). Based on the mass balance of the instrument cuvette, the derivative of the cuvette concentration over time can be used to adjust *A*_CO2_ and apparent transpiration rates. A storage flux term for CO_2_ and H_2_O was computed from changes in cuvette concentration between measurements taken every 2 s and applied to adjust *A*_CO2_ and transpiration rates according to Equations 1 and 2, respectively.

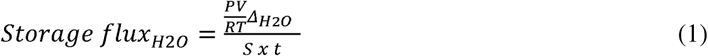

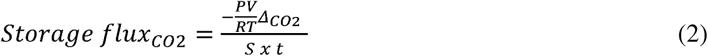

In equations 1 and 2, *P* represents pressure (*P*_a_, from instrument), *V* represents cuvette volume (10.72 × 10^−5^ m^3^), *R* represents the molar gas constant, and *T* represents air temperature inside the cuvette to calculate the change in moles of gas (ΔCO_2_ or ΔH_2_O) using instrument recordings of current and preceding logs. *S* represents leaf area (m^2^) and *t* represents the time since last log (s) and was used to convert the molar concentrations to flux per area.

### 2.3 Dual-Klas-NIR measurements

A Dual-KLAS-NIR (DKN) analyser (Heinz Walz GmbH, Effeltrich, Germany) was used in parallel to gas exchange measurements to access chlorophyll fluorescence, and P700 redox changes (Klughammer and Schreiber 2016; Schreiber and Klughammer 2016). The DKN was equipped with a gas-exchange cuvette (3010-DUAL), connected to a LI-6800 portable infrared gas analyser (IRGA) system (software version 1.4.05, LI-COR, Lincoln, NE, USA) through a custom chamber manifold (6800-19), to control the conditions inside the DKN chamber to a flow rate of 200 µmol s^−1^, CO_2__r of 410 µmol mol^−1^, and 55% relative humidity. Leaf temperature was controlled through a 3010-I/Box with the GFS-Win software (Heinz Walz GmbH, Effeltrich, Germany), according to the temperature that plants were measured: 25°C, 15°C, or 7°C. Before each measurement, plants were dark adapted for 1 h at the desired temperature, and the four pairs of pulse modulated NIR measuring sensors were zeroed and calibrated. Differential model plots (DMP) were generated and maximum oxidation of P700 was determined running the NIRmax script as described by the manufacturer and used to normalize redox-associated absorption changes.

Minimum and maximum dark-adapted fluorescence (*F*_0_ and *F*_m,_ respectively) were determined after 1 hour of dark-adaptation. During the light response curves and FL responses, chlorophyll fluorescence parameters were determined by using a measuring light intensity of 20 μmol m^−2^ s^−1^, and saturating pulse of 25,736 μmol m^−2^ s^−1^. For the PSI redox changes, measuring light intensity was 14 μmol m^−2^ s^−1^.

Photosystem II quantum yield (ϕ_PSII_) was calculated as ϕ_PSII_ = (*F*_m_^’^ – *F’*)/ *F*_m_^’^, where *F*_m_^’^ is the maximal fluorescence yield from a light-adapted leaf, and *F*’ is the steady-state fluorescence emission from leaves under actinic light. Photosystem I quantum yield (ϕ_PSI_) was calculated from NIR absorptance changes as ϕ_PSI_ = (*P*_m_^’^− *P*)/ *P*_m_, where *P*_m_ is P700 in the fully oxidised state, *P*_m_^’^ is the maximum change of the deconvoluted P700 signal from light-adapted leaf after a saturating pulse, and *P* is the steady-state P700 signal. For the steady-state light response curves, flashes were applied after 3 min at each light intensity. For FL measurements, flashes were applied at the beginning of the high-light phase (at 1803 s for all cycles), and low-light phase (2103 s for 300 s cycle, and 2133s for 30 and 6 s cycles); at the end of the high light phase (2697 s for 300 s cycle; and 2667 s for 30 and 6 s cycles) and low-light phase (at 2997 s for all cycles). In all the time points described above, flashes were applied 3 s before or after the light switch (see scheme of flashes in Fig. S2).

The electron transport rate per assimilated CO_2_ was estimated by the ϕ_PSII_/□CO_2_ ratio (Genty et al. 1989, Oberhuber and Edwards 1993), denoted here as e^−^PSII/CO_2_ fixed. Non photochemical quenching (NPQ) was calculated as (*F*_m_*−F*_m_□) /*F*_m_□. Quinone A (Q_A_) redox state (1-*q*L) was calculated according to Kramer et al. (2004) where *q*_L_=(1/*F*O*−*1/*F*_m_□) / (1/*F*_o_□*−*1/*F*_m_□).

### Metabolites

#### a) Temperature and FL: sampling for metabolites profiles

Plants grown at the same condition and at the same phenological stage as described for physiological measurements were used for metabolite sampling. Plants were acclimated for 1 h at the desired temperature (7, 15 or 25°C) before starting the light fluctuating cycle. Specific sampling times in both experiments are indicated in Fig S8Z and S10Z.

To ensure accurately time metabolite sampling, a fast quenching system described by Xu et al. (2021) was used. Briefly, the leaf was enclosed in a LI-6800 IRGA with a 9 cm^2^ leaf chamber (6800-12A) equipped with a 6800-02 light source, set up with the same environmental conditions as described in Materials and Methods section 2.2.1 “Gas exchange measurements”. The fluctuating light cycle was started, and sampling was performed at least 30 minutes after the beginning of the program to avoid initial variation in the fluctuation patterns. To quench metabolism at a given time point, the 6800-12A leaf temperature thermocouple was removed and liquid nitrogen was immediately sprayed onto the leaf surface using a cryospray nozzle, through the thermocouple port. The frozen leaf portion was then quickly removed, dropped inside liquid nitrogen, and placed into a pre-cooled microtube. Samples were subsequently stored at –80°C until metabolite analyses.

#### b) Temperature: sampling for cell enrichment (gradient)

For the cell separation experiment, plants were acclimated for one hour at the desired temperature (25, 15 or 7°C) with the lights in the growth cabinet turned on to a *Q* of 600 μmol m^−2^ s^−1^. Afterwards, the youngest completely expanded leaf was clamped into a 9 cm^2^ leaf chamber at 1500 µmol m^−2^ s^−1^ actinic red light. Gas exchange data was logged every two seconds for 30 min, and metabolism was quenched at steady state after 30 min (Fig. S18). Eleven samples from individual plants were pooled together for each cell separation, totalling 6 pooled replicates of 11 plants each. This was needed to provide ∼1 g of fresh weight (FW) which is required for the cell separation protocol. Sampled maize leaves were fractionated as in Stitt and Heldt (1985ab). Four fractions were obtained by homogenizing ∼1 g FW material, resuspending and filtering sequentially through 200, 80, and 40 µm nylon meshes in the presence of liquid N_2_ (Sefar, Switzerland). Following leaf fractionation, PEP, pyruvate, 3PGA, and DHAP concentrations were determined for each fraction (see section c). In addition, activity of the MC marker enzyme phosphoenolpyruvate carboxylase (PEPC) was measured as in Gibon et al. (2004) with 400-fold dilution (FW/extract volume). For the BSC, activity of the marker enzyme phosphoribulokinase (PRK) was measured according to Leegood (1990) with a 200-fold dilution (FW/extract volume). For each metabolite, the ratio of (PEPC/PRK; x-axis) and the ratio (metabolite X/PRK; y-axis) were plotted against each other, and a linear regression calculated (Fig S14-17). The intercept on the y-axis represented the proportion of metabolite X in the BSCs. If plotted against the alternative marker enzyme, i.e., PRK/PEPC on the x-axis and the ratio metabolite X/PEPC on the y-axis, the y-intercept represented the proportion of metabolite X in the MCs.

#### c) Metabolite analyses

Maize material was ground to fine powder using a ball mill (Tesch, Haan, Germany) at liquid N_2_ temperature and stored at –80°C. Samples were analysed by LC–MS/ MS and GC–MS with reference standards for accurate metabolite quantification as in Arrivault et al. (2009). The total amounts of PEP, pyruvate, 3PGA, and DHAP were determined enzymatically in freshly prepared trichloroacetic acid extracts as described in Merlo et al. (1993) using a spectrophotometer (Shimadzu, Kyoto, Japan). Alanine was determined enzymatically. 40 μL of extract was added to an assay buffer containing 0.1 M Tris-HCl pH 10.1, 2 mM EDTA, and 50 mM NAD+. Reactions were performed at 30°C after adding 0.5 U μL^−1^ of alanine dehydrogenase.

### Statistical analysis

Statistical analyses were performed in R 4.3.2 (R Core Team 2023) on RStudio (2023.12.1, Posit Team 2023). One-way, two-way, or three-way ANOVA was used to test the effect of temperature (7°C vs. 15°C vs. 25°C), fluctuation length (6s, 30s, 300s, and steady state), and measurement time (start or end of the light fluctuation) on the different parameters measured during this study (described in detail on each figure caption). Data for the different traits were tested for homogeneity of variances by Levene’s test (α = 0.05) and normality of studentized residual distribution using Shapiro-Wilk test (α = 0.05). When these tests were not satisfied, variables were transformed prior to ANOVA or non-parametric tests were applied (as indicated in the figure captions). When ANOVA effects were significant at the 95% confidence level, Tukey post hoc comparisons were used to compare group means. All plots were generated using ggplot2 (Wickham 2016). For light response curve fitting, package *segmented* was used (Muggeo 2008). For the area under the curve to obtain integrated *A*_CO2_, bayestestR library was used (Makowski et al. 2019). Package *factoextra* was implemented for principal component analyses (Kassambara and Mundt 2020).

## Acknowledgments

We thank Prof Julian Hibberd at University of Cambridge for providing the B73 seeds used in the study, Dr Shaun Nielsen (NSW Health, Sydney, Australia) for help with data analysis; and Dr Chiara Airoldi and Diana Reis (Max Planck Institute of Molecular Plant Physiology, Germany) for assistance in the laboratory work during visit to Max Planck Institute.

## Author contributions

J.K. and C.R.G.S. conceived the study. J.K and C.R.G.S designed the experiments. C.R.G.S. carried out all physiology experiments and part of the metabolite analyses, all data analysis and interpretation, and drafted the manuscript with input of J.K. S.A. provided support with all the metabolite methodology, and carried out most of the metabolite analyses; T.T. and V.C. provided support and helped with the cell enrichment experiment; R.L.V. helped with the fluctuating light experimental setup and provided support with gas exchange experiments; L.A.C. helped with data analyses and interpretation; M.S. hosted C.R.G.S at Max Planck Institute of Molecular Plant Physiology and provided valuable guidance in the metabolite data interpretation. All authors contributed to the manuscript and approved the final version.

## Funding

This work was supported by the Biotechnology and Biological Sciences Research Council via grant BB/T007583/1 and the UK Research and Innovation - Future Leaders Fellowships scheme via an award to JK (MR/T042737/1).

## Conflict of interest statement

None declared.

## Data availability

All data obtained for this study are presented within the supplementary materials and main manuscript.

## Supplemental data

The following materials are available in the online version of this article.

**Supplemental Table S1.** Parameters estimated from the *A*x*Q* curve and *A*x*c*_i_ curves performed in maize plants acclimated at 7, 15 or 25°C.

**Supplemental Table S2**. Slopes and intercepts from linear regression analysis of metabolite contents in fractionated leaf tissue.

**Supplemental Figure S1.** Net CO_2_ assimilation (*A*_CO2_) in maize plants measured under three different fluctuating light regimes.

**Supplemental Figure S2.** Scheme showing timing of chlorophyll fluorescence and P700 redox changes measurements in maize plants under three different fluctuating light regimes.

**Supplemental Figure S3.** Quinone A redox state in maize leaves as a function of temperature and fluctuating light regime.

**Supplemental Figure S4.** Non-photochemical quenching (NPQ) in maize leaves as a function of temperature and fluctuating light regime.

**Supplemental Figure S5.** Plastocyanin redox state in maize leaves as a function of temperature and fluctuating light regime.

**Supplemental Figure S6.** Ferredoxin redox state in maize leaves as a function of temperature and fluctuating light regime.

**Supplemental Figure S7.** Instantaneous quantum yield of CO_2_ fixation (ϕ_CO2_) in maize leaves as a function of temperature and fluctuating light regime.

**Supplemental Figure S8.** Metabolite profiles of maize leaves exposed to constant or fluctuating light.

**Supplemental Figure S9.** Selected ratios between metabolite profiles of maize leaves exposed to constant or fluctuating light.

**Supplemental Figure S10.** Metabolite profiles of maize leaves exposed to fluctuating light regimes at three different temperatures.

**Supplemental Figure S11.** Intercellular CO_2_ concentration (*c*_i_) in maize plants measured under three different fluctuating light regimes.

**Supplemental Figure S12.** Selected ratios between metabolite profiles of maize leaves exposed to fluctuating light regimes at three different temperatures

**Supplemental Figure S13.** Activities of phosphoribulokinase (PRK) and phosphoenolpyruvate kinase (PEPC) in whole leaf extract (WL) and in four leaf fractions.

**Supplemental Figure S14.** Linear regression analysis of pyruvate concentration in four leaf fractions.

**Supplemental Figure S15.** Linear regression analysis of phosphoenol-pyruvate (PEP) concentration in four leaf fractions.

**Supplemental Figure S16.** Linear regression analysis of 3-phosphoglycerate (3PGA) concentration in four leaf fractions.

**Supplemental Figure S17.** Linear regression analysis of dihydroxyacetone-phosphate (DHAP) concentration in four leaf fractions.

**Supplemental Figure S18.** Leaf CO_2_ assimilation (*A*_CO2_) logged before sampling for fractionation of leaf samples.

## Notes

### Competing Interest Statement

The authors have declared no competing interest.

