## Supplemental Table S1-2 and Fig S1-18 for "Sub-optimal temperature leads to tighter coupling between photosynthetic electron transport and CO_2_ assimilation under fluctuating light in maize"

**Running head:** Effects of fluctuating light and low temperature on maize photosynthesis

Cristina R. G. Sales^1,2^, Stéphanie Arrivault^3^, Tomás Tonetti^4^, Vittoria Clapero^3^, Richard L. Vath^1,5^, Lucía Arce Cubas^1^, Mark Stitt^3^, Johannes Kromdijk,^1,6*^

^1^ Department of Plant Sciences, University of Cambridge, Cambridge CB2 3EA, UK

^2^ Wild Bioscience, Abingdon, OX14 4SA, UK

^3^ Max Planck Institute of Molecular Plant Physiology, Am Muehlenberg 1, D-14476 Potsdam-Golm, Germany

^4^ Instituto de Agrobiotecnología del Litoral (IAL-CONICET), Facultad de Bioquímica y

Ciencias Biológicas, Universidad Nacional del Litoral, Santa Fe, Argentina.

^5^ LI-COR Biosciences, Lincoln, NE 68504, United States.

^6^ Institute for Genomic Biology, University of Illinois at Urbana-Champaign, IL61801 Urbana, USA.

**Supplemental Table S1.** Parameters estimated from the *A*x*Q* curve and *A*x*c*_i_ curves performed in maize plants acclimated at 7, 15 or 25°C.

**Supplemental Figure S18.** Leaf CO_2_ assimilation (*A*_CO2_) logged before sampling for fractionation of leaf samples.

**Supplemental Table 1.** Parameters estimated from the response curves of leaf CO_2_ assimilation (*A*_CO2_) to increasing photosynthetic active radiation (*Q*, *A*x*Q* curve), and to increasing intercellular CO_2_ concentration (*c*_i_, *A*x*c*_i_ curve) in maize plants acclimated at 7, 15 or 25 °C.

| **Parameter** | **Curve** | **7°C** | **15°C** | **25°C** | ***p*-value** |
| --- | --- | --- | --- | --- | --- |
| *A*_sat_ (μmol m^-2^ s^-1^) | *A*x*Q* | 4.8±0.7^c^ | 16.8±1.2^b^ | 35.1±2.5^a^ | <0.001 |
| φCO_2_ (μmol mol ^-1^) |  | 0.039±0.005^b^ | 0.062±0.004^a^ | 0.068±0.004^a^ | 0.001 |
| *R*_d_ (μmol m^-2^ s^-1^) |  | 0.10±0.07^c^ | 0.73±0.07^b^ | 1.36±0.11^a^ | <0.001 |
| LCP (μmol m^-2^ s^-1^) |  | 2.7±1.9^c^ | 12.0±0.7^b^ | 20.6±1.3^a^ | <0.001 |
| θ (dimensionless) |  | 0.737±0.141^a^ | 0.300±0.040^b^ | 0.482±0.095^ab^ | 0.034 |
| *V*_pmax_ (μmol m^-2^ s^-1^) | *A*x*c*_i_ | 11.5±2.7^c^ | 34.0±3.3^b^ | 73.2±4.0^a^ | <0.001 |
| *V*_max_ (μmol m^-2^ s^-1^) |  | 7.0±1.0^c^ | 15.5±1.0^b^ | 34.9±2.6^a^ | <0.001 |

Abbreviations: *A*_sat_, *A*_CO2_ at saturating *Q* and ambient CO_2_ (41 Pa); ϕCO, maximum quantum yield of CO_2_ assimilation; *R*_d_, dark respiration, LCP, light compensation point; θ, convexity of the non-rectangular hyperbolic response of *A*_CO2_ to increasing *Q*; *V*_pmax_ maximum carboxylation rate of phosphoenolpyruvate carboxylase; *V*_max_, *c*_i_-saturated rate of photosynthesis. Values are means ± SEM (*n* = 4-5 biological replicates). *p*-values are from one-way ANOVA testing the effect of temperature. Different letters denote significant differences in the same parameter (Tukey HSD).

**Supplemental Table S2.** Slopes and intercepts from linear regression analysis of metabolite contents in fractionated leaf tissue obtained from serial filtration over liquid nitrogen to estimate proportion of pyruvate, PEP, 3PGA and DHAP in bundle sheath or mesophyll cells. Leaf tissue was sampled during

gas exchange at three different temperatures: 7, 15 and 25 ^o^C. Maize plants were acclimated for one

hour at the desired temperature (25, 15 or 7°C) with the lights in the cabinet turned on to a photosynthetic active radiation (*Q*) of 600 μmol m^-2^ s^-1^. Afterwards, the youngest completely expanded leaf was clamped into a 9 cm^2^ leaf chamber at 1500 µmol m^-2^ s^-1^ actinic red light, and samples were snap-frozen inside the gas exchange instrument after 30 min when steady state had been reached. n = 5-6 experimental replicates. Each replicate consisted of pooled samples from 11 plants. Final estimates

are shown in Fig. 8B and C in main text. Linear regressions for each of the metabolites are shown in Supplemental Figures S14-S17.

| Metabolite proportion in bundle sheath cells | | | | |
| --- | --- | --- | --- | --- |
| Metabolite | Temperature | Intercept from %XX/%PRK against %PEPC/%PRK | Slope from %XX/%PEPC against %PRK/%PEPC | Proportion based on average of intercept and slope |
| Pyruvate | 7°C | 0.408 ± 0.200 | 0.392 ± 0.154 | 0.400 ± 0.126 |
|  | 15°C | 0.509 ± 0.115 | 0.558 ± 0.076 | 0.534 ± 0.069 |
|  | 25°C | 0.742 ± 0.109 | 0.699 ± 0.103 | 0.720 ± 0.075 |
| PEP | 7°C | 0.748 ±0.219 | 0.519 ± 0.241 | 0.633 ± 0.163 |
|  | 15°C | 0.249 ± 0.122 | 0.339 ± 0.083 | 0.294 ± 0.074 |
|  | 25°C | 0.524 ± 0.115 | 0.484 ± 0.096 | 0.504 ± 0.075 |
| 3PGA | 7°C | 1.056 ± 0.172 | 1.042 ± 0.177 | 1.049 ± 0.123 |
|  | 15°C | 1.214 ± 0.205 | 0.964 ± 0.164 | 1.089 ± 0.131 |
|  | 25°C | 0.871 ± 0.142 | 0.912 ± 0.141 | 0.891 ± 0.100 |
| DHAP | 7°C | 0.620 ± 0.250 | 0.386 ± 0.098 | 0.503 ± 0.134 |
|  | 15°C | 0.294 ± 0.077 | 0.383 ± 0.064 | 0.338 ± 0.050 |
|  | 25°C | 0.697 ± 0.148 | 0.527 ± 0.134 | 0.612 ± 0.100 |
| Metabolite proportion in mesophyll cells | | | | |
| Metabolite | Temperature | Intercept from %XX/%PEPC against %PRK/%PEPC | Slope from %XX/%PRK against %PEPC/%PRK | Proportion based on average of intercept and slope |
| Pyruvate | 7°C | 0.619 ± 0.204 | 0.598 ± 0.158 | 0.609 ± 0.129 |
|  | 15°C | 0.385 ± 0.099 | 0.440 ± 0.106 | 0.412 ± 0.073 |
|  | 25°C | 0.242 ± 0.146 | 0.201 ± 0.084 | 0.222 ± 0.084 |
| PEP | 7°C | 0.566 ± 0.327 | 0.320 ± 0.179 | 0.443 ± 0.186 |
|  | 15°C | 0.629 ± 0.108 | 0.728 ± 0.113 | 0.678 ± 0.078 |
|  | 25°C | 0.479 ± 0.137 | 0.435 ± 0.088 | 0.457 ± 0.081 |
| 3PGA | 7°C | -0.124 ± 0.254 | -0.145 ± 0.134 | -0.134 ± 0.144 |
|  | 15°C | -0.037 ± 0.218 | -0.302 ± 0.174 | -0.169 ± 0.139 |
|  | 25°C | -0.011 ± 0.200 | 0.027 ± 0.109 | 0.008 ± 0.114 |
| DHAP | 7°C | 0.618 ± 0.143 | 0.515 ± 0.197 | 0.567 ± 0.122 |
|  | 15°C | 0.595 ± 0.084 | 0.691 ± 0.065 | 0.643 ± 0.053 |
|  | 25°C | 0.473 ± 0.190 | 0.298 ± 0.113 | 0.385 ± 0.111 |

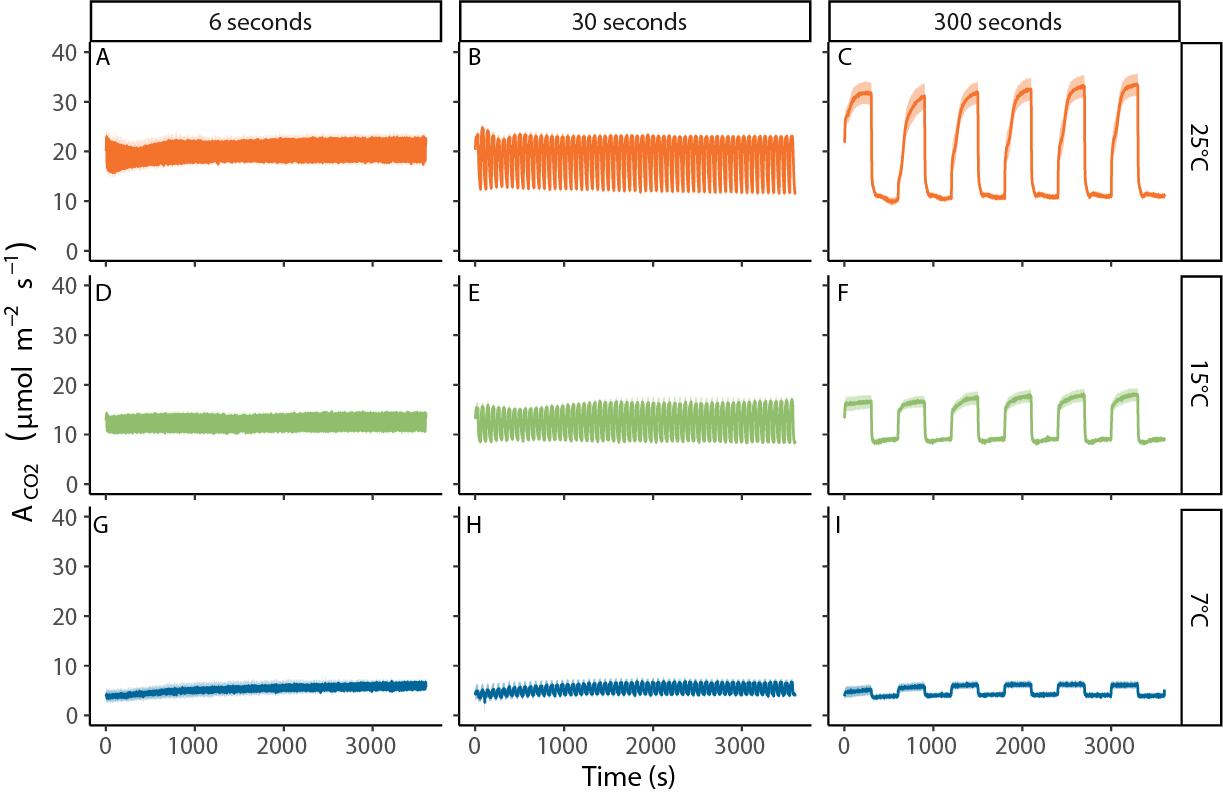

**Supplemental Figure S1.** Net CO_2_ assimilation (*A*_CO2_) in maize plants measured under three different fluctuating light regimes, at 25 °C (A, B, and C), 15 °C (D, E, and F), or 7 °C (G, H, and I). In each fluctuating light regime, leaves were exposed to repetitive changes between low (200 µmol m^-2^ s^-1^) and high (1500 µmol m^-2^ s^-1^) light-steps with duration of either 6 s (FL6; A, D, G), 30 s (FL30; B,E, H) or 300 s (FL300; C, F, I). Measurements were performed on maize plants acclimated at 7, 15 or 25°C for at least 2 h. Fluctuating light regimes were started after leaves were acclimated to steady state at light intensity of 600 µmol m^-2^ s^-1^ and lasted 1 hour. Ribbons represent standard error of the mean (n=4-5).

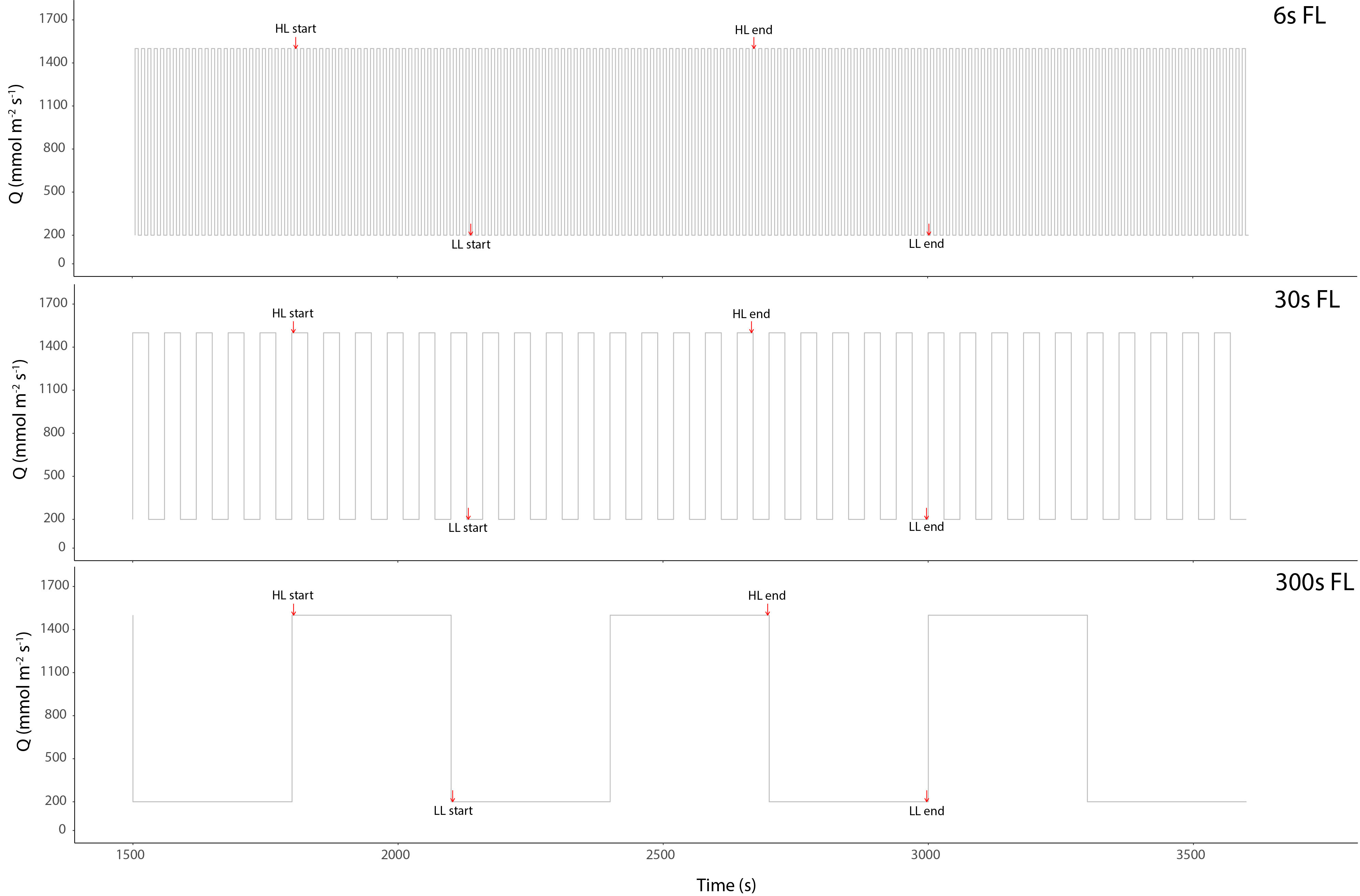

**Supplemental Figure S2.** Scheme showing timing of chlorophyll fluorescence and P700 redox changes measurements in maize plants under three different fluctuating light regimes. Each light regime consisted of alternating 1500 and 200 µmol m^-2^ s^-1^ photosynthetic active radiation (*Q*) steps, where each light step lasted 6, 30, or 300 seconds. Chlorophyl fluorescence and P700 redox changes measurements were taken in the second half of each experimental timeseries (from 1500 s onwards), when gas exchange responses to repeated fluctuations had become highly similar (see Supp. Figure S1) and were spaced out by at least 5 min to avoid impacting the dynamic responses of photosynthesis. Measurements were timed either 3 s after a change in light intensity (referred to as ‘Start’ in Figures 4-6) or 3 s before a change in light intensity (referred to as ‘End’ in Figures 4-6). Specific timings for measurements were as follows. For FL300: 1803 s (HL ‘Start’), 2103 s (LL ‘Start’), 2697 s (HL ‘End’), 2997 s (LL ‘End’). For FL30 and FL6: 1803 s (HL ‘Start), 2133 s (LL ’Start’), 2667 s (HL ‘End’), and 2997 s (LL ‘End’).

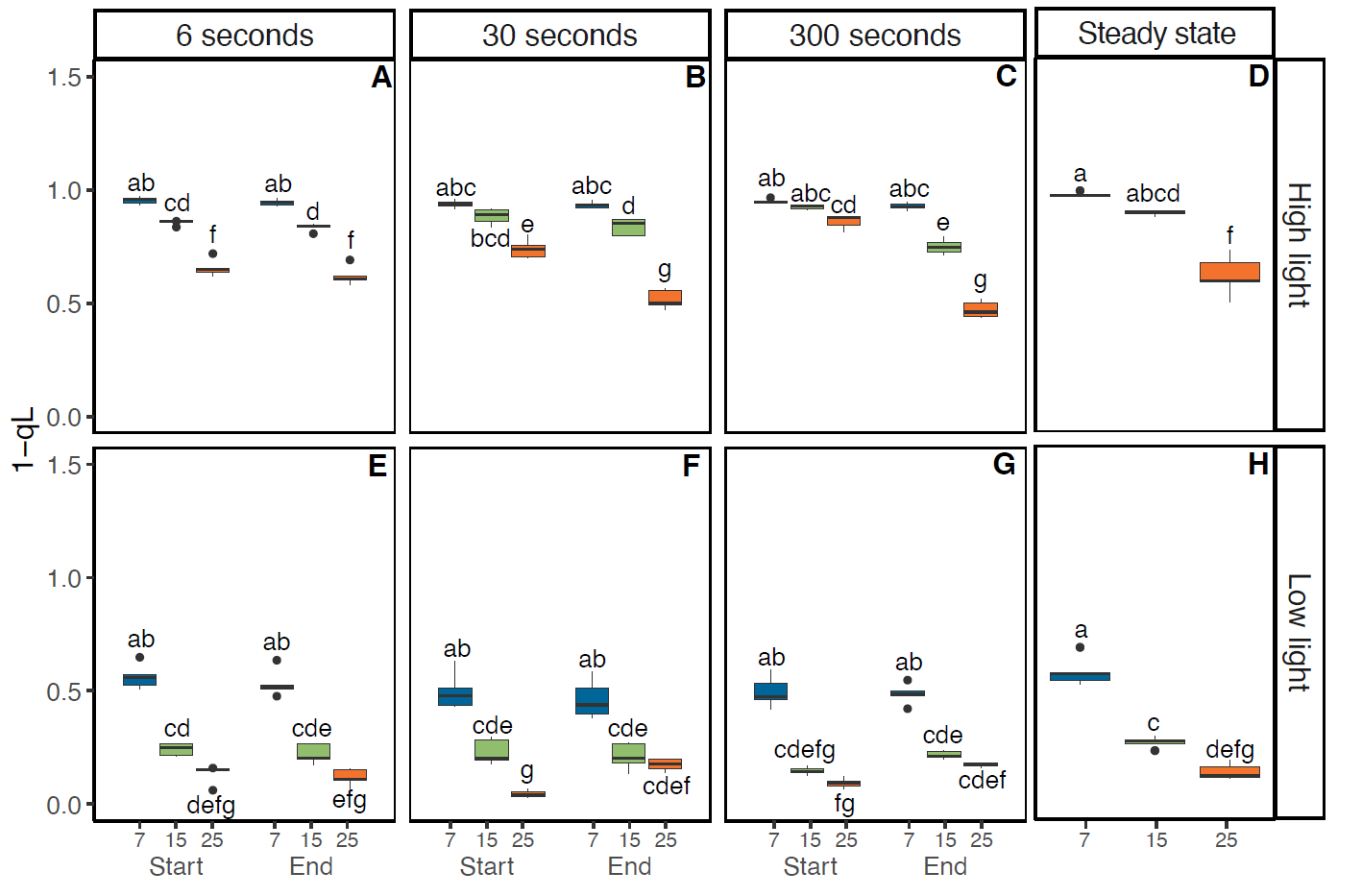

**Supplemental Figure S3.** Quinone A redox state in maize leaves as a function of temperature and fluctuating light regime. Quinone A redox state was estimated from chlorophyll fluorescence measurements as 1-*q*L based on Kramer et al. (2004). Measurements were performed at 25 °C, 15 °C, or 7 °C and three fluctuating light regimes. Maize plants were acclimated at 7, 15 or 25°C for at least 2 h prior to measurements. In each fluctuating light regime, leaves were exposed to repetitive changes between low (200 µmol m^-2^ s^-1^) and high (1500 µmol m^-2^ s^-1^) light-steps with duration of either 6, 30 or 300 s (FL6, FL30, FL300). In each fluctuating light regime, 1-qL was determined 3 s into a new light intensity (‘Start’) and 3 s before switching (‘End’). Thus, a total of four measurements were taken for each biological replicate, two measurements (‘Start’ and ‘End’) during high light (A-C) and two measurements during low light (E-G). Precise timings of measurements in each fluctuating light regime are provided in Supplemental Fig. S2. Measurements at steady state at matching light intensities and temperatures were obtained from light response curves shown in Fig 1A and included for comparison (D, H). Box edges represent the lower and upper quartiles, the solid line indicates the median, and points represent outliers beyond 1.5 times the interquartile range (*n* = 4-5 biological replicates). Statistical analyses were run on Box-Cox transformed data. Three-way ANOVA was used to test the effect of temperature, fluctuation length, and measurement time. Different letters indicate statistical differences according to Tukey test (*p*<0.05)*.*

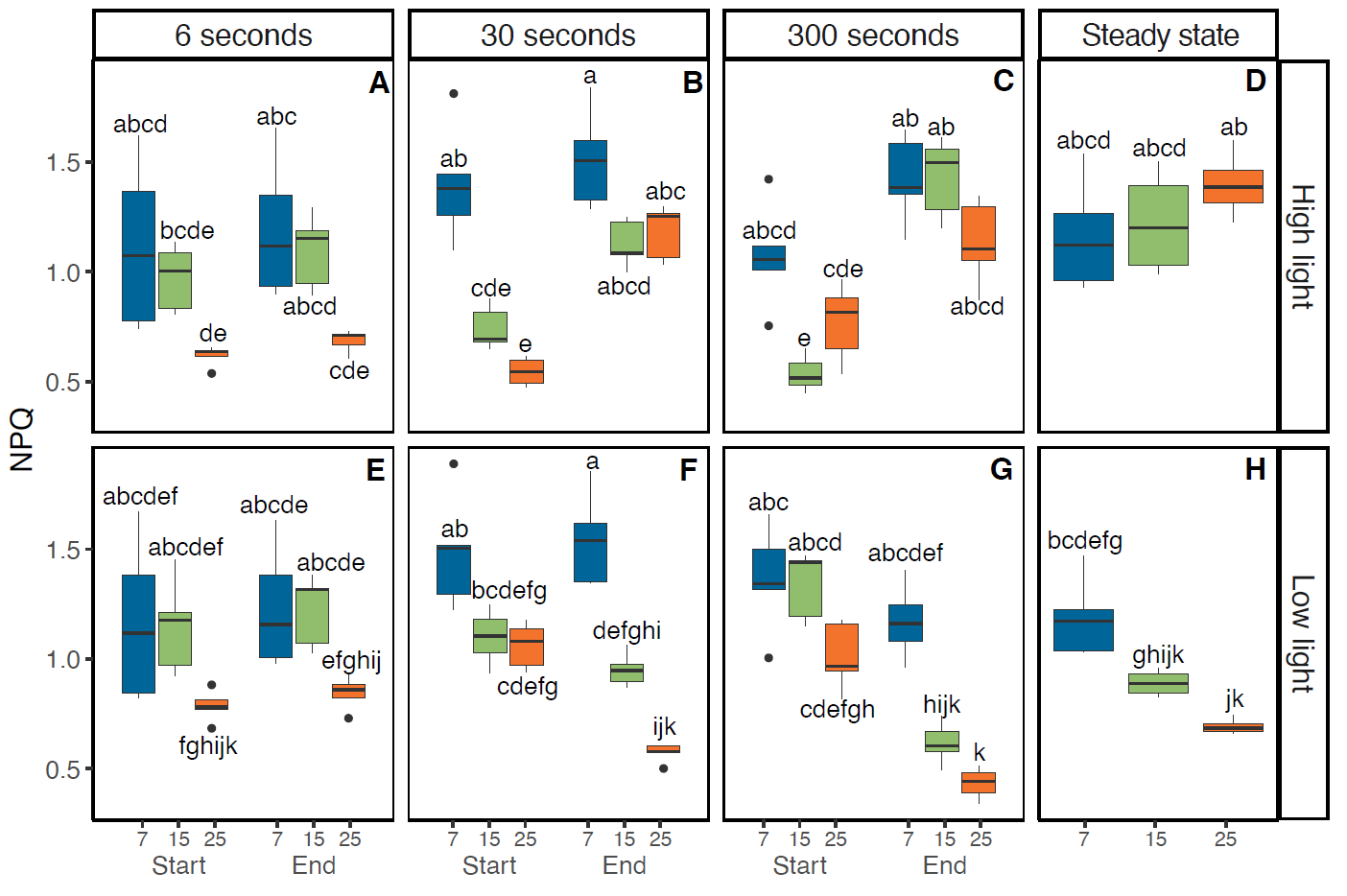

**Supplemental Figure S4.** Non-photochemical quenching (NPQ) in maize leaves as a function of temperature and fluctuating light regime. NPQ was estimated from chlorophyll fluorescence measurements based on Stern-Volmer quenching *F*_m_/*F*_m_’ - 1. Measurements were performed at 25 °C, 15 °C, or 7 °C and three fluctuating light regimes. Maize plants were acclimated at 7, 15 or 25°C for at least 2 h prior to measurements. In each fluctuating light regime, leaves were exposed to repetitive changes between low (200 µmol m^-2^ s^-1^) and high (1500 µmol m^-2^ s^-1^) light-steps with duration of either 6, 30 or 300 s (FL6, FL30, FL300). In each fluctuating light regime, NPQ was determined 3 s into a new light intensity (‘Start’) and 3 s before switching (‘End’). Thus, a total of four measurements were taken for each biological replicate, two measurements (‘Start’ and ‘End’) during high light (A-C) and two measurements during low light (E-G). Precise timings of measurements in each fluctuating light regime are provided in Supplemental Fig. S2. Measurements at steady state at matching light intensities and temperatures were obtained from light response curves shown in Fig 1A and included for comparison (D, H). Box edges represent the lower and upper quartiles, the solid line indicates the median, and points represent outliers beyond 1.5 times the interquartile range (*n* = 4-5 biological replicates). Three-way ANOVA was used to test the effect of temperature, fluctuation length, and measurement time. Different letters indicate statistical differences according to Tukey test (*p*<0.05)*.*

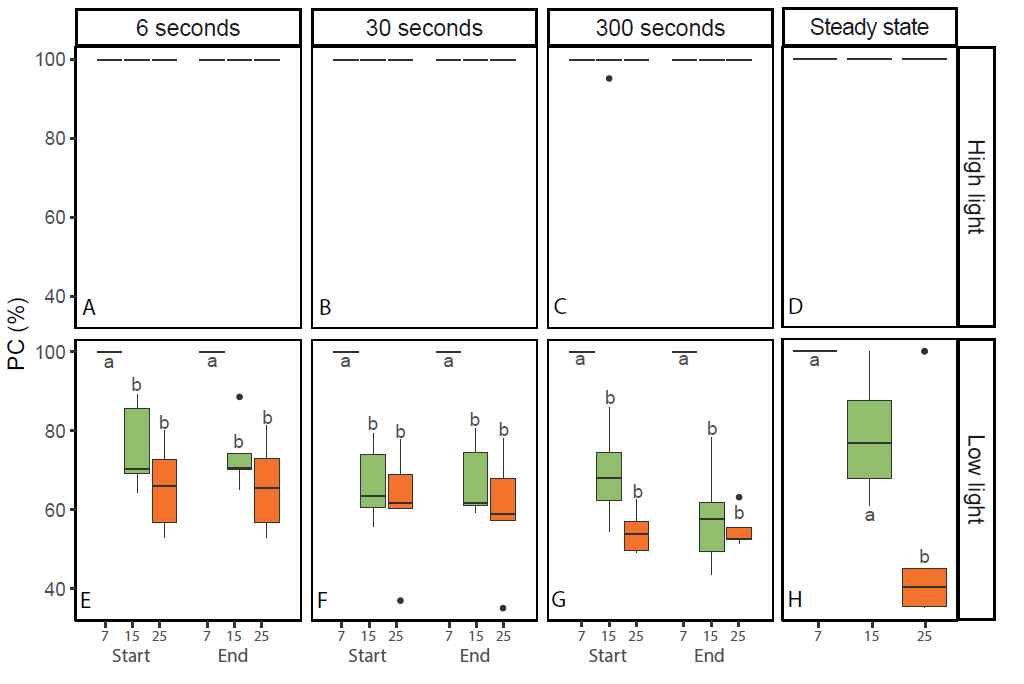

**Supplemental Figure S5.** Plastocyanin redox state in maize leaves as a function of temperature and fluctuating light regime. Plastocyanin redox state (%) was estimated from near infrared differential absorption measurements (100% represents fully oxidized). Measurements were performed at 25 °C, 15 °C, or 7 °C and three fluctuating light regimes. Maize plants were acclimated at 7, 15 or 25°C for at least 2 h prior to measurements. In each fluctuating light regime, leaves were exposed to repetitive changes between low (200 µmol m^-2^ s^-1^) and high (1500 µmol m^-2^ s^-1^) light-steps with duration of either 6, 30 or 300 s (FL6, FL30, FL300). In each fluctuating light regime, Plastocyanin redox state was determined 3 s into a new light intensity (‘Start’) and 3 s before switching (‘End’). Thus, a total of four measurements were taken for each biological replicate, two measurements (‘Start’ and ‘End’) during high light (A-C) and two measurements during low light (E-G). Precise timings of measurements in each fluctuating light regime are provided in Supplemental Fig. S2. Measurements at steady state at matching light intensities and temperatures were obtained from light response curves shown in Fig 1A and included for comparison (D, H). Box edges represent the lower and upper quartiles, the solid line indicates the median, and points represent outliers beyond 1.5 times the interquartile range (*n* = 4-5 biological replicates). Different letters indicate statistical differences within each FL regime by light intensity combination according to Kruskal-Wallis test followed by post hoc Dunn test (*p*<0.05)*.*

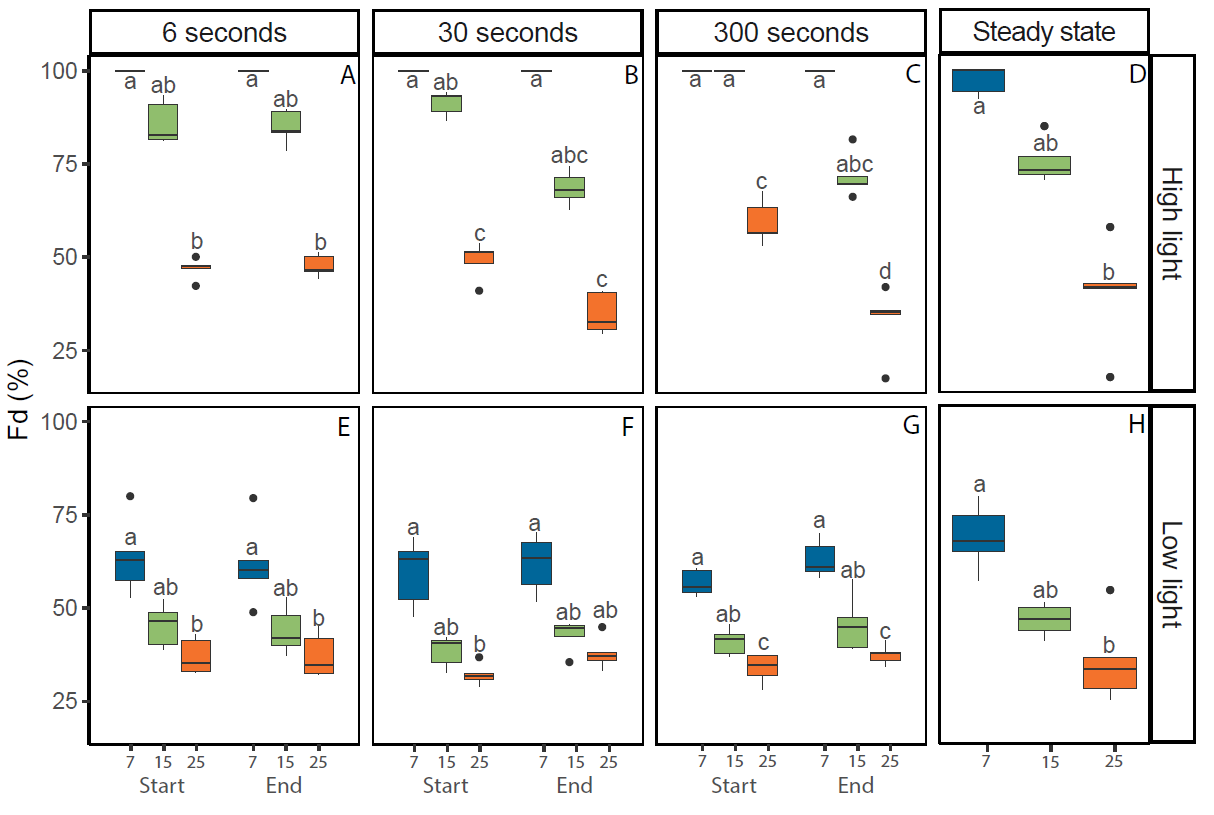

**Supplemental Figure S6.** Ferredoxin redox state in maize leaves as a function of temperature and fluctuating light regime. Ferredoxin redox state (%) was estimated from near infrared differential absorption measurements (100% represents fully reduced). Measurements were performed at 25 °C, 15 °C, or 7 °C and three fluctuating light regimes. Maize plants were acclimated at 7, 15 or 25°C for at least 2 h prior to measurements. In each fluctuating light regime, leaves were exposed to repetitive changes between low (200 µmol m^-2^ s^-1^) and high (1500 µmol m^-2^ s^-1^) light-steps with duration of either 6, 30 or 300 s (FL6, FL30, FL300). In each fluctuating light regime, Ferredoxin redox state was determined 3 s into a new light intensity (‘Start’) and 3 s before switching (‘End’). Thus, a total of four measurements were taken for each biological replicate, two measurements (‘Start’ and ‘End’) during high light (A-C) and two measurements during low light (E-G). Precise timings of measurements in each fluctuating light regime are provided in Supplemental Fig. S2. Measurements at steady state at matching light intensities and temperatures were obtained from light response curves shown in Fig 1A and included for comparison (D, H). Box edges represent the lower and upper quartiles, the solid line indicates the median, and points represent outliers beyond 1.5 times the interquartile range (*n* = 4-5 biological replicates). Different letters indicate statistical differences within each FL regime by light intensity combination according to Kruskal-Wallis test followed by post hoc Dunn test (*p*<0.05)*.*

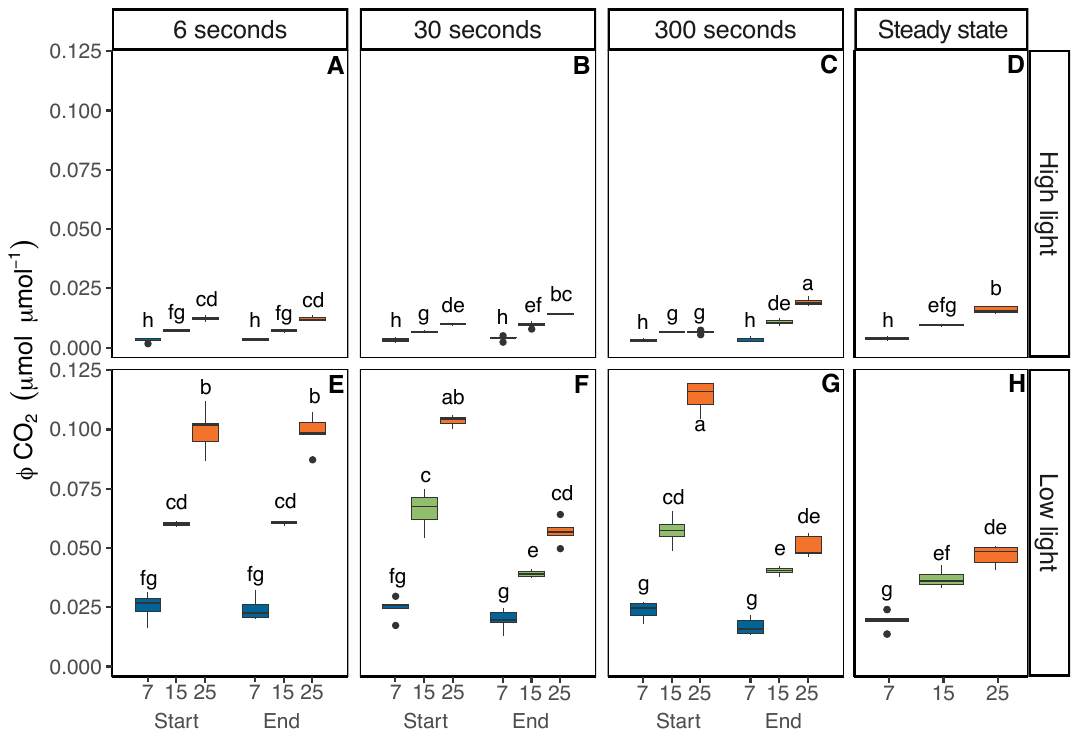

**Supplemental Figure S7.** Instantaneous quantum yield of CO_2_ fixation (φ_CO2_) in maize leaves as a function of temperature and fluctuating light regime. Measurements were performed at 25 °C, 15 °C, or 7 °C and three fluctuating light regimes. Maize plants were acclimated at 7, 15 or 25°C for at least 2 h prior to measurements. In each fluctuating light regime, leaves were exposed to repetitive changes between low (200 µmol m^-2^ s^-1^) and high (1500 µmol m^-2^ s^-1^) light-steps with duration of either 6, 30 or 300 s (FL6, FL30, FL300). In each fluctuating light regime, φ_CO2_ was determined 3 s into a new light intensity (‘Start’) and 3 s before switching (‘End’) to match φ_PSII_ and φ_PSI_ observations shown in Figs 4 and 5. Thus, a total of four measurements were taken for each biological replicate, two measurements (‘Start’ and ‘End’) during high light (A-C) and two measurements during low light (E-G). Precise timings of measurements in each fluctuating light regime are provided in Supplemental Fig. S2. Measurements of φ_CO2_ at steady state at matching light intensities and temperatures were obtained from light response curves shown in Fig 1A and included for comparison (D, H). Box edges represent the lower and upper quartiles, the solid line indicates the median, and points represent outliers beyond 1.5 times the interquartile range (n = 4-5 biological replicates). Statistical analyses were run on Box-Cox transformed data. Three-way ANOVA was used to test the effect of temperature, fluctuation length, and flash time. Different letters indicate statistical differences according to Tukey test (*p*<0.005)*.*

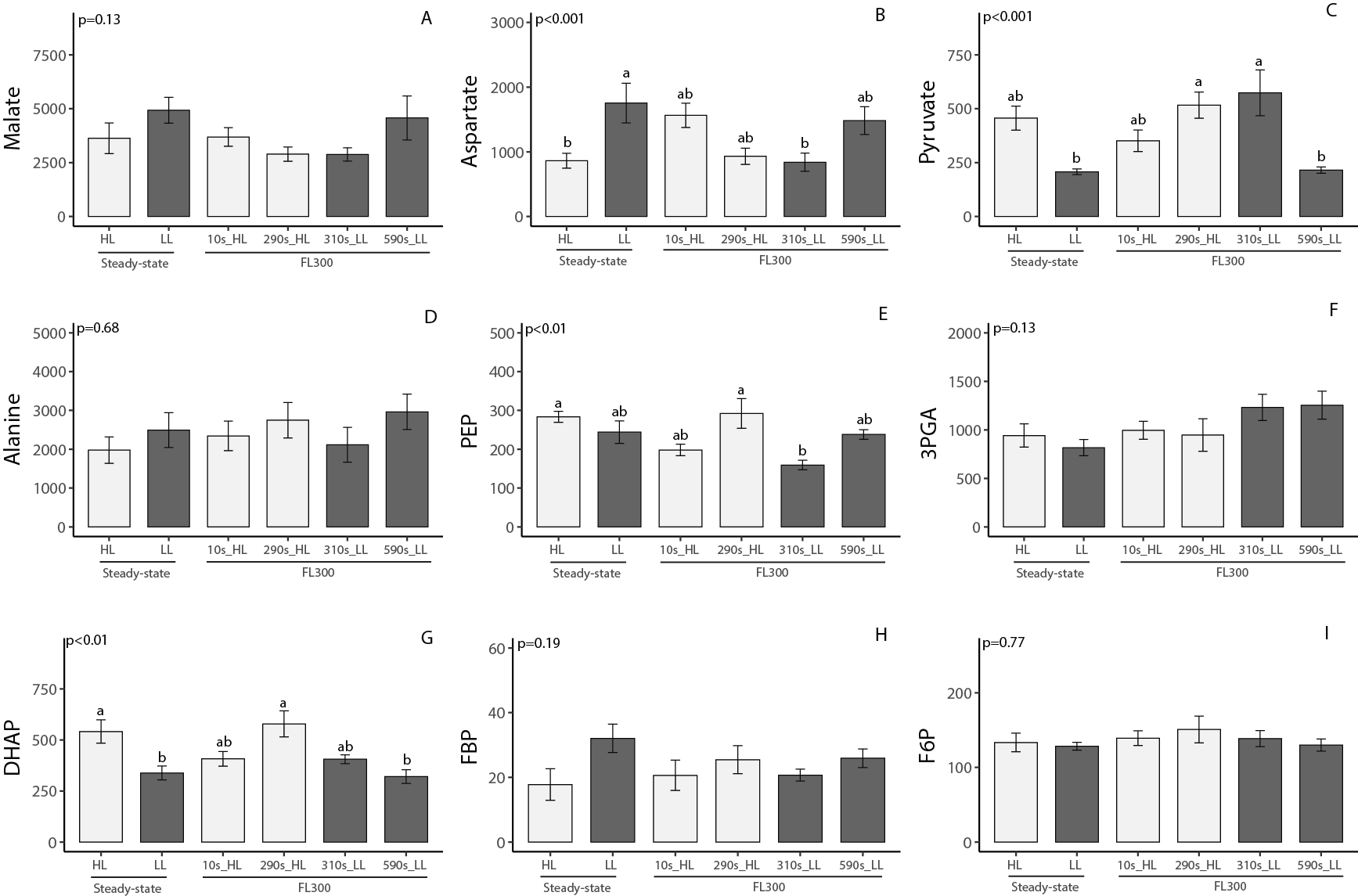

*cont.*

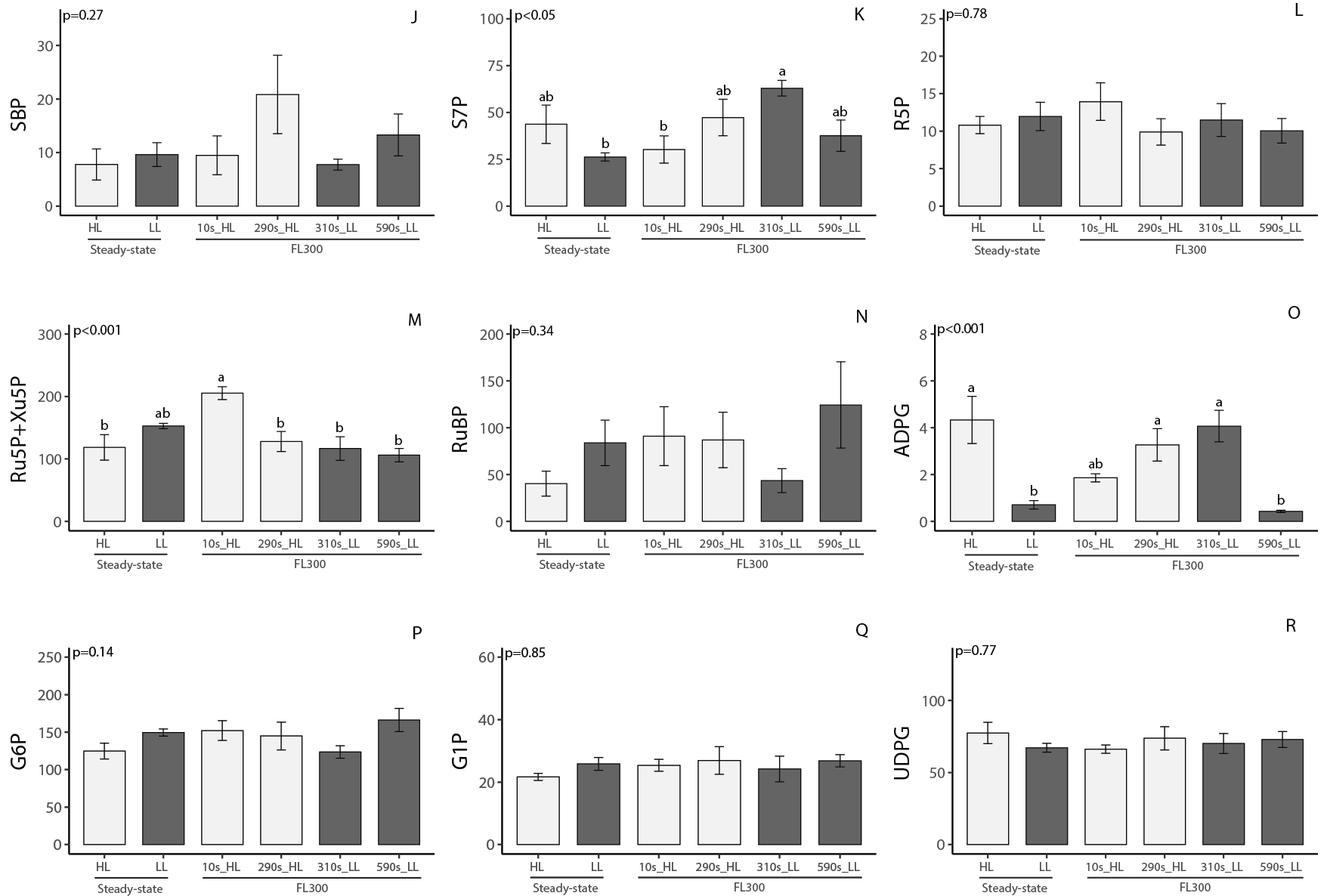

*cont.*

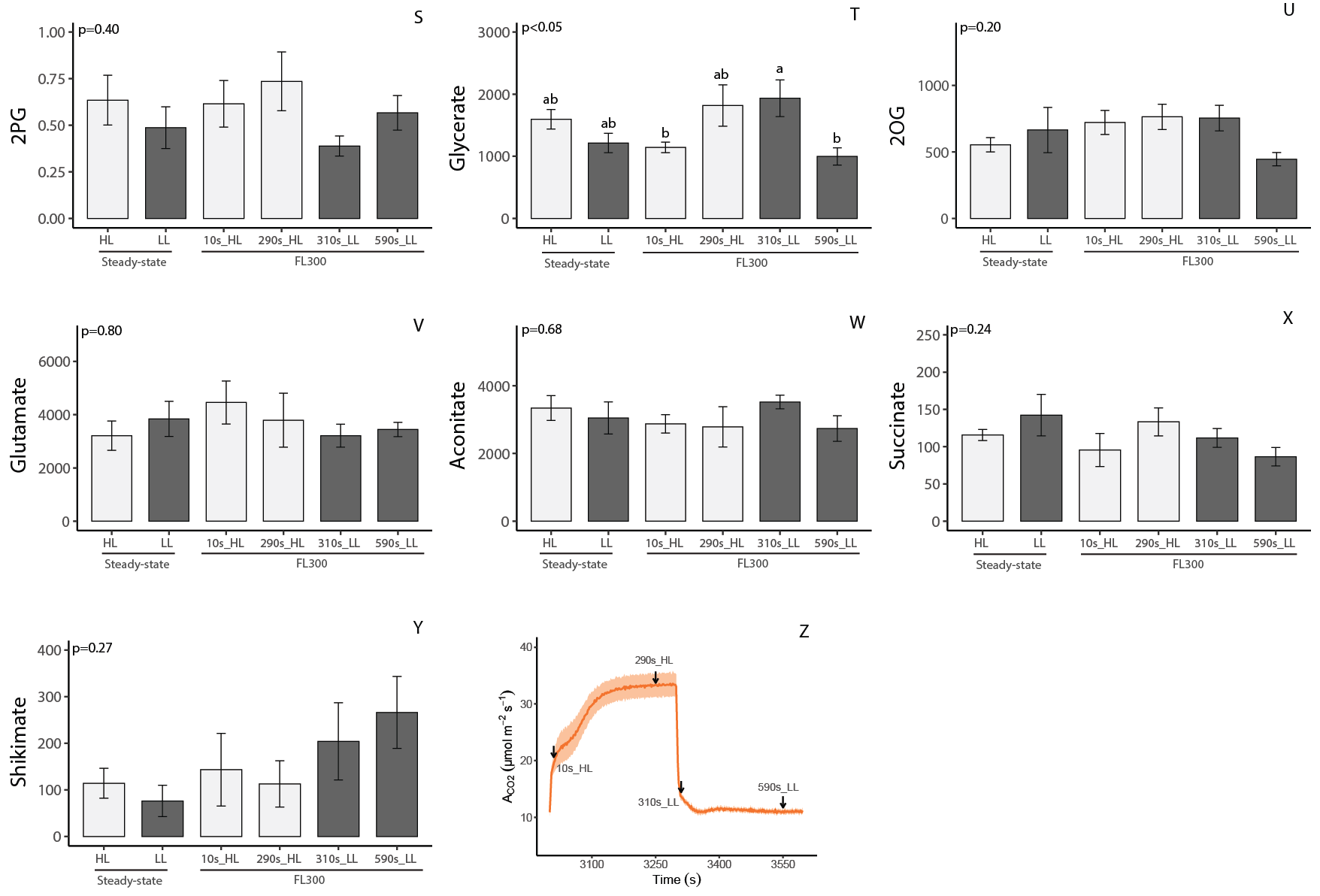

**Supplemental Figure S8.** Metabolite profiles of maize leaves exposed to constant or fluctuating light. For constant light, leaves were sampled following exposure with low (200 µmol m^-2^ s^-1^) and high (1500 µmol m^-2^ s^-1^) light intensity. For the fluctuating light regime, leaves were exposed to repetitive changes between low and high light-steps with duration of 300 s (FL300). Samples were taken at four different time points: 10 s after switching from low to high light (FL300_10s_HL), 290 s after switching from low to high light (FL300_290s_HL), 10 s after switching from high to low light (FL300_310s_LL), and 290 s after switching from high to low light (FL300_590s_LL). Plants were acclimated to 25 °C for at least 2h prior to sampling. Sampling timepoints relative to gas exchange timeseries are indicated in panels XX. Metabolite contents are shown in nmol g FW^−1^. Values are means ± SEM (n = 5-6 biological replicates). Data were analyzed using ANOVA to test the effect of sampling time, significance is indicated by *p*-values inset. Where a significant sampling time effect was found by ANOVA, different letters indicate statistical differences according to Tukey test (*α*<0.05).

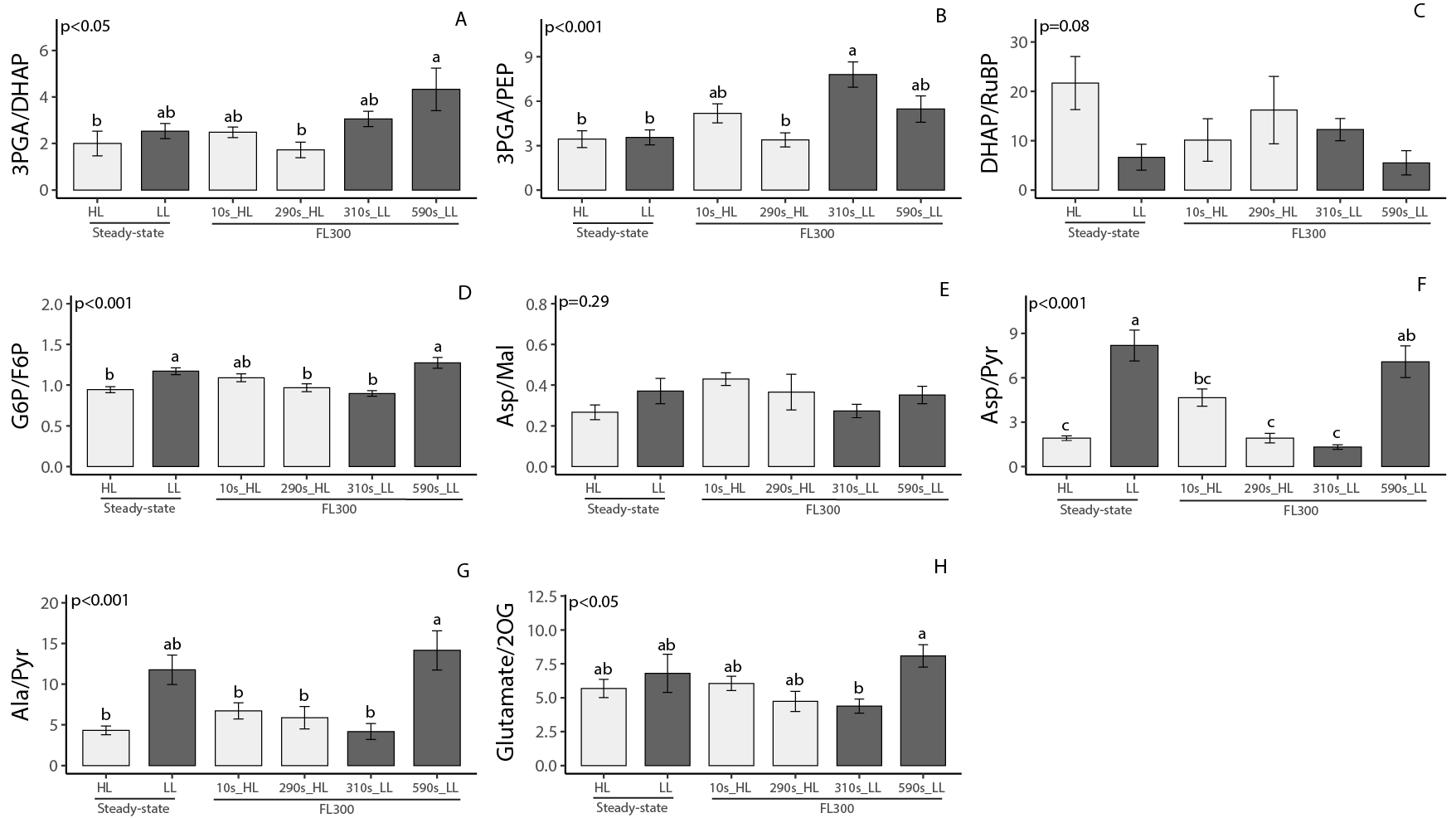

**Supplemental Figure S9.** Selected ratios between metabolite profiles of maize leaves exposed to constant or fluctuating light shown in Supplemental Figure S13. For constant light, leaves were sampled following exposure with low (200 µmol m^-2^ s^-1^) and high (1500 µmol m^-2^ s^-1^) light intensity. For the fluctuating light regime, leaves were exposed to repetitive changes between low and high light-steps with duration of 300 s (FL300). Samples were taken at four different time points: 10 s after switching from low to high light (FL300_10s_HL), 290 s after switching from low to high light (FL300_290s_HL), 10 s after switching from high to low light (FL300_310s_LL), and 290 s after switching from high to low light (FL300_590s_LL). Plants were acclimated to 25 °C for at least 2h prior to sampling. Values are means ± SEM (n = 5-6 biological replicates). Data were analyzed using ANOVA to test the effect of sampling time, significance is indicated by inset *p*-values. Where a significant sampling time effect was found by ANOVA, different letters indicate statistical differences according to Tukey test (*α*<0.05)

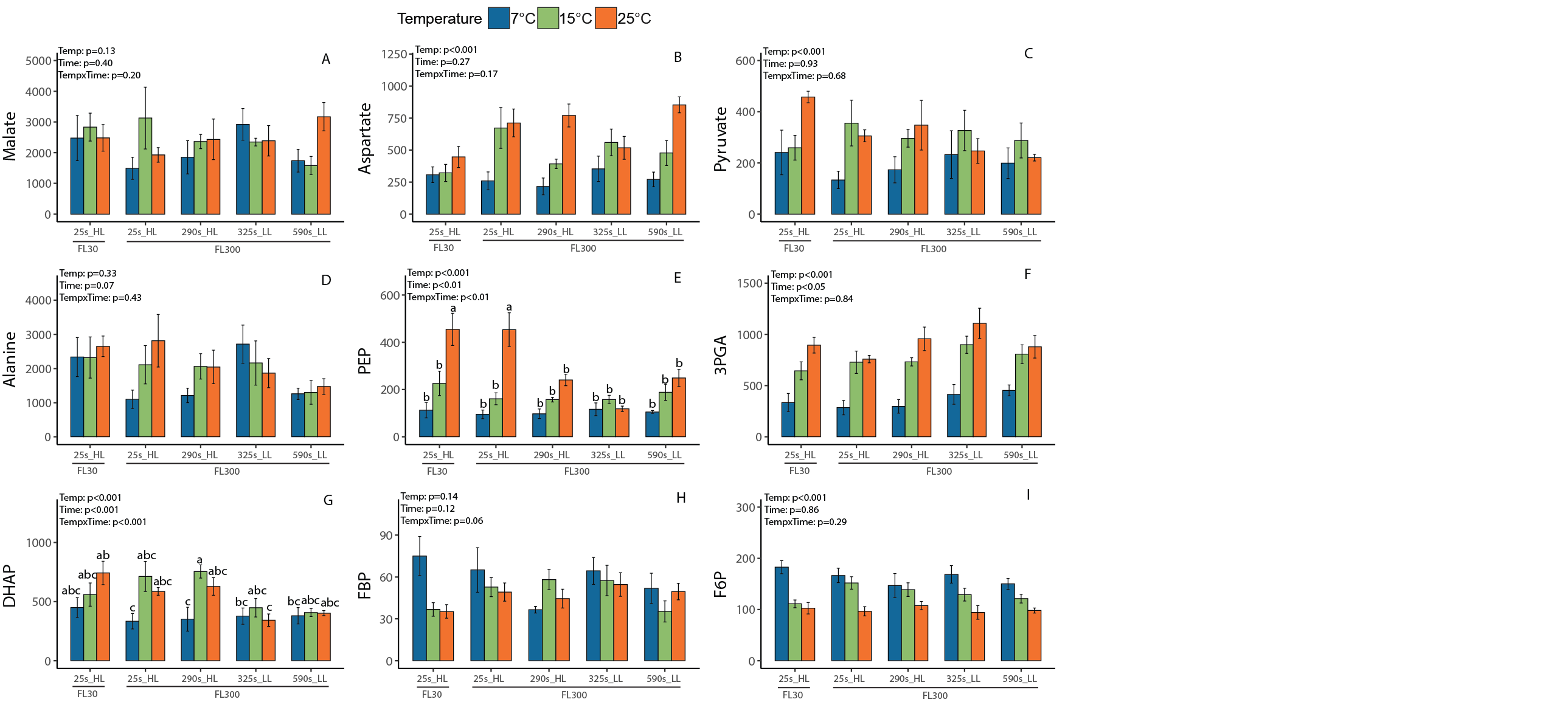

*cont.*

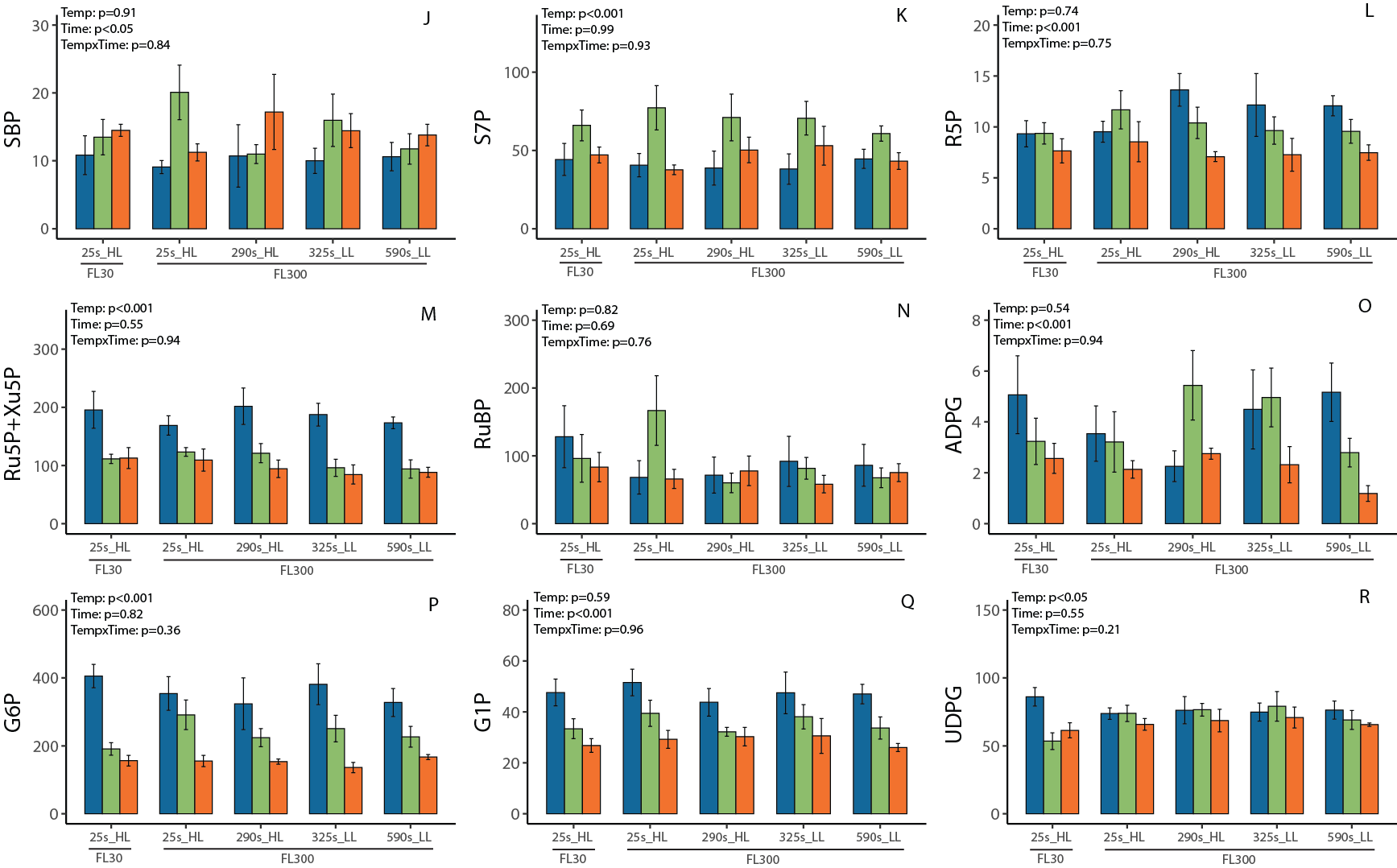

*cont.*

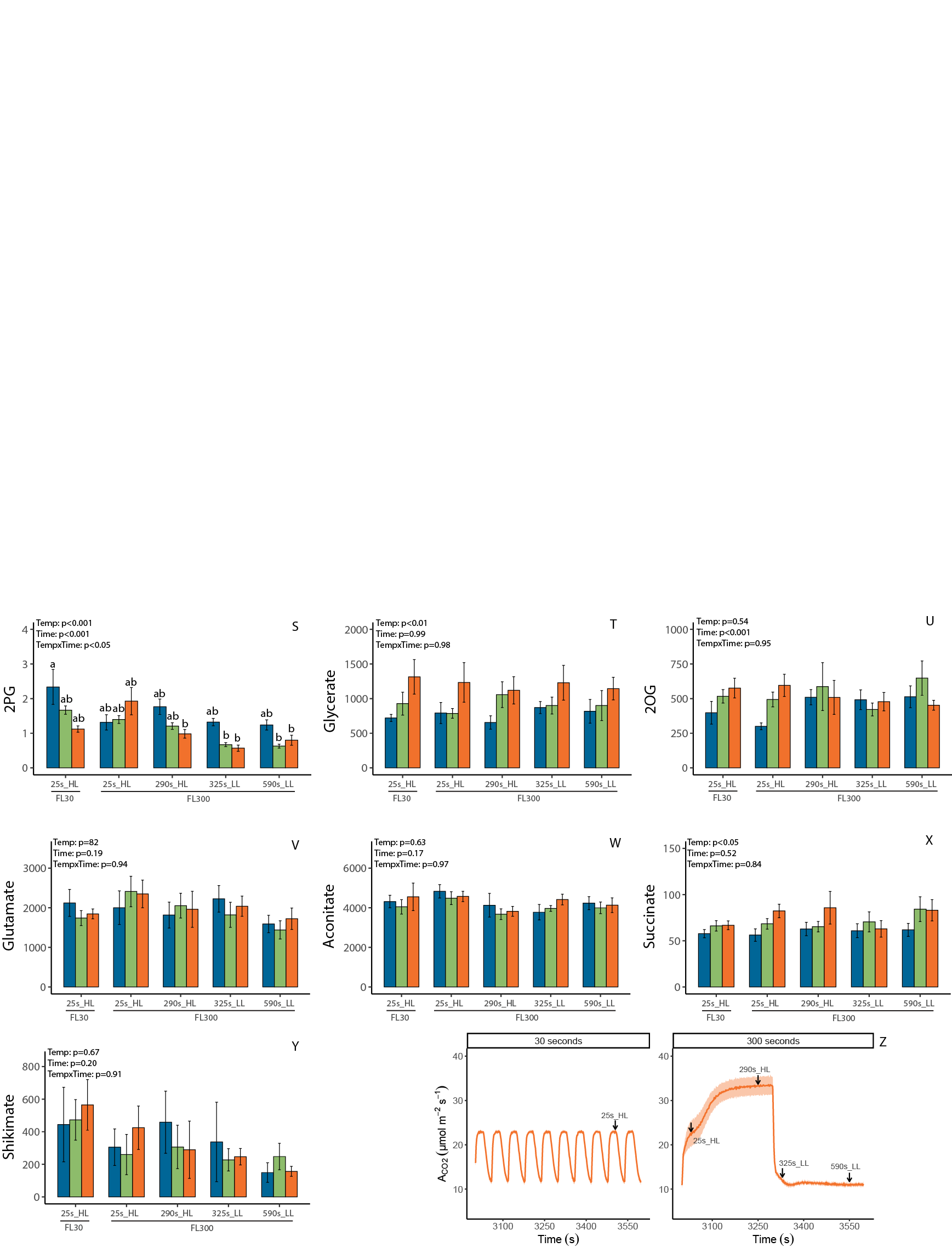

**Supplemental Figure S10.** Metabolite profiles of maize leaves exposed to fluctuating light regimes at three different temperatures. In each fluctuating light regime, leaves were exposed to repetitive changes between low (200 µmol m^-2^ s^-1^) and high (1500 µmol m^-2^ s^-1^) light-steps with duration of either 30 or 300 s (FL30, FL300). For the FL30 light regime, samples were taken 25 s after switching from low to high light (FL30_25s_HL). For the FL300 light regime, samples were taken at four different time points: 25 s after switching from low to high light (FL300_25s_HL), 290 s after light switches from low to high light phase (FL300_290s_HL), 25 s after switching from high to low light (FL300_325s_LL), and 290 s after switching from high to low light (FL300_590s_LL). Sampling timepoints relative to gas exchange timeseries are indicated in panel Z. Plants were acclimated to 25 °C, 15 °C, or 7 °C for at least 2h prior to sampling. Metabolite contents are shown in nmol g FW^−1^. Values are means ± SEM (n = 5-6 biological replicates). Data were analyzed using two-way ANOVA to test for effects of temperature (Temp), sampling time (Time), and their interaction (TempxTime). Significance of ANOVA effects is indicated by p-value insets. In case of significant TempxTime interaction, different letters indicate statistical differences according to Tukey test (*p*<0.05).

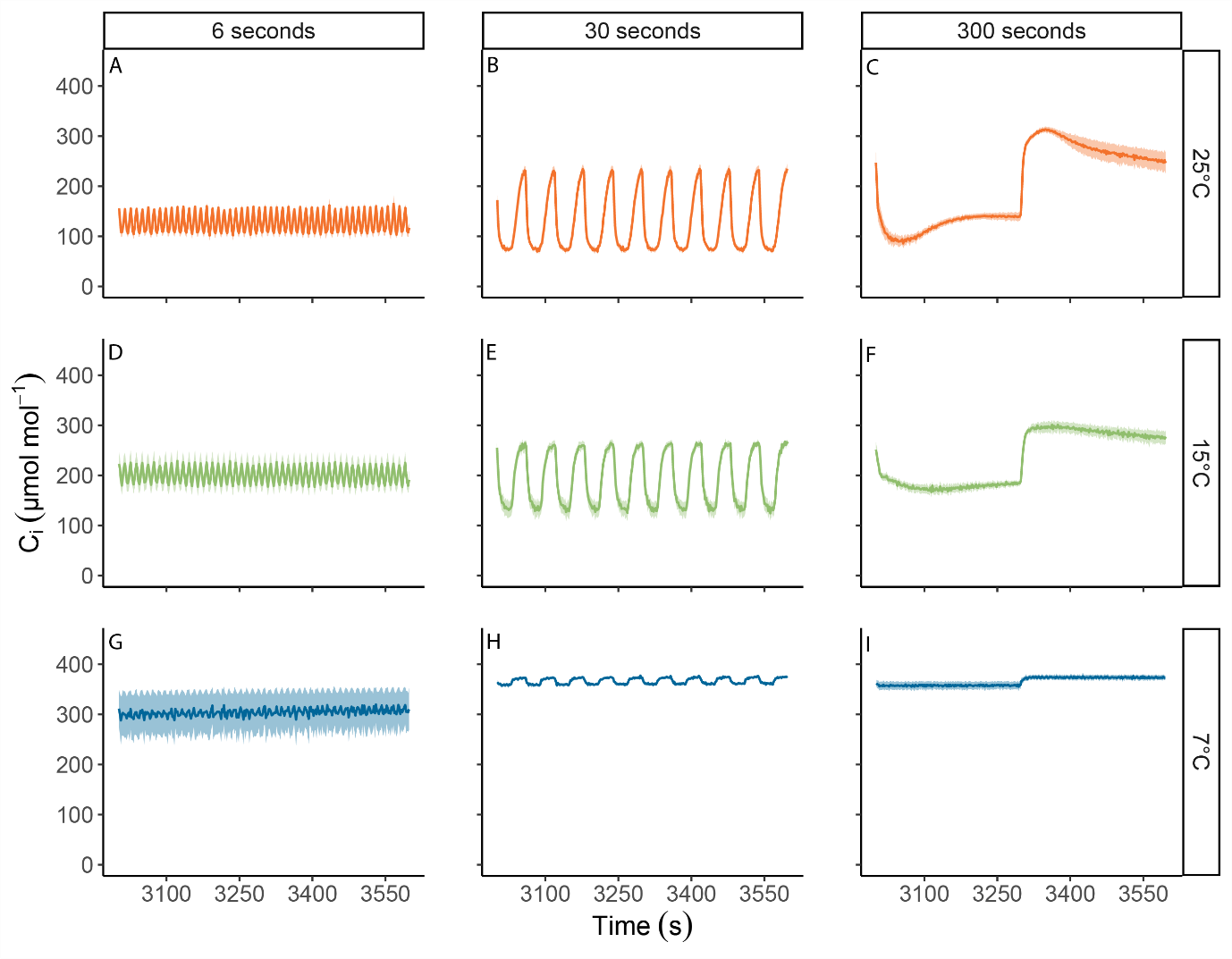

**Supplemental Figure S11.** Intercellular CO_2_ concentration (*c*_i_) in maize plants measured under three different fluctuating light regimes, at 25 °C (A, B, and C), 15 °C (D, E, and F), or 7 °C (G, H, and I). In each fluctuating light regime, leaves were exposed to repetitive changes between low (200 µmol m^-2^ s^-1^) and high (1500 µmol m^-2^ s^-1^) light-steps with duration of either 6 s (FL6; A, D, G), 30 s (FL30; B,E, H) or 300 s (FL300; C, F, I). Measurements were performed on maize plants acclimated at 7, 15 or 25°C for at least 2 h. Fluctuating light regimes were started after leaves were acclimated to steady state at light intensity of 600 µmol m^-2^ s^-1^ and lasted 1 hour. Data are shown from the final ten minutes of each light regime. Ribbons represent standard error of the mean (*n*=4-5).

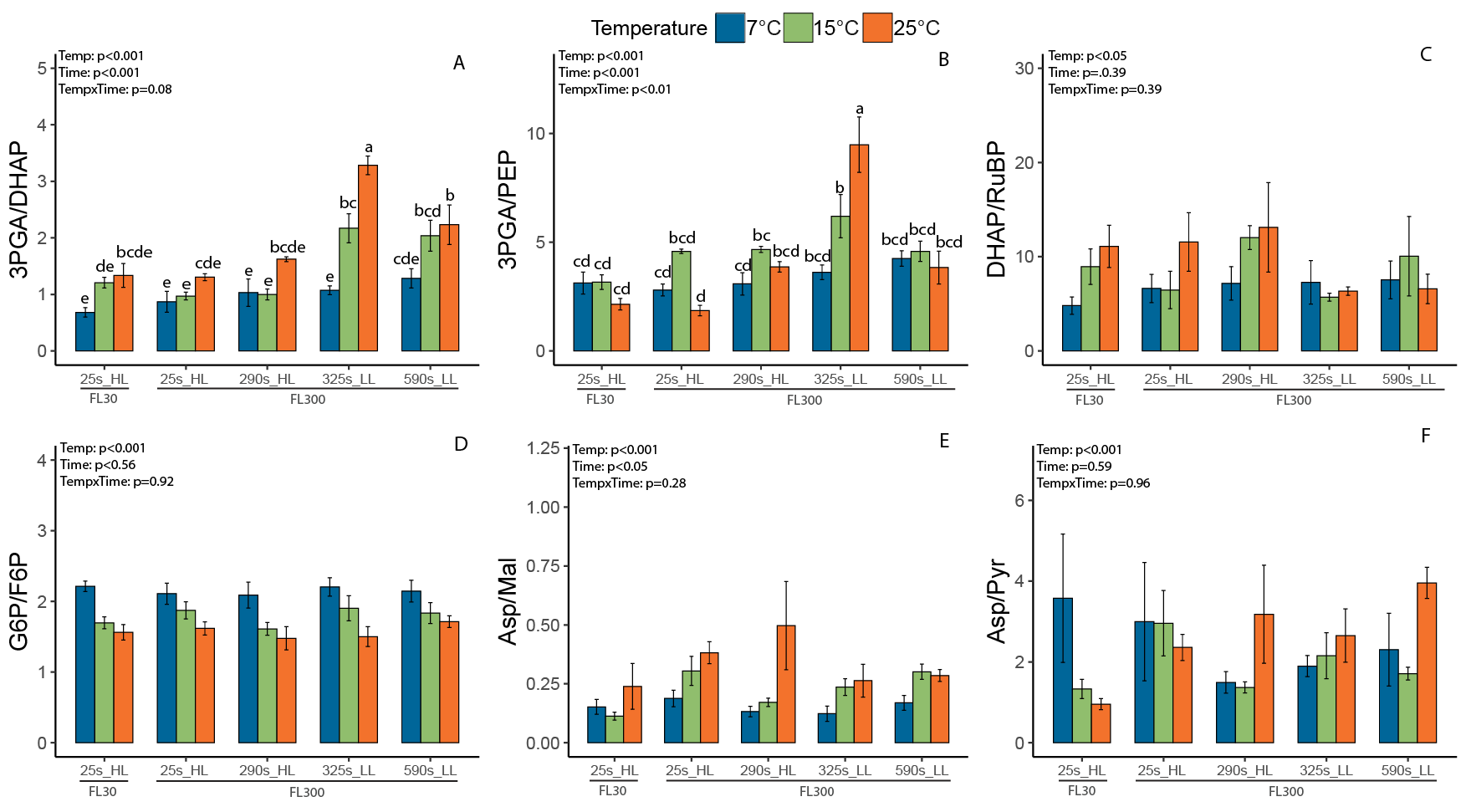

**Supplemental Figure S12.** Selected ratios between metabolite profiles of maize leaves exposed to fluctuating light regimes at three different temperatures. In each fluctuating light regime, leaves were exposed to repetitive changes between low (200 µmol m^-2^ s^-1^) and high (1500 µmol m^-2^ s^-1^) light-steps with duration of either 30 or 300 s (FL30, FL300). For the FL30 light regime, samples were taken 25 s after switching from low to high light (FL30_25s_HL). For the FL300 light regime, samples were taken at four different time points: 25 s after switching from low to high light (FL300_25s_HL), 290 s after light switches from low to high light phase (FL300_290s_HL), 25 s after switching from high to low light (FL300_325s_LL), and 290 s after switching from high to low light (FL300_590s_LL). Plants were acclimated to 25 °C, 15 °C, or 7 °C for at least 2h prior to sampling. Values are means ± SEM (n = 5-6 biological replicates). Ratios were analyzed using a two-way ANOVA to test the effects of temperature (Temp), sampling time (Time), and their interaction (TempxTime). Significant TempxTime interaction effects (indicated by inset p-values) were followed by Tukey testing. Different letters indicate statistical differences (*α*<0.05).

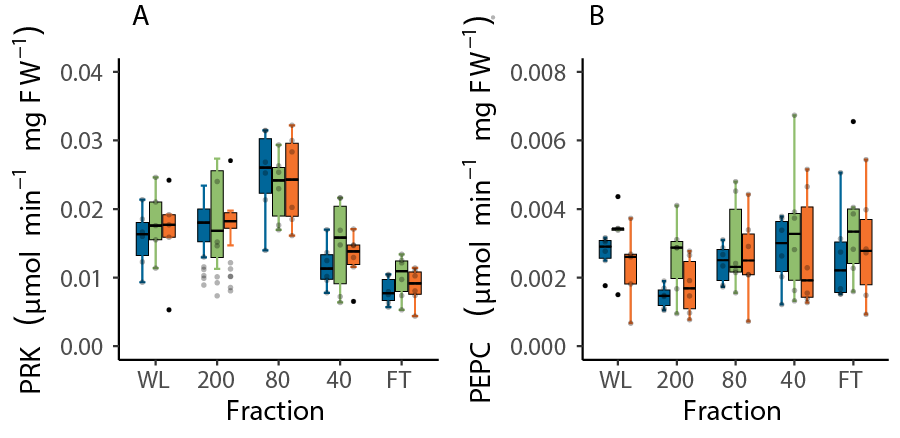

**Supplemental Figure S13.** Activities of phosphoribulokinase (PRK, A) and phosphoenolpyruvate kinase (PEPC, B) in whole leaf extract (WL) and in four leaf fractions obtained by filtering leaf material over liquid nitrogen sequentially through 200, 80, and 40 µm nylon meshes. FT is the flow-through collected from the final filtration at 40 µm. Maize plants were acclimated for one hour at the desired temperature (25, 15 or 7°C) with the lights in the cabinet turned on to a photosynthetic active radiation (*Q*) of 600 μmol m^-2^ s^-1^. Afterwards, the youngest completely expanded leaf was clamped into a 9 cm^2^ leaf chamber at 1500 µmol m^-2^ s^-1^ actinic red light, and samples were snap-frozen inside the gas exchange instrument after 30 min when steady state had been reached. n = 5-6 experimental replicates. Each replicate consisted of pooled samples from 11 plants.

.

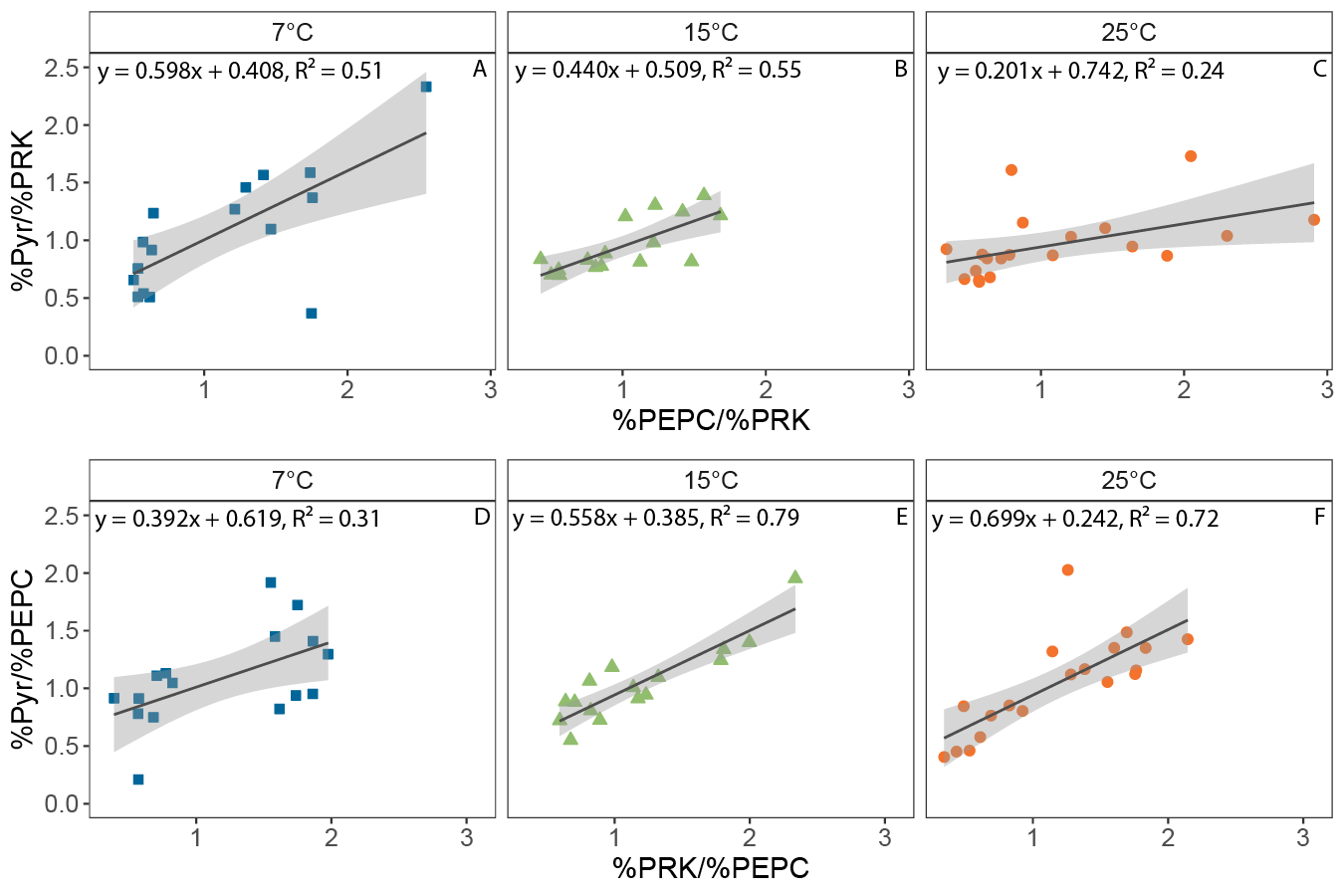

**Supplemental Figure S14.** Linear regression analysis of pyruvate concentration in four leaf fractions obtained by filtering leaf material over liquid nitrogen sequentially through 200, 80, and 40 µm nylon meshes. Pyruvate concentrations were normalized against fractional marker enzyme activities for mesophyll cells (phosphoenol-pyruvate carboxylase, PEPC) and bundle sheath cells (phosphoribulokinase, PRK), which were determined in parallel for each sample. Slopes and y-intercepts were used to determine proportional pyruvate abundance in each cell type, as summarized in Supplemental Table S2. Maize plants were acclimated for one hour at the desired temperature (25, 15 or 7°C) with the lights in the cabinet turned on to a photosynthetic active radiation (*Q*) of 600 μmol m^-2^ s^-1^. Afterwards, the youngest completely expanded leaf was clamped into a 9 cm^2^ leaf chamber at 1500 µmol m^-2^ s^-1^ actinic red light, and samples were snap-frozen inside the gas exchange instrument after 30 min when steady state had been reached. n = 5-6 experimental replicates. Each replicate consisted of pooled samples from 11 plants. Each symbol represents one of the four fractions from a fractionated pooled sample of 11 plants.

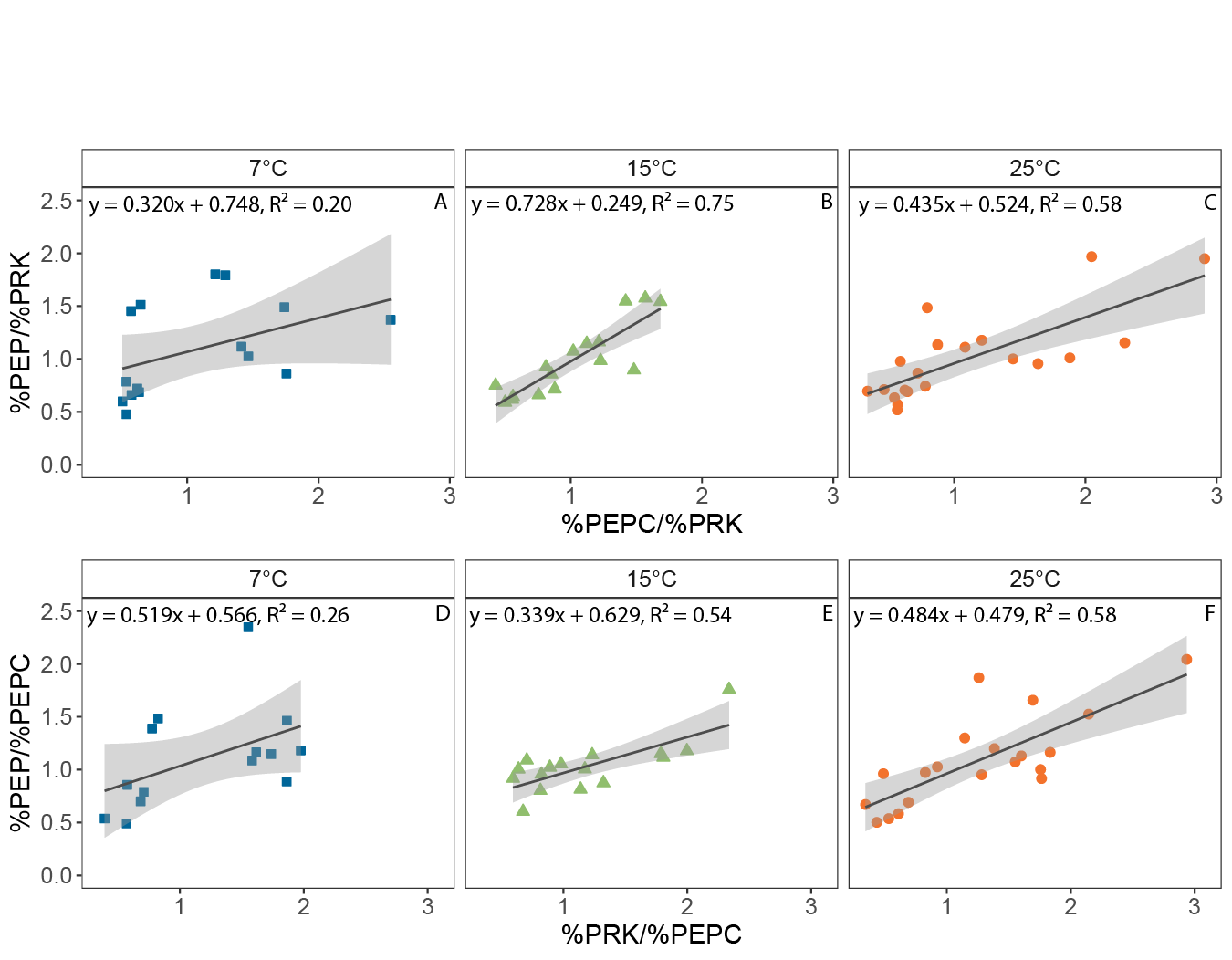

**Supplemental Figure S15.** Linear regression analysis of phosphoenol-pyruvate (PEP) concentration in four leaf fractions obtained by filtering leaf material over liquid nitrogen sequentially through 200, 80, and 40 µm nylon meshes. PEP concentrations were normalized against fractional marker enzyme activities for mesophyll cells (phosphoenol-pyruvate carboxylase, PEPC) and bundle sheath cells (phosphoribulokinase, PRK), which were determined in parallel for each sample. Slopes and y-intercepts were used to determine proportional PEP abundance in each cell type, as summarized in Supplemental Table S2. Maize plants were acclimated for one hour at the desired temperature (25, 15 or 7°C) with the lights in the cabinet turned on to a photosynthetic active radiation (*Q*) of 600 μmol m^-2^ s^-1^. Afterwards, the youngest completely expanded leaf was clamped into a 9 cm^2^ leaf chamber at 1500 µmol m^-2^ s^-1^ actinic red light, and samples were snap-frozen inside the gas exchange instrument after 30 min when steady state had been reached. n = 5-6 experimental replicates. Each replicate consisted of pooled samples from 11 plants. Each symbol represents one of the four fractions from a fractionated pooled sample of 11 plants.

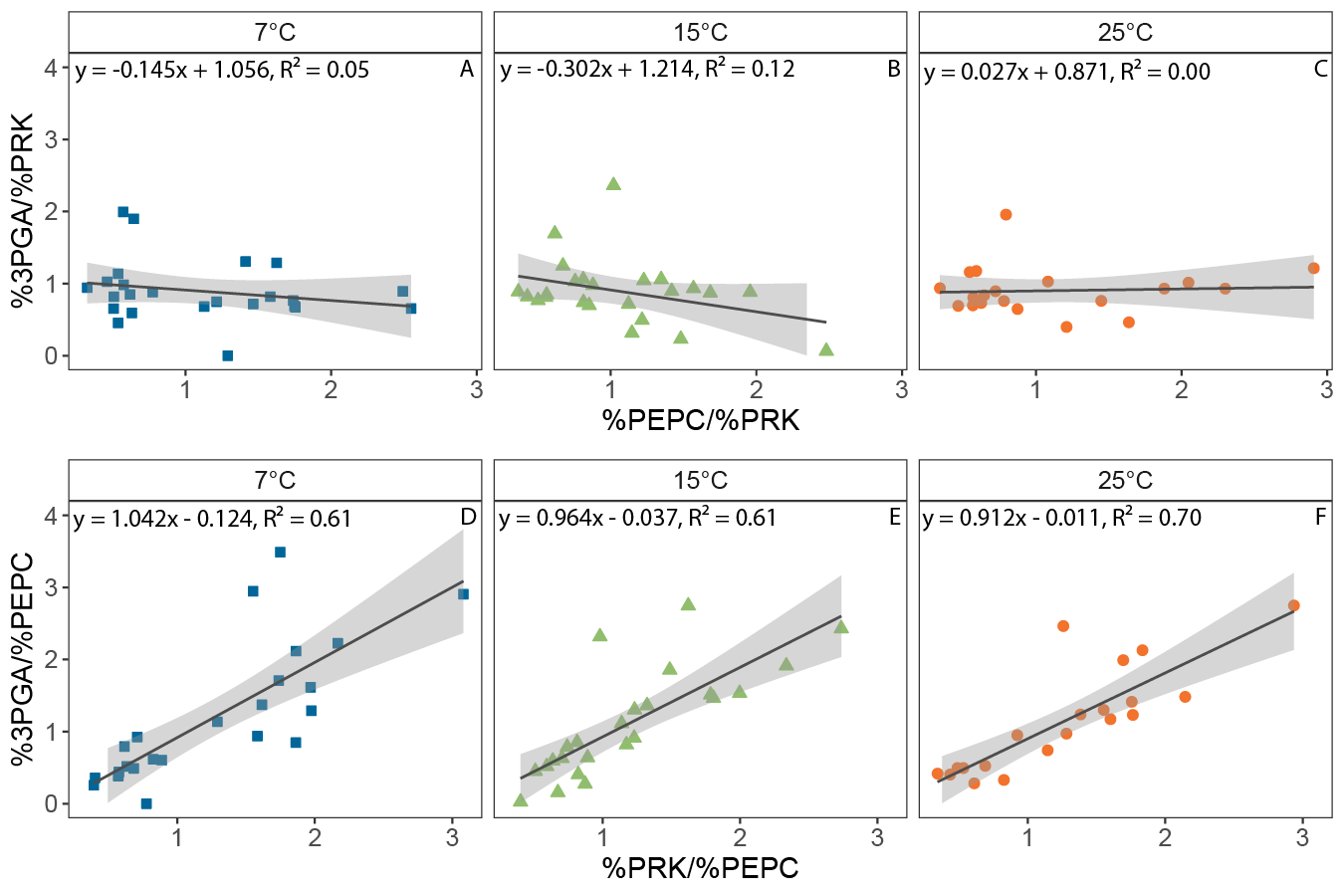

**Supplemental Figure S16.** Linear regression analysis of 3-phosphoglycerate (3PGA) concentration in four leaf fractions obtained by filtering leaf material over liquid nitrogen sequentially through 200, 80, and 40 µm nylon meshes. 3PGA concentrations were normalized against fractional marker enzyme activities for mesophyll cells (phosphoenol-pyruvate carboxylase, PEPC) and bundle sheath cells (phosphoribulokinase, PRK), which were determined in parallel for each sample. Slopes and y-intercepts were used to determine proportional 3PGA abundance in each cell type, as summarized in Supplemental Table S2. Maize plants were acclimated for one hour at the desired temperature (25, 15 or 7°C) with the lights in the cabinet turned on to a photosynthetic active radiation (*Q*) of 600 μmol m^-2^ s^-1^. Afterwards, the youngest completely expanded leaf was clamped into a 9 cm^2^ leaf chamber at 1500 µmol m^-2^ s^-1^ actinic red light, and samples were snap-frozen inside the gas exchange instrument after 30 min when steady state had been reached. n = 5-6 experimental replicates. Each replicate consisted of pooled samples from 11 plants. Each symbol represents one of the four fractions from a fractionated pooled sample of 11 plants.

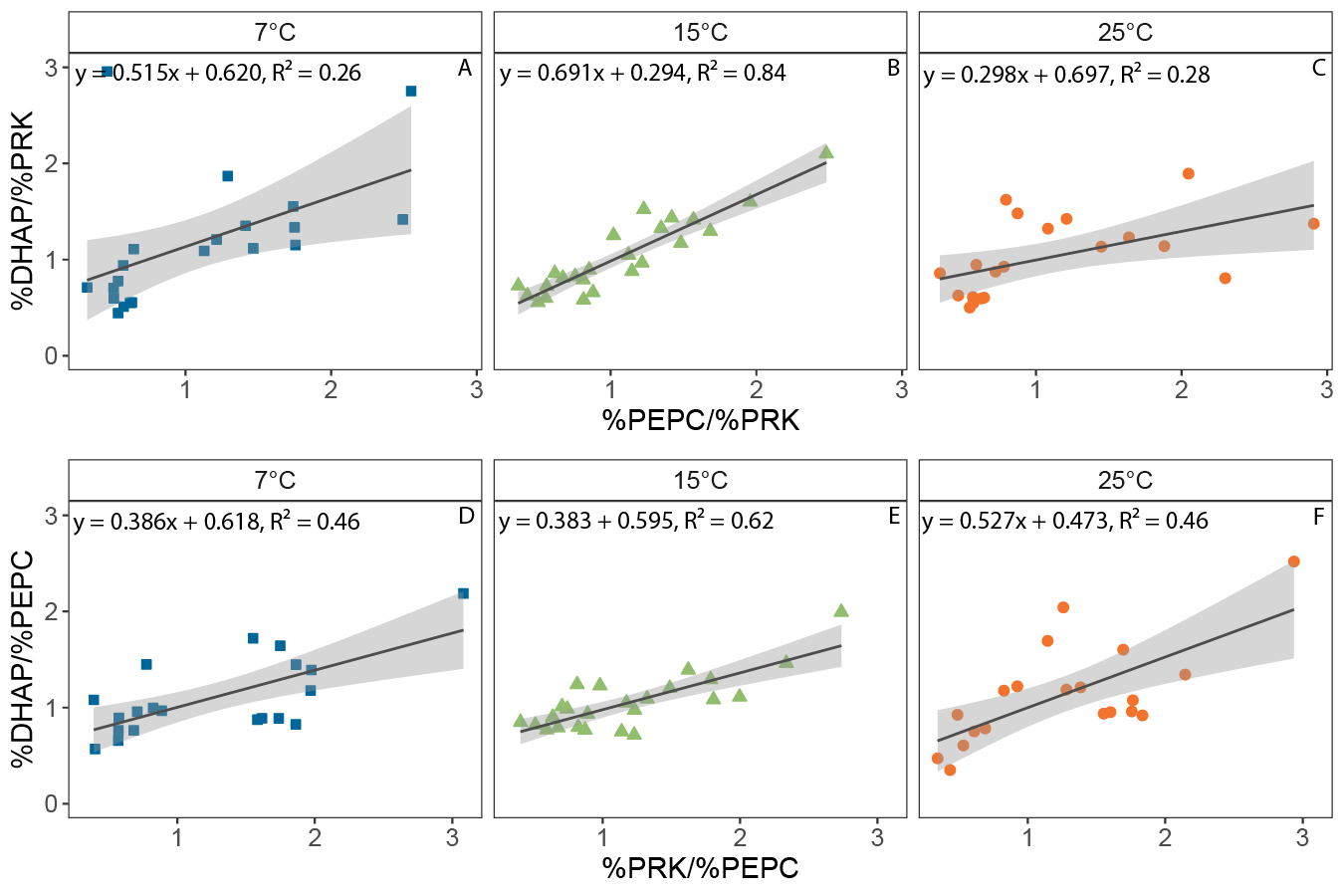

**Supplemental Figure S17.** Linear regression analysis of dihydroxyacetone-phosphate (DHAP) concentration in four leaf fractions obtained by filtering leaf material over liquid nitrogen sequentially through 200, 80, and 40 µm nylon meshes. DHAP concentrations were normalized against fractional marker enzyme activities for mesophyll cells (phosphoenol-pyruvate carboxylase, PEPC) and bundle sheath cells (phosphoribulokinase, PRK), which were determined in parallel for each sample. Slopes and y-intercepts were used to determine proportional DHAP abundance in each cell type, as summarized in Supplemental Table S2. Maize plants were acclimated for one hour at the desired temperature (25, 15 or 7°C) with the lights in the cabinet turned on to a photosynthetic active radiation (*Q*) of 600 μmol m^-2^ s^-1^. Afterwards, the youngest completely expanded leaf was clamped into a 9 cm^2^ leaf chamber at 1500 µmol m^-2^ s^-1^ actinic red light, and samples were snap-frozen inside the gas exchange instrument after 30 min when steady state had been reached. n = 5-6 experimental replicates. Each replicate consisted of pooled samples from 11 plants. Each symbol represents one of the four fractions from a fractionated pooled sample of 11 plants.

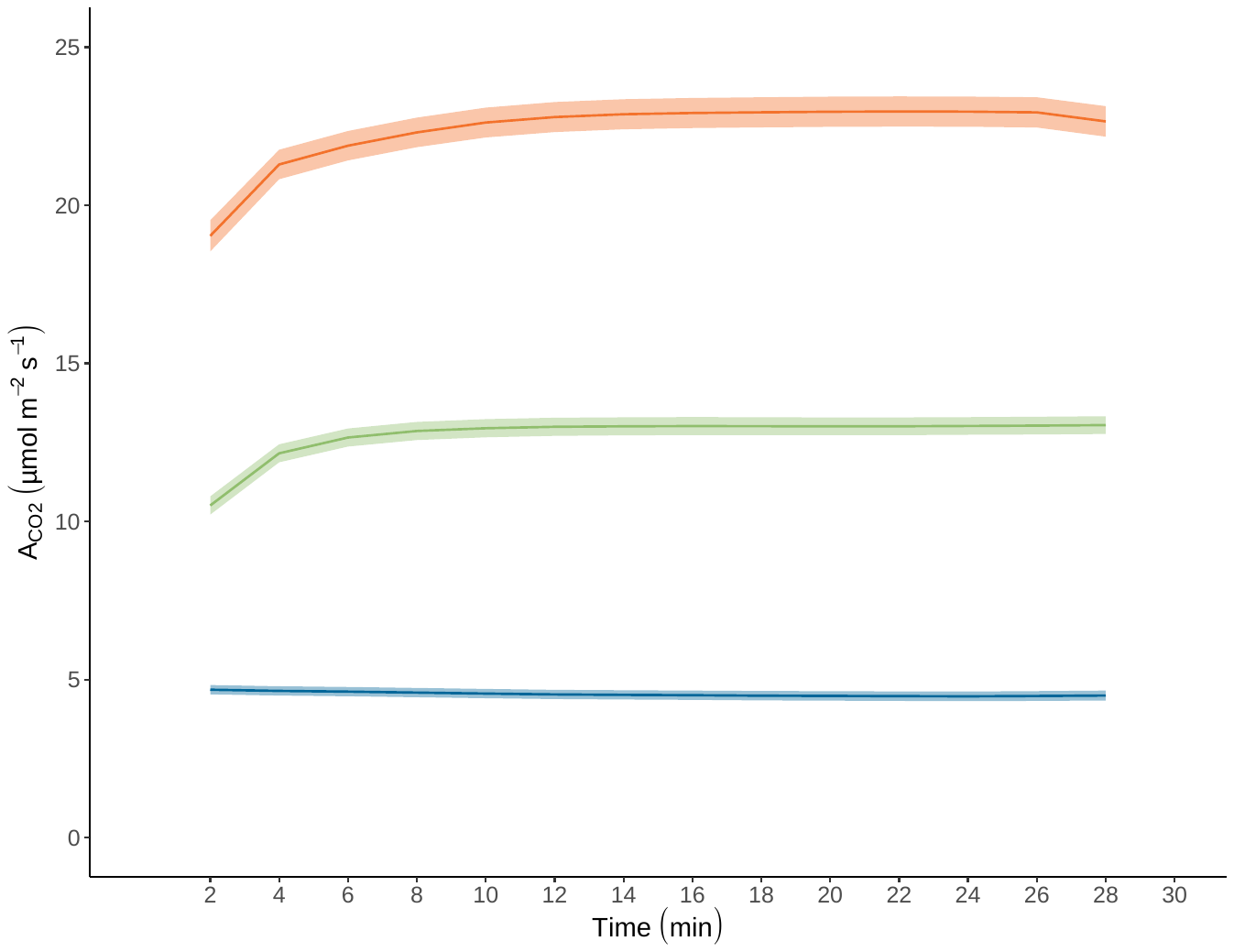

**Supplemental Figure S18.** Leaf CO_2_ assimilation (*A*_CO2_) logged before sampling for fractionation of leaf samples over liquid nitrogen sequentially through 200, 80, and 40 µm nylon meshes. Maize plants were acclimated for one hour at the desired temperature (25, 15 or 7°C) with the lights in the cabinet turned on to a photosynthetic active radiation (*Q*) of 600 μmol m^-2^ s^-1^. Afterwards, the youngest completely expanded leaf was clamped into a 9 cm^2^ leaf chamber at 1500 µmol m^-2^ s^-1^ actinic red light. Gas exchange data was logged every two seconds for 30 minutes, and samples were snap-frozen inside the gas exchange instrument after 30 min when steady state had been reached. Samples from 11 plants were pooled per experimental replicate for leaf fractionation.
